# An ALE meta-analytic review of top-down and bottom-up processing of music in the brain

**DOI:** 10.1101/2021.04.26.441417

**Authors:** Victor Pando-Naude, Agata Patyczek, Leonardo Bonetti, Peter Vuust

## Abstract

A remarkable feature of the human brain is its ability to integrate information from the environment with internally generated content. The integration of top-down and bottom-up processes during complex multi-modal human activities, however, is yet to be fully understood. Music provides an excellent model for understanding this since music listening leads to the urge to move, and music making entails both playing and listening at the same time (i.e. audio-motor coupling). Here, we conducted activation likelihood estimation (ALE) meta-analyses of 130 neuroimaging studies of music perception, production and imagery, with 2660 foci, 139 experiments, and 2516 participants. We found that music perception and production rely on auditory cortices and sensorimotor cortices, while music imagery recruits distinct parietal regions. This indicates that the brain requires different structures to process similar information which is made available either by an interaction with the environment (i.e. bottom-up) or by internally generated content (i.e. top-down).

## 1. Introduction

Music is a highly multifarious aspect of the human experience and therefore a privileged tool to gain insights into the key features of the human brain, its structure, and supporting neural processes. One of such key features is the ability to constantly integrate information and coordinate a variety of different tasks. Indeed, a successful existence depends on the brain’s capacity to combine external stimuli with internal representations to ultimately select the best response to the ever-changing environment^1^. The processes which regulate such needs have often been framed in the notorious “top-down/bottom-up” dichotomy. This simplistic approach has been redefined by the predictive coding theory (PC) which intends to set a framework that explains the complexity of the flow of information in the brain. Within this framework, music is an ideal example of the constant integration of multi-modal top-down and bottom-up information, as it includes different processes such as perception, imagery, and production/creation^2^. Despite decades of research in the field of music cognition, there is no comprehensive study investigating brain activation associated to these three distinct modalities of music processing.

Throughout the years, key examples of both top-down and bottom-up processes have been revealed. On the one hand, it has been shown that environmental stimuli activate different brain cortices, coherently with the chosen sensory modality^3^ (e.g. visual inputs activate visual brain regions, while auditory inputs activate auditory cortices). According to the PC, bottom-up processing is represented as a series of loops formed by ascending and descending connections between neighbouring levels in a hierarchically organized system (feedforward-feedback). This system is reliable and fast, with a semi-hardwired architecture sufficiently stable to allow for rapid processing of stimuli, and that can be dynamically re-shaped. On the other hand, top-down mechanisms have been described in diverse manners including anatomical (connections between levels of hierarchy), cognitive (hypothesis-driven), *gestaltist* (modulation of bottom-up), and dynamic (entrainment of neuronal populations by oscillatory activity)^4^; all suggesting that the system allows individuals to predict future events and stimuli. Notably, the term top-down should always be challenged when used^5^. According to the PC, top-down processes can be seen as influences that allow the semi-hardwired network to be flexible and ensure efficient and reliable sensory processing. In other words, the system takes as much new information as possible (perception) to construct predictions about the environment to rapidly apply it in a behaviour (action). Taking this in consideration, it is clear that bottom-up and top-down mechanisms are not opposites, and that ascending and descending connections are involved in both processes.

In this context, music is an ideal domain to study the constant integration of multi-modal mechanisms^6^, as music engages audio-motor coupling and other cognitive and emotional mechanisms such as pleasure^7^. Thus, how the brain processes music may represent a privileged tool to gain insights into such central topics. Music is built upon several concurrent bottom-up processes such as perception of sounds and rhythms. Moreover, it also comprises a number of top-down mechanisms such as music production and music imagery, the latter being the abstract mental manipulation of musical sounds^8^. On the neural level, previous studies investigating such processes have provided a wide array of underlying active brain areas, spanning from auditory and sensorimotor cortices^9^ to cerebellum^2^, cingulum^10^, basal ganglia^11^, hippocampus^12^, and amygdala^13^. In light of this, music may offer an ideal opportunity to investigate similarities and differences between top-down and bottom-up processes in complex multi-modal brain functioning.

A large number of high-quality studies on music neuroscience already exists. However, individual studies do carry the limitation of small sample size and therefore consequent low reliability^14^. Recent advancements in statistical methods and meta-analyses allow us to overcome restrictions related to data protection regulation^15^, gaining insights from a large population size and embrace “big data” human imaging. Currently, there are a number of coordinate-based algorithms for meta-analysis of neuroimaging studies such as the Activation Likelihood Estimation (ALE)^16^ and kernel density analysis (KDA)^17^. The ALE method reports location probabilities associated with each foci (set of x, y, and z coordinates), whereas KDA shows the number of foci surrounding a specific voxel.

In the field of music cognition, the ALE methodology has been used to investigate several independent features of music cognition including recruitment of motor areas^2^, auditory-motor entrainment^18^, hierarchical auditory organisation^19^, and music-evoked emotions^20^. Further, projects following open science principles such as the BrainMap^21^ platform and its GingerAle^14^ meta-analysis algorithm enable sharing of data and so accelerate progress in human brain mapping^22^. These factors converge to make the BrainMap ALE meta-analysis a convenient and appropriate method to compare brain activation across studies of music cognition.

Thus, for the first time, we have performed coordinate-based meta-analyses (CBMA) of a wide range of functional magnetic resonance imaging (fMRI) studies using state-of-the-art methods such as ALE, aiming to assess the neural mechanisms underlying music cognition. As a secondary aim, we sought to assess the broad nature of musical processing as a model of cognition itself. Specifically, appraising perception, production, and imagery in light of top-down and bottom-up processes and the recent theory of predictive coding. In this way, we explore the common and distinct patterns of brain activation during music perception, production, and imagery; and discuss how the brain activity associated with the three modalities may reflect the top-down and bottom-up nature of the brain.

## 2. Results

A total of 1707 articles were identified through database searching, and after removing 497 duplicates, 1210 articles were initially screened by title and abstract. After excluding 907 records, 303 articles were assessed for eligibility in the full-text screening stage. From these, 130 fulfilled criteria and were included in both qualitative and quantitative synthesis (ALE meta-analysis) (**Supplementary Figure 1**).

### 2.1. Characteristics of studies

The characteristics of all studies included in the final qualitative and quantitative synthesis of the meta-analysis are shown in **Table 1**. In music perception, a total of 105 studies and 2,035 participants (122 female; age = 26.6±6.7 years) was identified. Musicians were reported in 12% of the included studies, non-musicians in 69% and both in 13%. Neuroimaging method comprised of both fMRI (96%) and PET (4%). Musical features varied across studies including emotion (24%), melody (11%), harmony (9%), timbre (4%), tonality (4%), memory (3%), pitch (3%), structure (3%), rhythm (2%), tension (2%), creativity (2%), while 15% of studies did not specified the musical feature. Most of the auditory stimuli included unfamiliar stimuli (76%).

**Table 1.**
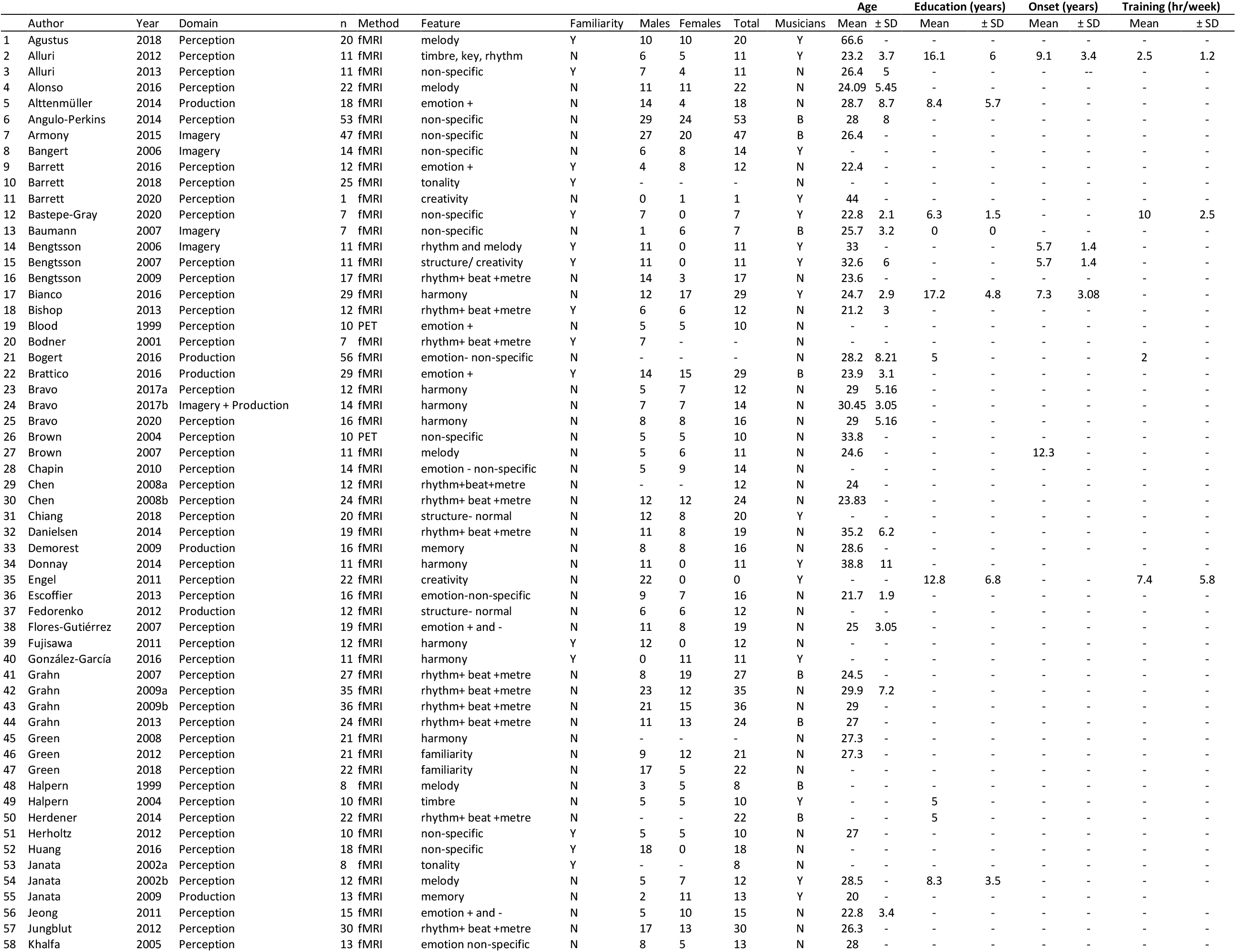
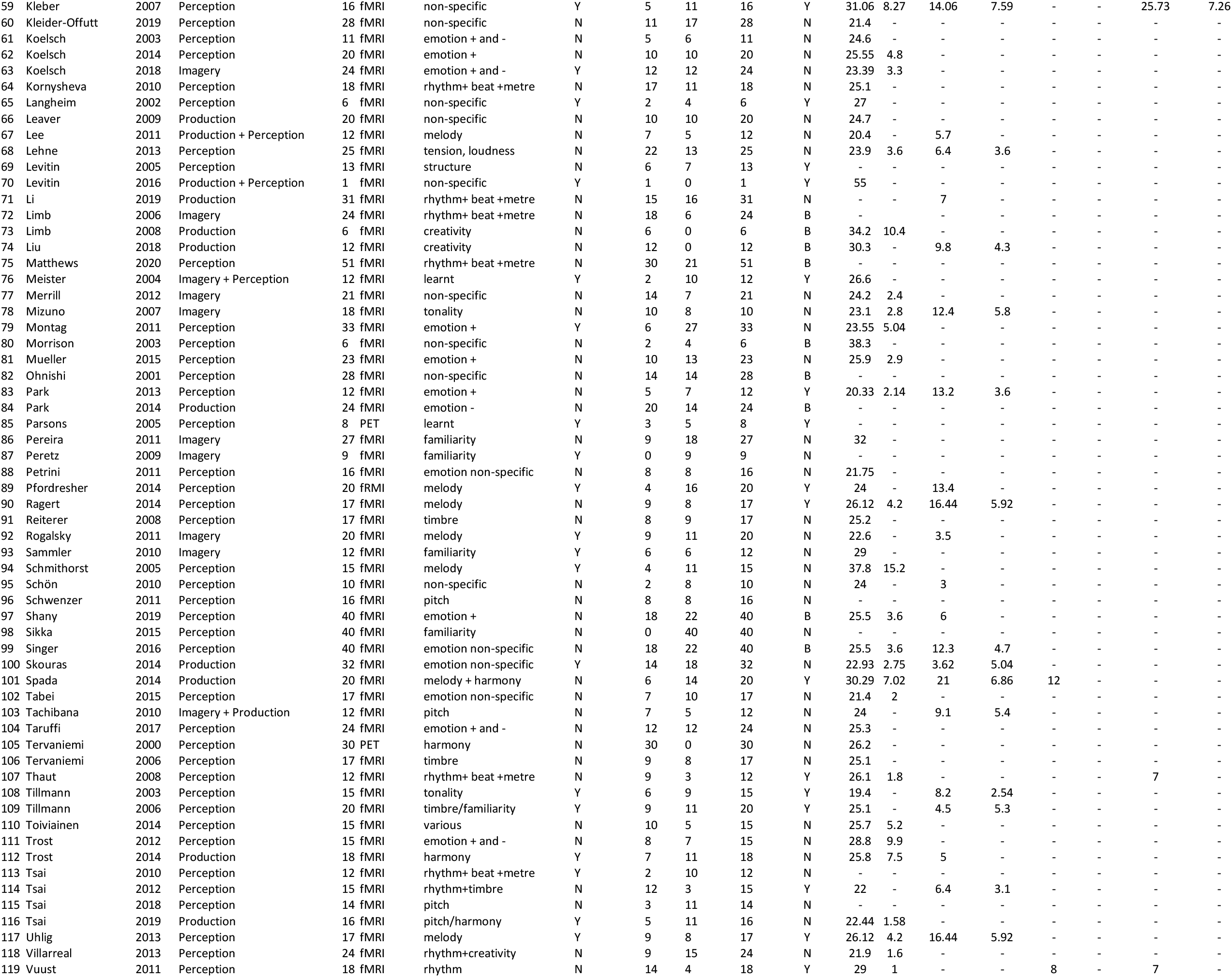
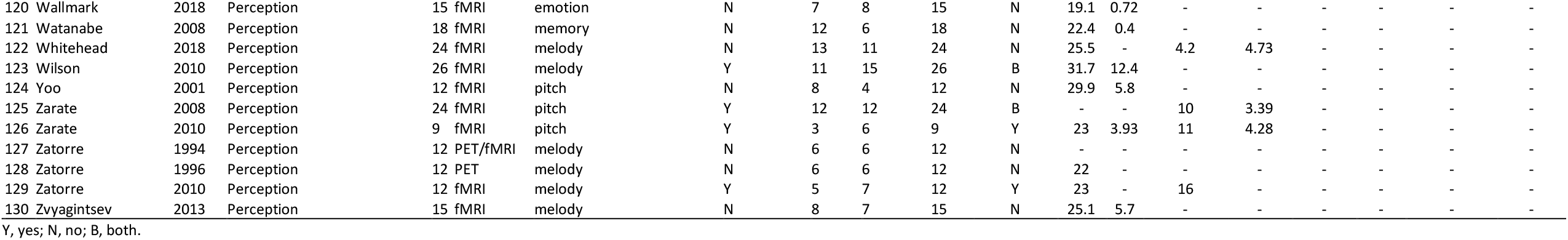
Characteristics of the studies included in meta-analysis.

In music production, a total of 19 studies and 292 participants (122 female; age= 29.8±6.1 years) was identified. Musicians were reported in 63% of the included studies, non-musicians in 16%, and both in 21%. Neuroimaging method comprised of both fMRI (95%) and PET (5%). Musical features varied across studies including harmony (11%), melody (5%), pitch (10%) and rhythm, metre and/or beat (11%). Overall, most of the studied focused on learnt abstracts (68%) compared to creative improvisation (47%). Most of the tasks included unfamiliar content (53%).

In music imagery, a total of 15 studies and 198 participants (108 female; age= 25.5±3 years) was identified. Musicians were reported in 40% of the included studies, non-musicians in 47%, and both in 13%. Neuroimaging method comprised of both fMRI (93%) and PET (7%). Musical features varied across studies including melody (27%), pitch (13%), timbre (7%) and non-specific (53%) of which most included unfamiliar imagery (60%).

### 2.2. MRI Quality

MRI quality of the included studies in the meta-analysis was assessed by a set of guidelines for the standardized reporting of MRI studies^23,24^. All studies reported their MRI design, software package and image acquisition, processing and analyses. Overall, good MRI practices were performed in the included studies (**Supplementary Table 1** and **Supplementary Table 2**). Neuroimaging data was acquired in either 1.5 T (27%), 3 T (66%), while 7% of the studies did not specified the magnetic field strength. Most of the studies used the Siemens MRI scanner (50%), others include General Electric (28%), Phillips Electronics (4%), Hitachi (1%), Brucker (4%), Magnex Eclipse (1%), CTI (1%), and few failed to report the scanner (2%). Most of the structural images were acquired using T1-weighted sequence (42%), magnetization-prepared rapid acquisition with gradient echo sequence (MPRAGE) (27%), spoiled gradient recalled acquisition in steady state (SPGR) (9%) with 1mm3-voxel size in 39% of the studies. T2-functional images were acquired mainly using echo-planar imaging (EPI) sequence (86%). Statistical analysis was conducted in either SPM (60%), AFNI (6%), FSL (7%), LIPSIA (5%) BrainVoyager (6%), MATLAB (2%), while 13% of studies did not specify the analysis software.

### 2.3. Primary outcomes: ALE meta-analyses of music perception, production, and imagery

#### 2.3.1. Music perception

The music perception ALE meta-analysis included 1898 foci, 105 experiments and 2035 subjects. Significant peak clusters resulted in the following areas: (1) right superior temporal gyrus (x=52, y=-20, z=4) extending over insula and inferior frontal gyrus; (2) left superior frontal gyrus (x=-54, y=-16, z=2) extending over transverse temporal gyrus, the insula and precentral gyrus; (3) left medial frontal gyrus (x= -2, y=-2, z=66) including the superior frontal gyrus and the right medial frontal gyrus; (4) right lentiform nucleus (putamen) (x=22, y=8, z=6) and the caudate body; (5) left lentiform nucleus (putamen) (x= -22, y= 4, z= 6) and the caudate body; (6) left cerebellum (lobule III) (x=-28, y=-64, z= -26); (7) left insula (x=-32, y=18, z=10) including the inferior frontal gyrus and (8) right precentral gyrus (x=54, y=0, z=46). Such results show consistency of cortical and subcortical areas presumably organized in a hierarchical manner processing both bottom-up and top-down musical information (**Figure 1, Table 2**).

**Figure 1.**
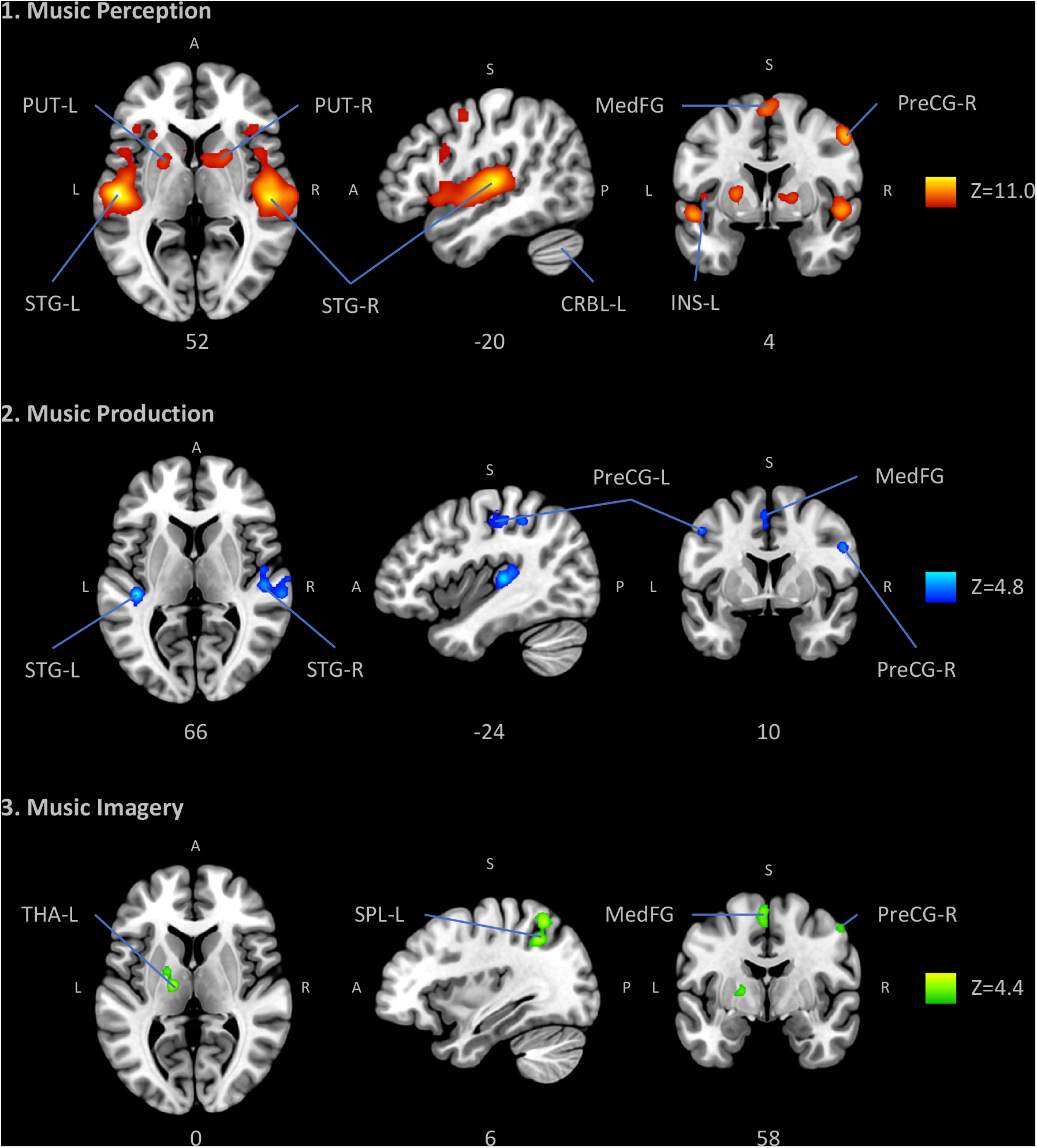
Anatomic likelihood estimation meta-analytic results for studies of music perception, production and imagery, at cluster level inference p < 0.05 (FWE). The primary outcomes included ALE meta-analyses of music perception, music production, and music imagery, independently. ROIs: CRBL, cerebellum; INS, insula; MedFG, medial frontal gyrus; PreCG, precentral gyrus (primary motor cortex or M1); PUT, putamen; SPL, superior parietal lobule; STG, superior temporal gyrus (primary auditory cortex); THA, thalamus; L, left; R, right; Z, peak Z-value. Figure created with Mango (http://rii.uthscsa.edu/mango//userguide.html)

**Table 2.**
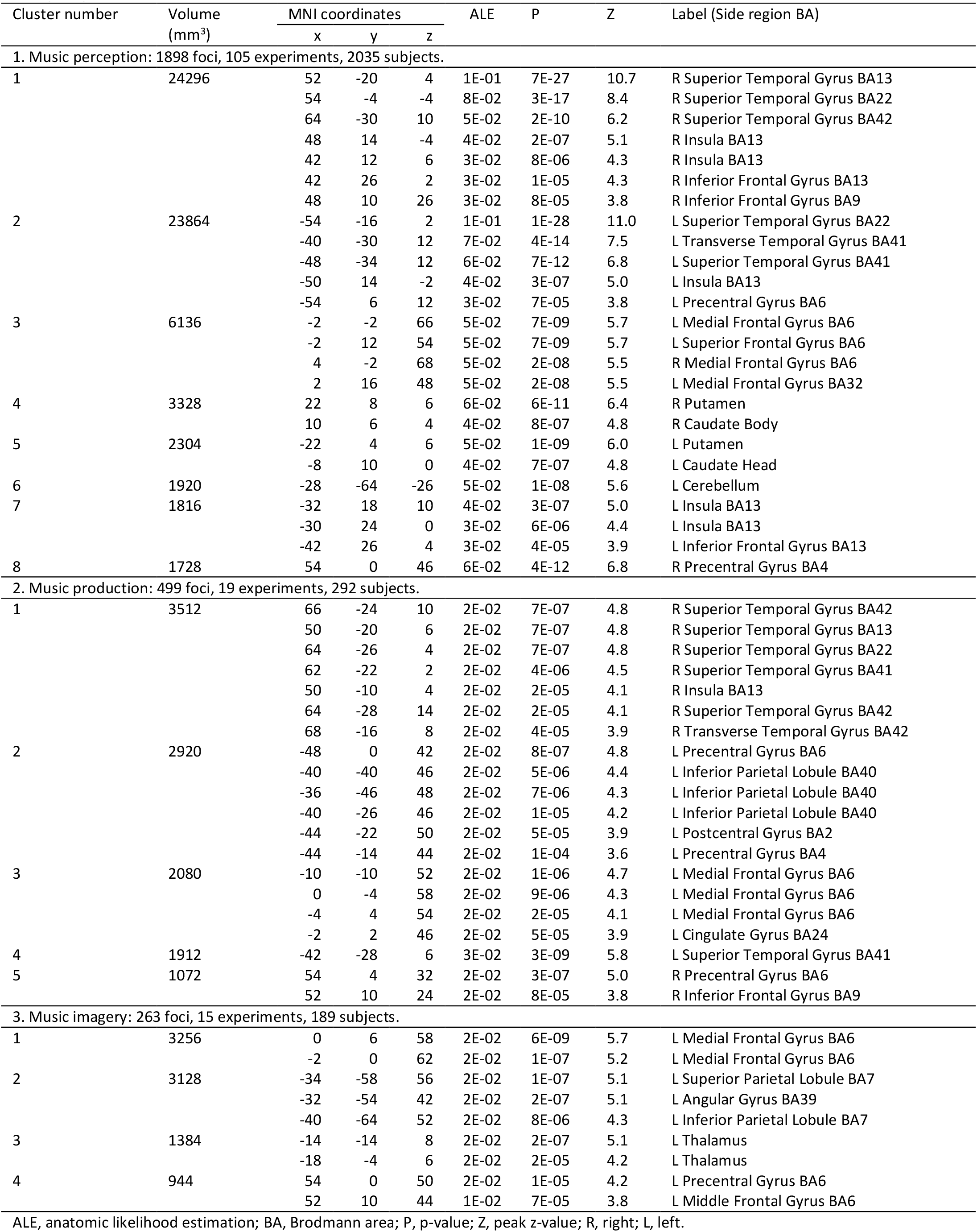
Anatomic likelihood estimation meta-analytic results for music perception, production and imagery at cluster level inference p < 0.05 (FWE).

#### 2.3.2. Music production

The music production ALE meta-analysis included 499 foci, 19 experiments and 292 subjects. Significant peak clusters resulted in the following areas: (1) right superior temporal gyrus (x=66, y=24, z=10) extending over Wernicke’s area, anterior and posterior transverse temporal area; (2) left precentral gyrus (x=-48, y=0, z=42) extending over precentral gyrus, inferior parietal lobule and post central gyrus; (3) left medial frontal gyrus (x=-10, y-10, z=52) including cingulate gyrus; (4) left superior temporal gyrus (x=-42, y=28, z=6); (5) right precentral gyrus (x=54, y= 4, z=32) extending over the inferior frontal gyrus. These results evidence not only the obvious involvement of the somatomotor system, but also limbic and executive cognitive functions while playing music. Interestingly, similar areas were found in music perception, further evidencing the integration of incoming information with high-order functions and motor behaviour (**Figure 1, Table 2**).

#### 2.3.3. Music imagery

The music imagery ALE meta-analysis included 263 foci, 15 experiments and 189 subjects. Significant peak clusters resulted in the following areas: (1) left medial frontal gyrus (x=0, y=6, z=58); (2) left superior parietal lobule (x=-34, y=-58, z=56) extending over angular gyrus and inferior parietal lobule; (3) left thalamus (x=-14, y=-14, z=8); (5) left precentral gyrus (x=52, y=10, z=4) encompassing the middle frontal gyrus. These results show that music imagination recruits motor areas involved in the generation of movements such as the premotor cortex and supplementary motor area; areas of the basal ganglia which are also involved in facilitation of movements; and parietal areas involved in perceptual-motor coordination and theory-of-mind (**Figure 1, Table 2**).

### 2.4. Contrast analyses

A subsequent contrast ALE meta-analysis was conducted to compare the primary outcomes, by means of conjunction and subtraction. The purpose of this analysis was to identify common and distinct brain activity between perception, production and imagery modalities of music cognition (**Supplementary Table 3**).

#### 2.4.1. Music perception vs music production

Clusters identified as likely to be active during both music perception and production (conjunction) include the right superior temporal gyrus (BA42), bilateral superior temporal gyrus (BA13), right superior temporal gyrus (BA22), left superior temporal gyrus (BA41), insula, right transverse temporal gyrus (BA42), left medial frontal gyrus (BA6) and inferior frontal gyrus (BA9). The contrast analysis (subtraction) yielded areas specific to perception (perception-production) and production (production-perception). Areas more likely to be active in music perception are the superior temporal gyrus (BA21) and superior temporal gyrus (BA22). Conversely, areas more likely to be active in music production include bilateral post central gyrus (BA2, BA40), bilateral precentral gyrus (BA4, BA6), bilateral superior, middle and medial frontal gyrus (BA6), left cingulate gyrus (BA24, BA32), left superior temporal gyrus (BA41, BA13, 22), left inferior parietal lobule (BA40), left insula (BA13), cerebellum (culmen) and the lentiform nucleus.

#### 2.4.2. Music production vs music imagery

Clusters that are presumably active during both music production and music imagery (conjunction) includes the left medial frontal gyrus (BA6). The contrast analysis (subtraction) resulted in areas either specific to music production (production-imagery) or to music imagery (imagery-production). Areas more likely to be active in music production include the left superior temporal gyrus (BA13, BA41, BA22), left transverse temporal gyrus (BA41), left postcentral gyrus (BA40) and the culmen. Areas more likely to be active while imagining music include the left superior parietal lobule (BA7), angular gyrus (BA39), medial frontal gyrus (BA6) and the precentral gyrus (BA6).

#### 2.4.3. Music imagery vs music perception

Clusters likely active during both music imagery and music perception (conjunction) include the left medial frontal gyrus (BA6), right precentral gyrus (BA6) and the globus pallidus. The contrast analysis (subtraction) resulted in areas either specific to music imagery (imagery-perception) or to music perception (perception-imagery). Areas more related to music imagery include left inferior parietal lobule (BA7, BA40), right middle and left medial frontal gyrus (BA6), right precentral gyrus (BA4) and the ventral lateral and ventral posterior lateral thalamus. Finally, areas more likely to be active during music perception include the bilateral superior temporal gyrus (BA22), left transverse temporal gyrus (BA41), right claustrum, left insula (BA13) and left caudate head.

### 2.5. Meta-analytic connectivity modelling (MACM)

MACM was performed to functionally segregate the behavioural contribution and the patterns of co-activation of each music-related region-of-interest (ROI) resulted from the primary outcomes (n=17). The ROIs were imported into the BrainMap database separately, to identify studies reporting activation within each ROI boundary (**Supplementary Table 4**). The functional characterization of each ROI is detailed in **Supplementary Table 5** and include the behavioural domains of action, perception, emotion, cognition and interoception.

#### 2.5.1. Music perception MACM

The right superior temporal gyrus ROI (1a) showed co-activation with left superior temporal gyrus, right claustrum, left medial frontal gyrus, right precentral gyrus, left insula, and left cerebellum. Relevant behavioural domains within its boundaries include execution, language, music, and auditory perception; and experimental paradigms including emotion induction, finger tapping, music comprehension, music production, and oddball, phonological, pitch, semantic, and tone discrimination.

The left superior temporal gyrus ROI (1b) showed co-activation with right superior temporal gyrus and right insula. Relevant behavioural domains within its boundaries include execution, language, music, positive emotion, and auditory perception; and experimental paradigms including emotion induction, finger tapping, music comprehension, music production, and oddball, phonological, pitch, semantic, and tone discrimination.

The left medial frontal gyrus ROI (1c) showed co-activation with left precentral gyrus, bilateral thalamus, bilateral cerebellum, bilateral superior temporal gyrus, and left superior parietal lobule. Relevant behavioural domains within its boundaries include execution, attention, music, and auditory perception; and experimental paradigms including emotion induction, finger tapping, music comprehension, music production, and pitch, semantic, tactile, and tone discrimination.

The right putamen ROI (1d) showed co-activation with left putamen, left medial frontal gyrus, and right inferior parietal lobule. Relevant behavioural domains within its boundaries include execution, attention, language, working memory, music, reasoning, reward, and auditory perception; and experimental paradigms including encoding, finger tapping, music comprehension, music production, reward, and pitch, semantic, tactile, and tone discrimination.

The left putamen ROI (1e) showed co-activation with right putamen, left medial frontal gyrus, bilateral superior temporal gyrus, and left cerebellum. Relevant behavioural domains within its boundaries include execution, attention, language, working memory, music, reward, and auditory perception; and experimental paradigms including emotion induction, finger tapping, music comprehension, and phonological, pitch, semantic, and tone discrimination.

The left cerebellum ROI (1f) showed co-activation with right cerebellum, bilateral insula, left medial frontal gyrus, bilateral superior temporal gyrus, left precentral gyrus, and right thalamus. Relevant behavioural domains within its boundaries include execution, attention, language, music, and auditory perception; and experimental paradigms including emotion induction, music comprehension, music production, and pitch, semantic, tactile, and tone discrimination.

The left insula ROI (1g) showed co-activation with right insula, left medial frontal gyrus, right inferior frontal gyrus, left inferior parietal lobule, left precentral gyrus, and right middle frontal gyrus. Relevant behavioural domains within its boundaries include attention, language, working memory, music, reasoning, reward, and auditory perception; and experimental paradigms including emotion induction, finger tapping, music comprehension, reward, and oddball, phonological, semantic, and tone discrimination.

The right precentral gyrus ROI (1h) showed co-activation with left precentral gyrus, left medial frontal gyrus, bilateral superior temporal gyrus, bilateral putamen, and right thalamus. Relevant behavioural domains within its boundaries include execution, attention, language, music, and auditory perception; and experimental paradigms including emotion induction, finger tapping, music comprehension, music production, reward, and pitch, semantic, and tone discrimination.

#### 2.5.2. Music production MACM

The right superior temporal gyrus ROI (2a) showed co-activation with left superior temporal gyrus, left medial frontal gyrus, left insula, left thalamus, and right precentral gyrus. Relevant behavioural domains within its boundaries include execution, attention, language, music, and auditory perception; and experimental paradigms including emotion induction, finger tapping, music comprehension, music production, reward, and oddball, phonological, pitch, semantic, and tone discrimination.

The left precentral gyrus ROI (2b) showed co-activation with right precentral gyrus, left medial frontal gyrus, left inferior parietal lobule, left middle temporal gyrus, left fusiform gyrus, right superior parietal lobule, and right cerebellum. Relevant behavioural domains within its boundaries include execution, attention, language, working memory, music, reasoning, and visual motor perception; and experimental paradigms including finger tapping, music comprehension, music production, and phonological, pitch, semantic, and tactile discrimination.

The left medial frontal gyrus ROI (2c) showed co-activation with left precentral gyrus, right cerebellum, left postcentral gyrus, and left claustrum. Relevant behavioural domains within its boundaries include execution, attention, and reward; and experimental paradigms including finger tapping, music comprehension, music production, reward, semantic discrimination, and tactile discrimination.

The left superior temporal gyrus ROI (2d) showed co-activation with right superior temporal gyrus, and left putamen. Relevant behavioural domains within its boundaries include execution, attention, language, music, social cognition, and auditory perception; and experimental paradigms including emotion induction, finger tapping, music comprehension, music production, and oddball, phonological, pitch, semantic, and tone discrimination.

The right precentral gyrus ROI (2e) showed co-activation with left precentral gyrus, anterior cingulate cortex, left cerebellum, bilateral superior parietal lobule, bilateral inferior parietal lobule, right insula, and right thalamus. Relevant behavioural domains within its boundaries include execution, language, reasoning, music, and social cognition; and experimental paradigms including emotion induction, finger tapping, music comprehension, music production, phonological discrimination, and pitch discrimination.

#### 2.5.3. Music imagery MACM

The left medial frontal gyrus ROI (3a) showed co-activation with bilateral insula, left superior parietal lobule, right inferior parietal lobule, and bilateral cerebellum. Relevant behavioural domains within its boundaries include execution, attention, language, memory, music, reasoning, and auditory perception; and experimental paradigms including emotion induction, finger tapping, music comprehension, music production, reward, and phonological, pitch, semantic, syntactic, tactile, and tone discrimination.

The superior parietal lobule ROI (3b) showed co-activation with right superior parietal lobule, left anterior cingulate cortex, right inferior frontal gyrus, and bilateral insula. Relevant behavioural domains within its boundaries include execution, attention, language, working memory, music, reasoning, reward, and auditory perception; and experimental paradigms including emotion induction, finger tapping, music comprehension, reward, and pitch, semantic, and tactile discrimination.

The left thalamus ROI (3c) showed co-activation with right thalamus, anterior cingulate cortex, left precentral gyrus, left inferior frontal gyrus, right insula, and right cerebellum. Relevant behavioural domains within its boundaries include execution, attention, working memory, music, reward, auditory perception; and experimental paradigms including emotion induction, finger tapping, motor learning, music comprehension, music production, reward, and oddball, phonological, tactile, and tone discrimination.

d. The right precentral gyrus ROI (3d) showed co-activation with left precentral gyrus, left medial frontal gyrus, right insula, bilateral putamen, and bilateral superior temporal gyrus. Relevant behavioural domains within its boundaries include execution, music, negative emotion, and auditory perception; and experimental paradigms including emotion induction, finger tapping, music comprehension, music production, and phonological, pitch, semantic, tactile, and tone discrimination.

## 3. Discussion

In the present study, we have conducted ALE meta-analyses of neuroimaging studies with a total of 2660 foci, 139 experiments, and 2516 subjects, to realise a comprehensive picture of the top-down and bottom-up mechanisms required by music processing. Our main results, whose robustness is guaranteed by the large numbers of studies included in the analyses, provide a complex and appetising picture of the brain activity underlying music processing. We show that music perception and music production rely on similar brain activation involving auditory cortices and sensorimotor cortices. In turn, music imagery presents a particular and unique activation of parietal and motor regions. Finally, our primary outcomes were complemented with contrast analyses and meta-analytic connectivity modelling, describing in full the complexity of the brain areas underlying music processing.

### 3.1. Characteristics of included studies

The publications included in this comprehensive systematic review and meta-analysis reported a clear research question, inclusion and exclusion criteria for participants, description of methods and explicit results. In line with reproducibility efforts, the included studies used state-of-the-art techniques and computational MRI tools important for the support of standardization of neuroimaging studies. However, some works lacked important demographic data such as the years of education, age of musical training onset, and current time of musical practice, which may have an effect on behavioural tasks and neuroimaging data.

### 3.2. Music perception

#### Temporal lobe areas

As expected, the main result concerning music perception involved temporal regions and especially auditory cortices. Indeed, we found convergence in various temporal lobe areas assumed to be related to auditory feature processing as well as semantic and conceptual aspects of music. Namely, primary auditory regions of Heschl’s gyrus (BA41, BA42), as well as secondary auditory area, namely Wernicke’s area (BA22). Mirrored by previous studies, these areas have been seen crucial for processing of auditory information including pattern recognition^25^ and syntax processing^26^. Hence, activation of both primary and secondary areas may reflect a hierarchical computation of auditory processing including lower-level extraction followed by alignment to preconstructed predictions in association areas. Our MACM results show that the superior temporal gyrus where the primary auditory cortex lies, co-activates with motor and pre-motor areas, insula, and cerebellum, supporting the idea that a large network is established while listening to music involving audio-motor coupling and limbic and paralimbic processing of emotional content.

#### Audio-motor coupling

We also found convergence in motor-associated areas, assumingly related to temporal processing, inherent audio-motor coupling and entrainment. These include areas within the basal ganglia (caudate, putamen, globus pallidus and thalamus), primary motor areas (precentral gyrus BA4), supplementary motor areas (premotor cortex, supplementary motor area BA6), and the cerebellum (lobule III). As music unfolds over time and does so at a specific periodicity (rhythm, beat, metre), it is not surprising to see brain activation in areas related to temporal processing in the motor domain^27^. Furthermore, previous studies have proposed significant connectivity within the fronto-temporal-cerebellar network during the perceptual analysis of music listening^28,29^. In concordance with neuroimaging results, electroencephalography (EEG) studies have found synchronised neural activity in the gamma-band within this network, in response to rhythmic expectancy^30,31^. Additionally, specific neuronal firing patterns were found in those areas with respect to time comprehension^32^. These include ramping spike activity, which refers to a consistent increase or decrease in neuronal firing rate, spectral responses and network-produced ensemble patterns^33^. In relation to the lentiform nucleus, the putamen has been proposed to be important for internal beat perception^34^ and for generation of internalised groupings or chunking of action representations^35^. MACM revealed an extensive network that can represent audio-motor coupling with areas such as primary auditory and motor cortices, premotor cortex, basal ganglia, insula, and cerebellum.

#### Broad activation of the insular cortex

Further, our ALE analysis outlined an arguably broad activation of the insular cortex (BA13) extending under the superior frontal gyrus, sub-lobar insular and superior temporal gyrus. The insula cortex, folded deep within the lateral sulcus and covered by the frontoparietal and temporal operculum^36^, has been primarily divided into two parts: the anterior and posterior insula^37^. However, more than a dozen subdivisions have been identified^38^. With rich connectivity with both cortical and subcortical structures, the insular cortex is seen as an anatomical integration hub with rich connectivity and heavy cross-modal input^39^. Within our MACM analysis, this is particularly evident with the functionally connectivity of the insula spreading across both cortical and subcortical structures, likely reflecting the breadth of its function as a multimodal integration center. As categorised by Kurth and colleagues^40^ in a meta-analysis of close to 1, 800 experiments, the insular cortex serves a number of categorically distinct functions including sensorimotor, olfacto-gustatory, socio-emotional and cognitive functions. With respect to music perception, such rich functional diversity aligns with previous findings presenting the role of the insular cortex in auditory processing including allocation of auditory attention^41^, time perception and interceptive salience^42^ as well as music evoked emotion and changes in physiological arousal^43^. With these in mind, a broad activation of the insular cortex in our results appraises the complexity and multiplicity of music as a stimulus to the human ear.

### 3.3. Music production

#### Motor areas

Playing an instrument or engaging in music production requires extremely intricate motor control functions as well as the ability to internalise and portray emotional meaning. Accordingly, and unsurprisingly, with regards to our music production analysis, we found convergence in areas related to motor planning and execution. Namely, areas within the precentral gyrus, (BA4), middle frontal gyrus (BA6) and the supramarginal gyrus (BA40). Aside from their obvious motor function in relation to instrument playing, these areas are also likely to be involved in various sensory-motor coupling. Previous studies have repeatedly outlined the PMC and the planum temporale to be the crucial players in various sensorimotor transformations^44^. Activation within the somatosensory association area such as the supramarginal gyrus has been previously linked to tactile interpretation and limb location^45^ crucial for use of an instrument. In relation to music, the rostral dPMC has been recruited during increasingly more complex rhythms^46^ and saliency^47^. This means that the PMC could be responsible for modifying motor sequences to reflect auditory sequential patterns in a predictive and anticipatory manner.

#### Limbic areas

Aside from motor dexterity crucial for the technical component of instrument playing, the performance must engage the listener with loaded emotional content. Although with reduced breadth of activation compared to motor areas, our results point to a number of regions involved in the processing and conveyance of emotional material including the anterior cingulate gyrus (BA24) and insula (BA13). Previous findings of neuroanatomical studies have outlined the temporal lobe having strong connectivity with the anterior cingulate gyrus^48^. Further, for an appropriate emotional response the anterior cingulate has shown to require specific sensory information^44^. Thus, the anterior cingulate gyrus may be crucial for top-down control of emotional meaning and bottom-up processing of emotional content. In complement, numerous neuroimaging studies have outlined the activation of cingulate gyrus during musical performance^49,50^. Interestingly, an emerging body of research is suggesting a role of the cerebellum in cognition and emotion with reciprocal connections to various limbic system regions. The fastigial nucleus in particular has been shown to have extensive topographic connections with non-motor systems including the limbic system encompassing areas such as the anterior cingulate gyrus^51^.

### 3.4. Music imagery

#### Frontal regions

*While imagining music*, our results point to a convergence in motor-associated areas within the medial frontal gyrus (supplementary motor area or SMA), precentral gyrus, putamen and thalamus. As previously mentioned, neuroimaging studies on rhythm, metre and beat have outlined the medial frontal gyrus, superior frontal gyrus along with the basal ganglia in the context of temporal feature processing and inherent audio-motor coupling^52^. In relation to imagery, activation within the SMA is particularly interesting, being repeatedly found in various imagery domains including visual imagery^53^. Furthermore, this region is also found in relation to imagination in hallucination studies^54^. Within our results, the SMA was specific to imagery suggesting its distinct role in mental manipulation and generation of internalised representations also during music.

#### Parietal regions

The main result from our ALE meta-analysis show music imagery to recruit parietal regions including the superior and inferior parietal lobule as well as the angular gyrus. With regards to the parietal lobules (BA7), this result is far from surprising. These areas have been previously discussed in the context of auditory attention^55^ and working memory^56^. Specific to imagery, previous research on motor imagery has reported the parietal lobule as a key region in several documentations using rTMS^57^ and fMRI^58^. Furthermore, activation within the angular gyrus (BA39) compliments the aforementioned parietal activation. Previous findings have suggested its role in memory retrieval^59^, working memory^60^, and perspective taking^61^. Further, our MACAM analysis specific to the parietal areas have revealed a distinct connection pattern with a sequence of areas including frontoparietal and motor regions. Together, their joint role could be interpreted as key in initiating and formulating the experience of imagery involving appropriate cognitive abilities.

Importantly, what must be noted is that despite several authors describing imagery to recruit secondary auditory areas, we found no temporal area activation as it did not survive the correction for multiple comparisons. However, such temporal areas were only found in the uncorrected analysis together with other limbic and paralimbic areas. This may be due to variability of designs in the imagery studies, individual differences in imagery vivacity or a low number of studies.

### 3.5. Comparison between music perception, production, and imagery

A direct comparison of music perception, production and imagery for each of the respective pairs revealed several indications of music processing specificity likely resulting from a bidirectional organisation of the brain. Music imagery showed a unique activation of left superior and inferior parietal lobule (BA7). Previously, studies outlined these areas to be key for top-down or goal directed attentional orienting^62^, and specific to imagery, the superior parietal lobule has been named a potential candidate for top-down generation and maintenance of domain-unspecific mental images^63^. Crucially, both music perception and production, when contrasted with music imagery, showed similar broad activation of temporal and frontal areas associated with auditory processing and motor planning and execution. This is particularly interesting as to show that both perception and production must involve both top-down and bottom-up processes, thus, supporting the embodied and predictive approach which appreciates the human motor system and its actions as a reciprocal part of perception and cognition itself^64^.

### 3.6. Limitations and future perspectives

Although our meta-analyses comprise a large number of studies and thus arguably provide reliable results, some limitations should be mentioned. First, by definition meta-analyses rely on the combination of the data from previous studies and therefore are highly dependent on the quality of such studies. In our case, we utilised studies that possessed high-quality standards, making feasible data extraction and conduction of statistical analysis. However, we should highlight that a thorough description of the methods and an even stronger focus on the description of the experimental settings would be highly beneficial for the field and thus we encourage future studies to further increase their already high-level quality. Second, we relied on the assumptions of the GingerALE method that was used in our study. Therefore, the accuracy of our findings necessarily relies on the statistical estimation of coordinate-based anatomic foci (input) which were treated as spatial probability distributions centred at the given coordinates. As conceivable, we could not control the heterogeneity of the methods used in the studies included in the meta-analysis (ranging from pre-processing software, statistical thresholds, smoothing, and participants’ characteristics) and we acknowledge that this fact could represent a potential confounder for our analyses.

As in traditional meta-analyses, we have assessed the publication bias^65^ and revealed that our results are robust as the clusters obtained from the primary outcomes presented FSN values within lower and upper boundaries. Importantly, this outcome proves a robust convergence of the foci emerged from our analyses even when the total amount of studies was small, particularly in the music production and imagery CBMA’s (details in **Supplementary Table 6**).

With regard to future perspectives, we believe that the neuroscientific community would benefit from designing new ad-hoc studies to directly investigate the relationship between brain activity and different features of music processing such as music perception, production and imagination. Although our results provide a robust picture of the brain networks recruited by those different processes, carefully designed experiments which systematically modulate and compare stimuli and conditions would further refine the knowledge on this broad and complex topic.

## 4. Conclusions

In this comprehensive review, we have conducted a series of neuroimaging meta-analyses on decades of studies investigating the brain activity underlying music processing. We have shed new light on how the brain processes and integrates top-down and bottom-up mechanisms during complex multi-modal human activities such as music. The outcomes of our synthesis highlight three main concurrent music procedures, known as music perception (both top-down and bottom-up), music production (both top-down and bottom-up) and imagery (top-down). Our results show that the brain relies on different structures and mechanisms to process similar musical information. Indeed, music perception and music production depend on auditory cortices, sensorimotor cortices and cerebellum. Differently, music imagery shows a key recruitment of parietal regions. Taken together, our findings provide robust evidence that the brain requires different structures to process similar information which is made available either by the interaction with the environment (i.e. bottom-up) or by internally generated content (i.e. top-down).

## 5. Methods and Materials

### 5.1. Search strategy, screening, and extraction

This comprehensive systematic review and meta-analysis followed procedures from the Cochrane Handbook for Systematic Reviews^66^, and from the Center for Reviews and Dissemination (https://www.york.ac.uk/crd/). The review protocol was pre-registered in PROSPERO (CRD42019140177), and was carried out in accordance with the PRISMA statement^67^. A systematic literature search was conducted in PubMed, Scopus and PsycInfo for articles published up to February 26^th^, 2021. The search strategy for music perception, production and imagery was developed using keywords and MeSH terms (details in **Supplementary Information** p.2).

To ensure concurrence between the research question and data extracted, the following inclusion criteria was established: (1) healthy adults (18-80 age range); (2) studies investigating music listening, music playing, or music imagery, with no restrictions on music genre, parameter, or instrument; (3) using fMRI or PET; (4) whole-brain analyses; and (5) results reported in stereotactic coordinates either Talairach or Montreal Neurological Institute (MNI) three-dimensional-coordinate system. Studies were excluded using the following criteria: (1) review articles with no original experimental data, (2) populations with any form of disorder or disability; (3) paediatric population; (4) auditory stimuli not related to music; (5) functional connectivity analyses; (6) region-of-interest (ROI) analyses.

Two reviewers (AP and VPN) independently screened by title and abstract and selected articles for full-text review and data extraction. Screening and data extraction were performed using the Covidence tool^68^. Any disagreements that arose between the reviewers were resolved through discussion.

The following variables were extracted from each study: first author, year of publication, number of participants, age, sex, musical feature, year of education, year of musical training, age of onset of musical training, training hours per week, and MRI acquisition, processing, and analysis parameters. The main outcome to extract was coordinates resulting from functional brain activity related to music perception, production and/or imagination. If any of these points were not reported in the original article, authors were contacted to retrieve this information. Nine authors were contacted, with 3 positive answers.

### 5.2. Quality assessment of MRI studies

A set of guidelines used for the standardization of reporting MRI results was utilized to assess the quality of the included studies^23,24^. Such guidelines dictate a more consistent and coherent policy for the reporting of MRI methods to ensure that methods can be understood and replicated.

### 5.3. Activation Likelihood Estimation (ALE) and meta-analytic connectivity modelling (MACM)

All meta-analyses were performed using the activation likelihood estimation (ALE) method, implemented in GingerALE software v3.0.2^69^, from the BrainMap^69^ platform. Reported significant coordinates (foci) were extracted from each study. If necessary, coordinates were converted from Talairach coordinates to MNI space using the Lancaster transform (icbm2tal) incorporated in GingerALE^14,16^. The ALE method extracts the coordinates from the included studies and tests for anatomical consistency and concordance between the studies. The coordinates are weighted according to the size of the sample (number of subjects), and these weightings contribute to form estimates of anatomic likelihood estimation for each intracerebral voxel on a standardized map. This method treats anatomic foci not as single points, but as spatial probability distributions centred at the given coordinates. Thus, the algorithm tests the correlation between the spatial locations of the foci across MRI studies investigating the same construct and assesses them against a null-distribution of random spatial association between experiments^46^. Statistical significance of the ALE scores was determined by a permutation test using cluster-level inference at p < 0.05 (FWE), with a cluster-forming threshold set at p < 0.001. The primary outcome was brain functional activity related to music cognition with the aim of comprehensively examine the brain regions associated with music perception, music production, and music imagery, independently.

Then, the resulting ALE maps from each group were tested for similarity (conjunction) and difference (subtraction) in a contrast analysis with the purpose of identifying common and distinct brain areas recruited while listening, playing and/or imagining music. The GingerALE software was used following procedures described in the manual^70^. In short, during the first step, the foci from the two groups of interest were merged into a single text file and an ALE meta-analysis was conducted on the new file. Then, three ALE maps are imported to the GingerALE algorithm which randomly divides the pooled foci (foci of two datasets to be compared) into two new datasets of the same size as the original dataset. Then, voxel-wise ALE scores are calculated by subtracting one ALE image from the other. This procedure is repeated 10,000 times to yield a higher number of permutations compared to that of original dataset^71^. Finally, the ‘true’ difference of ALE scores is established and compared to the differences under a null distribution. This yields a voxel-wise p value of the difference which was calculated using cluster-level inference at p < 0.05 (FWE), with a cluster-forming threshold set at p < 0.001.

Meta-analytic connectivity modelling (MACM) was conducted to identify patterns of co-activation from regions-of-interest (ROI) resulting from the primary outcomes, aiming to functionally segregate each region’s putative contribution to behavioural domains and paradigm classes^72,73^. Such analyses were performed using Sleuth^74^ to identify studies reporting activation within each music-related ROI boundary independently and included the experiment level search criteria of “context: normal mapping” and “activations: activation only”. Then, ALE meta-analyses were performed in GingerALE over all foci resulted after the search in Sleuth to identify regions of significant convergence. Statistical significance of the ALE scores was determined by a permutation test using cluster-level inference at p < 0.05 (FWE), with a cluster-forming threshold set at p < 0.001. Functional characterization of music-related clusters was based on the “Behavioral Domain” (BD) meta-data categories in the BrianMap database which include action, perception, emotion, cognition and interoception.

All meta-analytic results (ALE maps) were visualized using Mango^21^ on the MNI152 1mm standard brain, and resulting coordinates were cross-referenced to the Harvard-Oxford Cortical and Subcortical Atlas and the Juelich Histological Atlas via NeuroVault^75^ and FSLeyes^76^, respectively.

### 5.4. Fail-Safe N analysis (FSN)

Coordinate-based meta-analyses such as ALE can be subject to different forms of publication bias which may impact results and invalidate findings (e.g., the “file drawer problem”). The Fail-Safe N analysis (FSN)^65^ assesses the robustness of results against potential publication bias. This method refers to the amount of contra-evidence that can be added to a meta-analysis before the results change and can be obtained for each cluster that survives thresholding in an ALE meta-analysis. It is estimated that a 95% confidence interval for the number of studies that report no local maxima varies from 5 to 30 per 100 published studies of normal human brain mapping. Using the upper bound and the fact that the CBMA’s consist of 105 music perception studies, 19 music production studies, and 15 music imagery studies, an estimate for the number of unpublished experiments is 32, 6, and 5, respectively. Therefore, the minimum FSN was defined as 32 for music perception, 6 for music production, and 5 for music imagery. A higher FSN indicates more stable results and hence a higher robustness.

## Acknowledgments

The Center for Music in the Brain (MIB) is supported by The Danish National Research Foundation (grant number DNRF 117). The authors wish to thank Hella Kastbjerg for assistance with language check and proofreading.

## Authors’ contributions

VPN and AP designed the meta-analysis. VPN and AP conducted the screening of the studies and data extraction. LB performed the sensitivity analysis. VPN and AP wrote the first draft of the manuscript and prepared the initial versions of the figures and tables. LB edited and wrote paragraphs in the Introduction and Discussion. VPN prepared the final figures and tables. PV contributed to financially support the study and commented the final versions of the manuscript. VPN finalized the paper for submission and all authors edited the manuscript and approved its final version.

## Conflict of interest

There is no actual or potential financial and other conflict of interest related to this manuscript.

## Data availability

The data supporting the findings of this study is freely available at the Open Science Framework (OSF) website https://osf.io/mtrha/?view_only=c85a6244897b4a7f9f96c25f9149bbcf

## Notes

https://osf.io/mtrha/?view_only=c85a6244897b4a7f9f96c25f9149bbcf

### Competing Interest Statement

The authors have declared no competing interest.

