## Supplementary Information for "An ALE meta-analytic review of top-down and bottom-up processing of music in the brain"

**Search strings.**

Last search date: 26.02.2021

**PsycInfo**

( music imagery OR music listening OR music production OR music composition OR music performance OR music playing OR music perception OR auditory imagery ) AND magnetic resonance imaging

612 results

**Scopus**

(TITLE-ABS-KEY ( music imagery ) OR TITLE-ABS-KEY ( music listening ) OR TITLE-ABS-KEY ( music production ) OR TITLE-ABS-KEY ( music performance ) OR TITLE-ABS-KEY ( music playing ) OR TITLE-ABS-KEY ( music composition ) OR TITLE-ABS-KEY ( music perception ) ) AND TITLE-ABS-KEY ( magnetic resonance imaging )

963 results

**PubMed**

( “music imagery” [All Fields] OR “music listening” [All Fields] OR “music production” [All Fields] OR “music performance” [All Fields] OR “music playing” [All Fields] OR “music composition” [All Fields] OR “music perception” [All Fields]) AND (“magnetic resonance imaging” [MeSH Terms] OR “magnetic resonance imaging” [All Fields])

132 results

**Supplementary Figure 1. PRISMA flowchart for literature search process.**

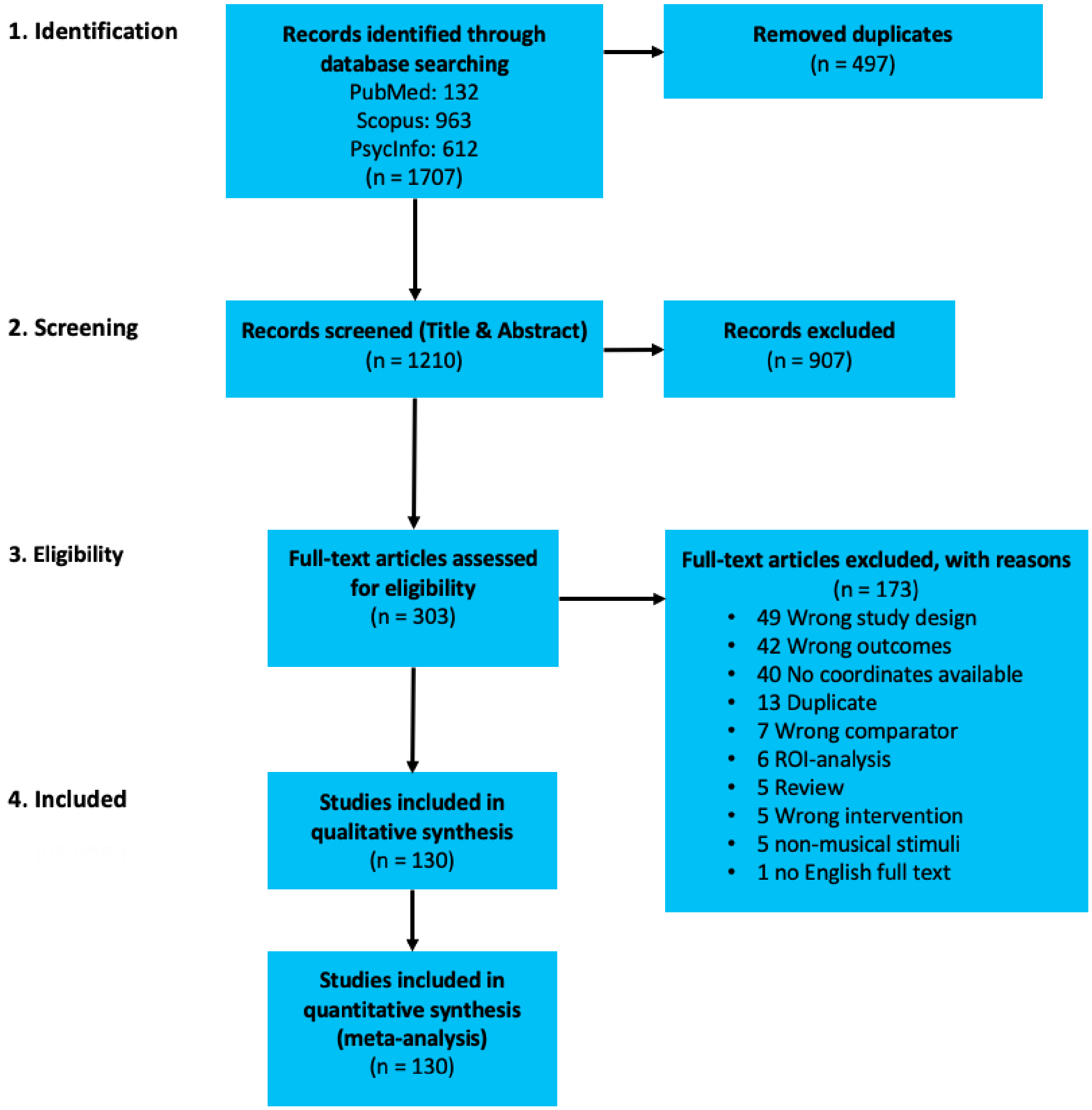

**Supplementary Figure 1**. PRISMA flowchart for literature search process.

**PRISMA Checklist**

SI = Supplementary Information

| Section/topic | # | Checklist item | Reported on page # |
|---|---|---|---|
| **TITLE** | | | |
| Title | 1 | Identify the report as a systematic review, meta-analysis, or both. | 1 |
| **ABSTRACT** | | | |
| Structured summary | 2 | Provide a structured summary including, as applicable: background; objectives; data sources; study eligibility criteria, participants, and interventions; study appraisal and synthesis methods; results; limitations; conclusions and implications of key findings; systematic review registration number. | 2 |
| **INTRODUCTION** | | | |
| Rationale | 3 | Describe the rationale for the review in the context of what is already known. | 3 |
| Objectives | 4 | Provide an explicit statement of questions being addressed with reference to participants, interventions, comparisons, outcomes, and study design (PICOS). | 4 |
| **METHODS** | | | |
| Protocol and registration | 5 | Indicate if a review protocol exists, if and where it can be accessed (e.g., Web address), and, if available, provide registration information including registration number. | 17 |
| Eligibility criteria | 6 | Specify study characteristics (e.g., PICOS, length of follow-up) and report characteristics (e.g., years considered, language, publication status) used as criteria for eligibility, giving rationale. | 17 |
| Information sources | 7 | Describe all information sources (e.g., databases with dates of coverage, contact with study authors to identify additional studies) in the search and date last searched. | 17 |
| Search | 8 | Present full electronic search strategy for at least one database, including any limits used, such that it could be repeated. | SI, p.2 |
| Study selection | 9 | State the process for selecting studies (i.e., screening, eligibility, included in systematic review, and, if applicable, included in the meta-analysis). | 17 |
| Data collection process | 10 | Describe method of data extraction from reports (e.g., piloted forms, independently, in duplicate) and any processes for obtaining and confirming data from investigators. | 17 |
| Data items | 11 | List and define all variables for which data were sought (e.g., PICOS, funding sources) and any assumptions and simplifications made. | 17 |
| Risk of bias in individual studies | 12 | Describe methods used for assessing risk of bias of individual studies (including specification of whether this was done at the study or outcome level), and how this information is to be used in any data synthesis. | 17 |
| Summary measures | 13 | State the principal summary measures (e.g., risk ratio, difference in means). | 18 |
| Synthesis of results | 14 | Describe the methods of handling data and combining results of studies, if done, including measures of consistency (e.g., $I^2$) for each meta-analysis. | 18 |

| Section/topic | # | Checklist item | Reported on page # |
| --- | --- | --- | --- |
| Risk of bias across studies | 15 | Specify any assessment of risk of bias that may affect the cumulative evidence (e.g., publication bias, selective reporting within studies). | 18 |
| Additional analyses | 16 | Describe methods of additional analyses (e.g., sensitivity or subgroup analyses, meta-regression), if done, indicating which were pre-specified. | 18 |
| **RESULTS** | | | |
| Study selection | 17 | Give numbers of studies screened, assessed for eligibility, and included in the review, with reasons for exclusions at each stage, ideally with a flow diagram. | 5 |
| Study characteristics | 18 | For each study, present characteristics for which data were extracted (e.g., study size, PICOS, follow-up period) and provide the citations. | 5 |
| Risk of bias within studies | 19 | Present data on risk of bias of each study and, if available, any outcome level assessment (see item 12). | 5 |
| Results of individual studies | 20 | For all outcomes considered (benefits or harms), present, for each study: (a) simple summary data for each intervention group (b) effect estimates and confidence intervals, ideally with a forest plot. | 6 |
| Synthesis of results | 21 | Present results of each meta-analysis done, including confidence intervals and measures of consistency. | 6-10 |
| Risk of bias across studies | 22 | Present results of any assessment of risk of bias across studies (see Item 15). | SI p.26 |
| Additional analysis | 23 | Give results of additional analyses, if done (e.g., sensitivity or subgroup analyses, meta-regression [see Item 16]). | 6-10 |
| **DISCUSSION** | | | |
| Summary of evidence | 24 | Summarize the main findings including the strength of evidence for each main outcome; consider their relevance to key groups (e.g., healthcare providers, users, and policy makers). | 11 |
| Limitations | 25 | Discuss limitations at study and outcome level (e.g., risk of bias), and at review-level (e.g., incomplete retrieval of identified research, reporting bias). | 14 |
| Conclusions | 26 | Provide a general interpretation of the results in the context of other evidence, and implications for future research. | 16 |
| **FUNDING** | | | |
| Funding | 27 | Describe sources of funding for the systematic review and other support (e.g., supply of data); role of funders for the systematic review. | 20 |

**Supplementary Table 1. Characteristics of MRI acquisition and analysis.**

| | | | MRI acquisition | | | | T1 | | | | T2 | | | | Analysis | |
|---|---|---|---|---|---|---|---|---|---|---|---|---|---|---|---|---|
| | Author | Year | Teslas | MRI-system | MRI-model | Head-coil | Sequence | TR (ms) | TE (ms) | Voxel size (mm) | Sequence | TR (ms) | TE (ms) | Voxel size (mm) | Software | Method |
| 1 | Agustus | 2018 | 3 | Siemens | Trio | 12-channel | T1-w | - | - | - | EPI | 11360 | 30 | 3x3x2 | SMP8 | MPM |
| 2 | Alluri | 2012 | 3 | GE | Signa | - | T1-w | - | - | - | EPI | 2000 | 32 | - | SPM8 | GLM, PCA |
| 3 | Alluri | 2013 | 3 | GE | Signa | standard | T1-w | 6.552 | 2.824 | 0.94x0.94x1.2 | EPI | 2200 | 30 | - | SPM8 | PCR |
| 4 | Alonso | 2016 | 3 | Siemens | Trio | - | MP2RAGE | 2300 | 4.18 | 1x1x1 | EPI | 2100 | 29 | 3x3x3 | SPM8 | GLM |
| 5 | Alttenmüller | 2014 | 3 | Siemens | Allegra | 8 | T1-w | 1550 | 7.3 | 1x1x1 | T2-w | 2000 | 30 | - | | GLM |
| 6 | Angulo-P | 2014 | 3 | GE | MR750 | 32 | T1-w | 2300 | 3 | 1x1x1 | EPI | 3000 | 40 | 2x2x3 | FSL | GLM |
| 7 | Armony | 2015 | 3 | GE | MR750 | 32 | T1-w | 2300 | 3 | 2x2x3 | EPI | 3000 | 40 | 2x2x3 | SPM8 | LOSO |
| 8 | Bangert | 2006 | 1.5 | GE | Signa | quadrature | SPGR | - | - | - | EPI | 4500 | 40 | - | SPM99 | GLM |
| 9 | Barrett | 2016 | 3 | Siemens | Trio | - | T1-w | 2500 | 4.82 | - | EPI | 2000 | 25 | 1x1x1 | SPM5 | GLM |
| 10 | Barrett | 2018 | 3 | Phillips | Achieva | 32 | MPRAGE | - | - | 0.7x0.7x0.7 | EPI | 2500 | 25 | - | SPM12 | Custom |
| 11 | Barrett | 2020 | 3 | Phillips | Achieva | 32 | MPRAGE | - | - | 1x1x1 | EPI | 2000 | 30 | - | SPM12 | GLM |
| 12 | Bastepe-G | 2020 | 1.5 | Siemens | Aera | 20 | MPRAGE | 1900 | 2.84 | - | EPI | 4000 | 50 | - | SPM12 | GLM |
| 13 | Baumann | 2007 | 3 | Phillips | Intera | body coil | T1-w | - | - | 0.98x0.98x0.75 | EPI | 2000 | 35 | 2.75x2.74x4.5 | Matlab | GLM |
| 14 | Bengtsson | 2006 | 1.5 | GE | Signa Horizon | - | T1-w | - | - | 1x1x1 | EPI | 4000 | 60 | - | SPM99 | GLM |
| 15 | Bengtsson | 2007 | 1.5 | GE | Signa Horizon | - | T1-w | - | - | 0.86x0.86x2 | EPI | 4000 | 60 | - | SPM99 | GLM |
| 16 | Bengtsson | 2009 | 3 | Siemens | Electra | - | T1-w | - | - | 1x1x1 | EPI | 3000 | 40 | - | SPM | - |
| 17 | Bianco | 2016 | 3 | Siemens | Trio | 32 | MP2RAGE | 5000 | 2.03 | 1x1x1 | EPI | 2000 | 30 | 2.3x2.3x2.3 | SPM8 | GLM |
| 18 | Bishop | 2013 | 3 | Siemens | Magnetom | - | MPRGE | 1830 | 4.43 | - | EPI | 3000 | 31 | 3x3x3 | SPM2 | - |
| 19 | Blood | 1999 | - | Siements | HR+ | - | - | - | - | - | - | - | - | - | - | GLM |
| 20 | Bodner | 2001 | 1.5 | Siemens | Magnetom | birdcage | - | 4000 | 4 | - | EPI | 3000 | 40 | - | APM99 | GLM |
| 21 | Bogert | 2016 | 3 | Siemens | Magnetom | 20 | T1-w | - | - | 1x1x1 | EPI | 2000 | 32 | 3x3x4 | SPM8 | GLM |
| 22 | Brattico | 2016 | 1.5 | GE | Signa | - | T1-w | - | - | - | EPI | 3000 | 32 | - | SPM8 | GLM |
| 23 | Bravo | 2017a | 3 | GE | Signa | - | T1-w | - | - | 1x1x1 | EPI | 3000 | 40 | - | SPM8 | PPI |
| 24 | Bravo | 2017b | 3 | GE | Signa | - | T1-w | - | - | 1x1x1 | EPI | 3000 | 40 | - | SPM8 | Regression |
| 25 | Bravo | 2020 | 3 | GE | Signa | - | T1-w | - | - | 1x1x1 | EPI | 3000 | 40 | - | SPM12 | - |
| 26 | Brown | 2004 | - | CTI | HR+ | - | - | - | - | - | - | - | - | - | - | - |
| 27 | Brown | 2007 | 2 | Elsinc | Gyrex | - | T1-w | - | - | 1x1x1 | EPI | 2000 | - | - | - | - |
| 28 | Chapin | 2010 | 3 | GE | Signa | - | SPGR | 325 | - | - | EPI | 1200 | 35 | 3.75x3.75x4 | AFNI | GLM |
| 29 | Chen | 2008a | 1.5 | Siemens | Sonata | - | T1-w | - | - | 1x1x1 | EPI | 1000 | 50 | 5x5x5 | AFNI | GLM |
| 30 | Chen | 2008b | 1.5 | Siemens | Sonata | - | T1-w | - | - | 1x1x1 | EPI | 10000 | 50 | 5x5x5 | AFNI | GLM |
| 31 | Chiang | 2018 | 3 | Siemens | Tim Trio | - | MPRAGE | 1900 | 2.26 | 1x1x1 | EPI | 3000 | 35 | 3x3x3.5 | FSL | GLM |
| 32 | Danielsen | 2014 | 1.5 | GE | Signa | - | FSPGR | - | - | - | EPI | 3000 | 40 | - | SPM8 | GLM |
| 33 | Demorest | 2009 | 1.5 | GE | Signa | custom | FSPGR | - | - | - | EPI | 3000 | 50 | - | FSL | ICA |
| 34 | Donnay | 2014 | 3 | Philips | - | quadrature | - | - | - | - | EPI | 2000 | 30 | - | - | - |
| 35 | Engel | 2011 | 3 | Brucker | Medspeck | birdcage | MPRAGE | 1300 | - | 1x1x1.5 | EPI | 1300 | 10 | - | SPM5 | GLM |
| 36 | Escoffier | 2013 | 3 | Siemens | Magentom | - | MPRAGE | 2530 | 1.64 | 1x1x1 | EPI | - | - | - | FSL | GLM |
| 37 | Fedorenko | 2012 | 3 | Siemens | Trio | 32 | T1-w | 2000 | 2.39 | 1.33x1.33x1.33 | EPI | 2000 | 30 | - | SPM5 | GLM |
| 38 | Flores-G | 2007 | 1.5 | GE | v9.x | quadrature | T1-w | - | - | - | EPI | 3000 | 60 | 4x4x8 | SPM2 | GLM |
| 39 | Fujisawa | 2011 | | Magnex | Marconi | | | | | | EPI | 6000 | 55 | 3x3x3 | SPM2 | Box-car |
| 40 | González-G | 2016 | 1.5 | Phillips | Achieva | - | T1-w | 10.2 | 4.2 | 1x1x1 | EPI | 1000 | 40 | 4x4x4 | SPM8 | GLM |
| 41 | Grahn | 2007 | 3 | Brucker | Medspec | - | SPGR | - | - | - | EPI | 1100 | 37.5 | - | SPM99 | GLM |
| 42 | Grahn | 2009a | 3 | Siemens | Trio | - | MRAGE | - | - | - | EPI | - | - | 1x1x1 | SPM5 | GLM |
| 43 | Grahn | 2009b | 3 | Siemens | Trio | - | MPRAGE | 2250 | 2.99 | 1x1x1 | EPI | 2190 | 30 | - | SPM5 | PPI |
| 44 | Grahn | 2013 | 3 | Siemens | Trio | - | MPRAGE | 2250 | 2.99 | 1x1x1 | EPI | 2190 | 30 | - | SPM5 | U |
| 45 | Green | 2008 | 1.5 | GE | Signa Excite | - | T1-w | - | - | - | EPI | 2700 | 40 | - | SPM5 | U |
| 46 | Green | 2012 | 1.5 | GE | Signa | - | T1-w | - | - | - | EPI | 2700 | 40 | - | SPM5 | - |
| 47 | Green | 2018 | 3 | Siemens | Magnetom | 32 | MPRAGE | - | - | - | EPI | 2000 | 24 | 3.4x3.4x3 | FSL | GLM |
| 48 | Halpern | 1999 | - | Siemens | Exact HR+ | - | - | - | - | - | - | - | - | - | - | - |
| 49 | Halpern | 2004 | 1.5 | Siemens | Vision | - | - | - | - | 1x1x1 | EPI | 1000 | | 5x5x5 | Matlab | U |
| 50 | Herdener | 2014 | - | - | - | - | - | - | - | - | - | - | - | - | BrainVoyager | GLM |
| 51 | Herholtz | 2012 | 3 | Siemens | Trio | 32 | T1-w | - | - | 1x1x1 | EPI | 2100 | 30 | 3.5x3.5x3.5 | FSL | GLM |
| 52 | Huang | 2016 | 3 | Siemens | Tim Trio | - | T1-w | 2530 | 2.43 | 1x1x1 | EPI | 2200 | 30 | - | SPM8 | GLM |
| 53 | Janata | 2002a | 3 | GE | Signa | - | T1-w | - | - | 2.6x2.6x5 | EPI | - | - | 2.6x2.6x5 | SPM99 | GLM |
| 54 | Janata | 2002b | 1.5 | GE | Horizon | birdcage | T1-w | 650 | 6.6 | - | EPI | 2000 | 35 | - | SPM99 | GLM |
| 55 | Janata | 2009 | 3 | Siemens | Trio | - | MPRAGE | 2000 | 4.82 | 1x1x1 | EPI | 2000 | 25 | - | SPM5 | GLM |

| | | | | | | | | | | | | | | | | |
|---|---|---|---|---|---|---|---|---|---|---|---|---|---|---|---|---|
| 56 | Jeong | 2011 | 3 | GE | Signa | 8 | SPGR | - | - | - | EPI | 3000 | 35 | - | SPM8 | GLM |
| 57 | Jungblut | 2012 | 3 | Siemens | Trio | - | - | - | - | - | EPI | 2200 | 30 | 3.44x3.44x3.74 | SPM8 | GLM |
| 58 | Khalfa | 2005 | 3 | Brucker | Medspec | - | T1-w | - | - | 1x0.75x1.22 | EPI | - | - | - | SPM99 | GLM |
| 59 | Kleber | 2007 | 1.5 | Siemens | Vision | - | MPRAGE | - | - | - | EPI | 3000 | 40 | - | SPM2 | GLM |
| 60 | Kleider-O | 2019 | 3 | Siemens | Tim Trio | 12 | MPRAGE | - | - | 1x1x1 | EPI | 2250 | 4.18 | 3.4x3.4x4 | SPM8 | TFCE |
| 61 | Koelsch | 2003 | 3 | Brucker | Medspec | - | T1-w | - | - | - | EPI | 3000 | - | - | LIPSIA | GLM |
| 62 | Koelsch | 2014 | 3 | Siemens | Magnetom | - | T1-w | - | - | 1x1x1 | EPI | 2000 | 30 | - | LIPSIA | GLM |
| 63 | Koelsch | 2018 | 3 | Siemens | Tim Trio | - | MPRAGE | - | - | 1x1x1 | EPI | 2000 | 30 | - | LIPSIA 2.1 | GLM |
| 64 | Kornysheva | 2010 | 3 | Siemens | Trio | - | T1-w | - | - | - | EPI | 2000 | 30 | - | LIPISIA | GLM |
| 65 | Langheim | 2002 | 1.5 | GE | Signa | quadrature | - | 3500 | 3.75 | - | Multi-slice | 2000 | 24 | 3.74x3.75x3.75 | SPM96 | GLM |
| 66 | Leaver | 2009 | 3 | Siemens | Trio | - | T1-w | - | - | 1x1x1 | EPI | 1000 | 30 | 3x3x3 | BrainVoyager | GLM |
| 67 | Lee | 2011 | 3 | Phillips | Achieva | - | MPRAGE | - | - | 1x1x1 | EPI | 2000 | 35 | 3x3x3 | SPM5 | U |
| 68 | Lehne | 2013 | 3 | Siemens | Tim Trio | - | MPRAGE | - | - | 1x1x1 | EPI | 2000 | 30 | - | SPM8 | GLM |
| 69 | Levitin | 2005 | 3 | GE | Signa | - | - | - | - | - | - | - | - | - | - | - |
| 70 | Levitin | 2016 | 3 | Siemens | Trio | 12 | MPRAGE | 2100 | 2.4 | - | EPI | 3000 | 30 | - | SPM12 | RDA |
| 71 | Li | 2019 | 3 | Siemens | - | - | T1-w | 2000 | 2.52 | | continuous | 2000 | 26 | - | SPM12 | GLM |
| 72 | Limb | 2006 | 3 | GE | - | quadrature | T1-w | - | - | - | EPI | 2000 | 30 | - | SPM99 | GLM |
| 73 | Limb | 2008 | 3 | GE | - | quadrature | - | - | - | - | EPI | 2000 | 30 | - | SPM99 | GLM |
| 74 | Liu | 2018 | 3 | GE | Signa | 8 | MPRAGE | - | - | - | EPI | 2000 | 30 | - | SPM8 | ICA |
| 75 | Matthews | 2020 | 3 | Siemens | Tim Trio | 32 | T1-w | 2420 | 3.7 | 1x1x1 | EPI | 2000 | - | 2.35x2.53x2.50 | SPM12 | GLM |
| 76 | Meister | 2004 | 3 | Philips | Gyroscan | quadrature | - | - | - | - | EPI | 3587 | 50 | - | SPM99 | LME |
| 77 | Merrill | 2012 | 3 | Siemenss | Trio | - | T1-w | 1300 | 7.4 | - | EPI | 2500 | 30 | 3x3x3 | SPM8 | GLM |
| 78 | Mizuno | 2007 | 1.5 | GE | Signa | - | T1-w | - | - | - | EPI | 2000 | 50 | - | SPM99 | GLM |
| 79 | Montag | 2011 | 1.5 | Siemens | Avanto | - | - | - | - | - | EPI | 3200 | 40 | - | SMP5 | GLM |
| 80 | Morrison | 2003 | 1.5 | GE | - | - | - | - | - | - | EPI | 2500 | 50 | - | MEDx | 3.4.1 |
| 81 | Mueller | 2015 | 3 | Brucker | Medspec | birdcage | MPRAGE | - | - | 1x1x1 | ISSS | - | - | 2.5x2.5x2.5 | SPM8 | GLM |
| 82 | Ohnishi | 2001 | 1.5 | Siemens | Magentom | - | - | - | - | - | - | 3000 | - | - | SPM99 | GLM |
| 83 | Park | 2013 | 3 | Siemens | Magnetom | TIM | MPRAGE | 2400 | 3.06 | - | EPI | 3000 | 30 | - | BrainVoyager | GLM |
| 84 | Park | 2014 | 3 | Siemens | Magentom | - | MPRAGE | 2400 | 3.06 | - | EPI | 3000 | 30 | - | SPM8 | GLM |
| 85 | Parsons | 2005 | - | GE | - | - | - | - | - | 1x1x1 | - | - | - | - | - | - |
| 86 | Pereira | 2011 | 1.5 | Phillips | Gyroscan | - | SPGR | - | - | 1x1x1 | EPI | 3000 | 50 | 3.56x3.56x4.0 | FSL | GLM |
| 87 | Peretz | 2009 | 1.5 | Siemens | Sonata | - | T1-1 | - | - | 1x1x1 | EPI | 1150 | 50 | 5x5x7 | Matlab | GLM |
| 88 | Petrini | 2011 | 3 | Siemens | Tim Trio | - | MPRAGE | 1900 | 2.52 | | EPI | 2000 | 30 | 3x3x3 | BrainVoyager | GLM |
| 89 | Pfordresher | 2014 | 3 | GE | Signa excite | 8 | - | - | - | - | EPI | 2000 | 35 | 1x1x1 | SPM5 | Box car |
| 90 | Ragert | 2014 | 3 | Siemens | Tim Trio | birdcage | T1-w | - | - | 1x1x1 | EPI | 2000 | 28 | 3x3x3 | model | GLM |
| 91 | Reiterer | 2008 | 1.5 | Siemens | Vision | - | - | - | - | - | EPI | 3000 | 40 | - | SPM2 | GLM |
| 92 | Rogalsky | 2011 | 3 | Siemens | Achieva | - | SPGR | 2500 | 1.3 | 1x1x1 | EPI | 2000 | 40 | 1x1x1 | AFNI | Regression |
| 93 | Sammler | 2010 | 3 | Siemens | Trio | - | MPRAGE | 2300 | 4.8 | 1x1x1 | EPI | 2120 | 25 | 3x3x3 | SPM5 | GLM |
| 94 | Schmithorst | 2005 | 3 | Bruker | Medspec | | T1-w | - | - | - | EPI | 3000 | 38 | - | SPM | ICA |
| 95 | Schön | 2010 | 3 | Bruker | Medspec | - | T1-w | - | - | - | EPI | 2166 | - | 3.5 | SPM2 | GLM |
| 96 | Schwenzer | 2011 | 1.5 | Siemens | Magnetom | - | - | - | - | - | EPI | 6000 | 17 | 3.6x3.6x4 | SPM2 | GLM |
| 97 | Shany | 2019 | 3 | GE | Signa | 8-channel | SPGR | 8.9 | 3.5 | 1x1x1 | EPI | 3000 | 35 | 3 | BrainVoyager | GLM |
| 98 | Sikka | 2015 | 3 | Siemens | Magnetom | 12-channel | MPRAGE | 1760 | 2.2 | 1x1x1 | EPI | 10500 | 30 | 3.3 | SPM8 | GLM |
| 99 | Singer | 2016 | 3 | GE | Signa | 8-channel | SPGR | 8.9 | 3.5 | 1x1x1 | EPI | 3000 | 35 | 3 | BrainVoyager | DCA |
| 100 | Skouras | 2014 | 3 | Siemens | Magnetom | 12-channel | MPRAGE | - | - | 1x1x1 | EPI | 2000 | 30 | 3 | LIPSIA | GLM |
| 101 | Spada | 2014 | 3 | Philips | Achieva | - | T1-w | 7.3 | 3.5 | 1x1x1 | EPI | 12000 | 30 | - | SPM8 | GLM |
| 102 | Tabei | 2015 | 1.5 | Siemens | Symphony | - | T1-w | 2200 | 3.93 | 1x1x1 | EPI | 4000 | 50 | 3 | SPM5 | GLM |
| 103 | Tachibana | 2010 | 3 | Siemens | Trio | - | T1-w | 2250 | 3.06 | 1x1x1 | EPI | 6000 | 30 | 3x3x5 | SPM5 | GLM |
| 104 | Taruffi | 2017 | 3 | Siemens | Trio | - | MPRAGE | - | - | 1x1x1 | EPI | 2250 | 30 | 3 | LIPSIA | ECM |
| 105 | Tervaniemi | 2000 | - | - | - | - | - | - | - | - | - | - | - | - | - | - |
| 106 | Tervaniemi | 2006 | 3 | Bruker | Medspec | - | T1-w | - | - | - | EPI | 3000 | 30 | - | LIPSIA | GLM |
| 107 | Thaut | 2008 | 1.5 | Siemens | Magnetom | quadrature | MPRAGE | 11.4 | 4.4 | 1x1x1 | EPI | 3000 | 40 | 2.7x2.7x2.7 | SPM99 | Box-car |
| 108 | Tillmann | 2003 | 1.5 | GE | Signa | - | T1-w | 650 | 6.6 | 0.937x0.937x5.0) | EPI | 2000 | 35 | 3.75x3.75x5 | SPM99 | GLM |
| 109 | Tillmann | 2006 | 3 | Siemens | Trio | - | T1-w | 1300 | 7.4 | 0.8x0.8x4 | EPI | 2000 | 30 | 3x3x4 | LIPSIA | GLM |
| 110 | Toiviainen | 2014 | 3 | GE | - | standard | T1-w | 6552 | 2.82 | 0.94x0.94x1.2 | EPI | 2200 | 30 | 3x3x3 | SPM8 | LASSO |
| 111 | Trost | 2012 | 3 | Siemens | Trio | - | MPRAGE | 1900 | 2.32 | 0.9x0.9x0.9 | EPI | 3000 | 30 | 3.5 | SPM5 | GLM |
| 112 | Trost | 2014 | 3 | Siemens | Trio | 12-channel | MPRAGE | 1900 | 2.32 | 0.9x0.9x0.9 | EPI | 1980 | 27.3 | 3.2 | SPM8 | GLM |
| 113 | Tsai | 2010 | 3 | Bruker | - | - | T1-w | - | - | 0.94x0.94x5 | EPI | 2000 | 30 | 3.75x3.75x5 | SPM2 | GLM |
| 114 | Tsai | 2012 | 3 | Bruker | - | - | T1-w | - | - | 0.94x0.94x5 | EPI | 3000 | 30 | 3.75x3.75x5 | SPM5 | GLM |
| 115 | Tsai | 2018 | 3 | Siemens | Magnetom | 20-channel | MPRAGE | - | - | 0.9x0.9x0.9 | EPI | 2500 | 30 | - | SPM12 | GLM |
| 116 | Tsai | 2019 | 3 | Siemens | Magnetom | 20-channel | MPRAGE | 2000 | 2.3 | 0.93x0.93x0.93 | EPI | 2500 | 30 | 2.5 | SPM12 | GLM |

| | | | | | | | | | | | | | | | | |
|---|---|---|---|---|---|---|---|---|---|---|---|---|---|---|---|---|
| 117 | Uhlig | 2013 | 3 | Siemens | Trio | Birdcage | - | - | - | - | EPI | 2000 | 28 | 3x3x3 | FSL | GLM |
| 118 | Villarreal | 2013 | 3 | GE | HDx | 8-channel | SPGR-IR | 13 | 6.1 | - | EPI | 2300 | 35 | 3.75x3.75x4 | SPM5 | GLM |
| 119 | Vuust | 2011 | 1.5 | GE | - | - | SPGR | 30 | - | - | EPI | 3200 | 40 | 4 | SPM2 | Box-car |
| 120 | Wallmark | 2018 | 3 | Siemens | Trio | - | MPRAGE | 1900 | 2.26 | - | EPI | 5000 | 34 | - | FSL | GLM |
| 121 | Watanabe | 2008 | 1.5 | Hitachi | - | - | T1-w | - | - | - | EPI | 14000 | 50 | 4x4x5 | SPM2 | LME |
| 122 | Whitehead | 2018 | - | - | - | - | - | - | - | - | multi-band | 529 | 35 | 2x2x2 | AFNI | GLM |
| 123 | Wilson | 2010 | 3 | GE | Signa XL | Birdcage | - | - | - | - | EPI | 3000 | 40 | 1.88x1.88x1 | SPM8 | GLM |
| 124 | Yoo | 2001 | 1.5 | Siemens | Magnetom | Birdcage | T1-w | 700 | 10 | - | EPI | 2500 | 50 | - | SPM99 | GLM+GRF |
| 125 | Zarate | 2008 | 1.5 | Siemens | Sonata | - | T1-w | - | - | 1x1x1 | T2*-w | 10000 | 85 | 5x5x5 | AFNI | GLM |
| 126 | Zarate | 2010 | 3 | Siemens | Trio | - | T1-w | - | - | 1x1x1 | T2*-w | 60000 | 10300 | 3.5x3.5x3.5 | AFNI | GLM |
| 127 | Zatorre | 1994 | 1.5 | Philips | Gyroscan | - | - | - | - | - | - | - | - | 1.5x1.5.1.5 | - | MSDCI |
| 128 | Zatorre | 1996 | 1.5 | Philips | Gyroscan | - | - | - | - | - | - | - | - | - | AFNI | RFT |
| 129 | Zatorre | 2010 | 3 | Siemens | Trio | 8-channel | T1-w | - | - | 1x1x1 | EPI | 2400 | | 3.5x3.5x3.5 | fMRISTAT | GLM |
| 130 | Zvyagintsev | 2013 | 3 | Siemens | Trio | 12-channel | MPRAGE | 2300 | 2.98 | 1x1x1 | EPI | 2000 | 28 | 3x3x3 | BrainVoyager | GLM |

GM, grey matter; WM, white matter; MRI, magnetic resonance imaging; FFE, fast field echo sequence; FLASH, fast low angle shot sequence; FSL, functional MRI of the brain software library; GE, gradient echo pulse; IR-FSPGR, fast spoiled gradient sequence with inversion preparation; MPRAGE, magnetization-prepared rapid acquisition with gradient echo sequence; MDEFT, modified driven equilibrium Fourier transform; SPGR, spoiled gradient recalled sequence; SPM, statistical parametric mapping; TFE, turbo field echo sequence; VBM, voxel-based morphometry.

**Supplementary Table 2. MRI quality.**

| | Author | Year | MRI design described | Age reported | Sample gender reported | Sample handedness reported | Ethics approval reported | Image acquisition described | Image processing described | Statistical MRI-analysis described | Software package specified | Multiple comparison correction | Figures and tables |
|---|---|---|---|---|---|---|---|---|---|---|---|---|---|
| 1 | Agustus | 2018 | Y | Y | Y | Y | Y | Y | Y | Y | Y | Y | Y |
| 2 | Alluri | 2012 | Y | Y | Y | Y | Y | Y | Y | Y | Y | Y | Y |
| 3 | Alluri | 2013 | Y | Y | Y | Y | Y | Y | Y | Y | Y | Y | Y |
| 4 | Alonso | 2016 | Y | Y | Y | Y | Y | Y | Y | Y | Y | N | Y |
| 5 | Alttenmüller | 2014 | Y | Y | Y | Y | Y | Y | Y | Y | Y | Y | Y |
| 6 | Angulo-P | 2014 | Y | Y | Y | Y | Y | Y | Y | Y | Y | Y | Y |
| 7 | Armony | 2015 | Y | Y | Y | Y | Y | Y | Y | Y | Y | N | Y |
| 8 | Bangert | 2006 | Y | Y | Y | Y | Y | Y | Y | Y | Y | Y | Y |
| 9 | Barrett | 2016 | Y | Y | Y | N | Y | Y | Y | Y | Y | Y | Y |
| 10 | Barrett | 2018 | Y | N | N | N | Y | Y | Y | Y | Y | Y | Y |
| 11 | Barrett | 2020 | Y | Y | Y | N | Y | Y | Y | Y | Y | Y | Y |
| 12 | Bastepe-G | 2020 | Y | Y | Y | N | Y | Y | Y | Y | Y | Y | Y |
| 13 | Baumann | 2007 | Y | Y | Y | Y | Y | Y | Y | Y | Y | Y | Y |
| 14 | Bengtsson | 2006 | Y | Y | Y | Y | Y | Y | Y | Y | Y | Y | Y |
| 15 | Bengtsson | 2007 | Y | Y | Y | Y | Y | Y | Y | Y | Y | Y | Y |
| 16 | Bengtsson | 2009 | Y | Y | Y | Y | Y | Y | Y | Y | Y | Y | Y |
| 17 | Bianco | 2016 | Y | Y | Y | N | Y | Y | Y | Y | Y | N | Y |
| 18 | Bishop | 2013 | Y | Y | Y | N | Y | Y | Y | Y | Y | N | Y |
| 19 | Blood | 1999 | Y | N | N | N | N | Y | Y | Y | N | N | Y |
| 20 | Bodner | 2001 | Y | N | N | N | Y | Y | Y | Y | Y | N | Y |
| 21 | Bogert | 2016 | Y | Y | Y | Y | Y | Y | Y | Y | Y | Y | Y |
| 22 | Brattico | 2016 | Y | Y | Y | Y | Y | Y | Y | Y | Y | Y | Y |
| 23 | Bravo | 2017a | Y | Y | Y | Y | | Y | Y | Y | Y | Y | Y |
| 24 | Bravo | 2017b | Y | Y | Y | Y | Y | Y | Y | Y | Y | Y | Y |
| 25 | Bravo | 2020 | Y | Y | Y | Y | Y | Y | Y | Y | Y | Y | Y |
| 26 | Brown | 2004 | Y | Y | Y | N | Y | Y | Y | Y | Y | Y | Y |
| 27 | Brown | 2007 | Y | Y | Y | N | Y | Y | N | N | N | N | Y |
| 28 | Chapin | 2010 | Y | Y | Y | N | Y | Y | Y | Y | Y | Y | Y |
| 29 | Chen | 2008a | Y | N | N | N | Y | Y | Y | Y | Y | Y | Y |
| 30 | Chen | 2008b | Y | Y | Y | N | Y | Y | Y | Y | Y | Y | Y |
| 31 | Chiang | 2018 | Y | Y | Y | Y | Y | Y | Y | Y | Y | Y | Y |
| 32 | Danielsen | 2014 | Y | Y | Y | N | Y | Y | Y | Y | Y | Y | Y |
| 33 | Demorest | 2009 | Y | Y | Y | Y | Y | Y | Y | Y | Y | Y | Y |
| 34 | Donnay | 2014 | Y | Y | Y | Y | Y | U | U | U | U | U | Y |
| 35 | Engel | 2011 | Y | Y | Y | Y | Y | Y | Y | Y | Y | Y | Y |
| 36 | Escoffier | 2013 | Y | Y | Y | Y | Y | Y | Y | Y | Y | Y | Y |
| 37 | Fedorenko | 2012 | Y | N | Y | N | Y | Y | Y | Y | Y | Y | Y |
| 38 | Flores-G | 2007 | Y | Y | Y | N | Y | Y | Y | Y | Y | Y | Y |
| 39 | Fujisawa | 2011 | Y | Y | Y | N | Y | Y | Y | Y | Y | N | Y |
| 40 | González-G | 2016 | Y | Y | Y | Y | Y | Y | Y | Y | Y | Y | Y |
| 41 | Grahn | 2007 | Y | Y | Y | N | Y | Y | Y | Y | Y | Y | Y |
| 42 | Grahn | 2009a | Y | Y | Y | Y | Y | Y | Y | Y | Y | N | Y |
| 43 | Grahn | 2009b | Y | Y | Y | Y | Y | Y | Y | Y | Y | Y | Y |
| 44 | Grahn | 2013 | Y | Y | Y | Y | Y | Y | Y | Y | Y | U | Y |

| | | | | | | | | | | | | | |
|---|---|---|---|---|---|---|---|---|---|---|---|---|---|
| 45 | Green | 2008 | Y | Y | N | N | Y | N | N | N | Y | N | N |
| 46 | Green | 2012 | Y | Y | Y | Y | Y | Y | Y | Y | Y | N | Y |
| 47 | Green | 2018 | Y | Y | Y | Y | Y | Y | Y | Y | Y | Y | Y |
| 48 | Halpern | 1999 | Y | Y | Y | N | Y | Y | N | N | N | N | Y |
| 49 | Halpern | 2004 | Y | Y | Y | Y | Y | Y | Y | Y | Y | N | Y |
| 50 | Herdener | 2014 | N | N | N | N | Y | N | N | Y | Y | Y | Y |
| 51 | Herholtz | 2012 | Y | Y | Y | Y | Y | Y | Y | Y | Y | Y | Y |
| 52 | Huang | 2016 | Y | N | Y | N | Y | Y | Y | Y | Y | Y | Y |
| 53 | Janata | 2002a | Y | Y | Y | Y | Y | Y | Y | Y | Y | Y | Y |
| 54 | Janata | 2002b | Y | Y | Y | Y | Y | Y | Y | Y | Y | Y | Y |
| 55 | Janata | 2009 | Y | Y | Y | N | Y | Y | Y | Y | Y | N | Y |
| 56 | Jeong | 2011 | Y | Y | Y | Y | Y | Y | Y | Y | Y | Y | Y |
| 57 | Jungblut | 2012 | Y | Y | N | Y | Y | Y | Y | Y | Y | Y | Y |
| 58 | Khalfa | 2005 | Y | Y | Y | Y | Y | Y | Y | Y | Y | Y | Y |
| 59 | Kleber | 2007 | Y | Y | Y | Y | Y | Y | Y | Y | Y | Y | Y |
| 60 | Kleider-O | 2019 | Y | Y | Y | Y | Y | Y | Y | Y | Y | Y | Y |
| 61 | Koelsch | 2003 | Y | Y | Y | N | Y | Y | Y | Y | Y | Y | Y |
| 62 | Koelsch | 2014 | Y | Y | Y | N | Y | Y | Y | Y | Y | Y | Y |
| 63 | Koelsch | 2018 | Y | Y | Y | Y | Y | Y | Y | Y | Y | Y | Y |
| 64 | Kornysheva | 2010 | Y | Y | Y | Y | Y | Y | Y | Y | Y | Y | Y |
| 65 | Langheim | 2002 | Y | Y | Y | Y | Y | Y | Y | Y | Y | Y | Y |
| 66 | Leaver | 2009 | Y | Y | Y | Y | Y | Y | Y | Y | Y | Y | Y |
| 67 | Lee | 2011 | Y | Y | Y | Y | Y | Y | Y | Y | Y | N | Y |
| 68 | Lehne | 2013 | Y | Y | Y | Y | Y | Y | Y | Y | Y | Y | Y |
| 69 | Levitin | 2005 | Y | Y | Y | Y | Y | Y | N | N | N | N | N |
| 70 | Levitin | 2016 | Y | Y | Y | Y | Y | Y | Y | Y | Y | N | Y |
| 71 | Li | 2019 | Y | Y | Y | Y | Y | Y | Y | Y | Y | Y | Y |
| 72 | Limb | 2006 | Y | Y | Y | Y | Y | Y | Y | Y | Y | Y | Y |
| 73 | Limb | 2008 | Y | Y | Y | Y | Y | Y | Y | Y | Y | Y | Y |
| 74 | Liu | 2018 | Y | Y | Y | N | Y | Y | Y | Y | Y | N | Y |
| 75 | Matthews | 2020 | Y | Y | Y | N | Y | Y | Y | Y | Y | Y | Y |
| 76 | Meister | 2004 | Y | Y | Y | Y | Y | Y | Y | Y | Y | N | Y |
| 77 | Merrill | 2012 | Y | Y | Y | Y | Y | Y | Y | Y | Y | Y | Y |
| 78 | Mizuno | 2007 | Y | Y | Y | Y | Y | Y | Y | Y | Y | Y | Y |
| 79 | Montag | 2011 | Y | Y | Y | N | Y | Y | Y | Y | Y | Y | Y |
| 80 | Morrison | 2003 | Y | Y | Y | N | Y | Y | Y | Y | Y | N | Y |
| 81 | Mueller | 2015 | Y | Y | Y | N | Y | Y | Y | Y | Y | Y | Y |
| 82 | Ohnishi | 2001 | Y | Y | Y | N | Y | Y | Y | Y | Y | N | Y |
| 83 | Park | 2013 | Y | Y | Y | Y | Y | Y | Y | Y | Y | Y | Y |
| 84 | Park | 2014 | Y | Y | Y | Y | Y | Y | Y | Y | Y | Y | Y |
| 85 | Parsons | 2005 | Y | Y | Y | Y | Y | Y | Y | Y | Y | N | Y |
| 86 | Pereira | 2011 | Y | Y | Y | Y | Y | Y | Y | Y | Y | Y | Y |
| 87 | Peretz | 2009 | Y | Y | Y | Y | Y | Y | Y | Y | Y | Y | Y |
| 88 | Petrini | 2011 | Y | Y | Y | N | Y | Y | Y | Y | Y | Y | Y |
| 89 | Pfordresher | 2014 | Y | Y | Y | Y | Y | Y | Y | Y | Y | Y | Y |
| 90 | Ragert | 2014 | Y | Y | Y | Y | Y | Y | Y | Y | Y | Y | Y |
| 91 | Reiterer | 2008 | Y | Y | Y | Y | Y | Y | Y | Y | Y | Y | Y |
| 92 | Rogalsky | 2011 | Y | Y | Y | Y | Y | Y | Y | Y | Y | Y | Y |
| 93 | Sammler | 2010 | Y | Y | Y | Y | Y | Y | Y | Y | Y | Y | Y |
| 94 | Schmithorst | 2005 | Y | Y | Y | Y | Y | Y | Y | Y | Y | Y | Y |
| 95 | Schön | 2010 | Y | Y | Y | Y | Y | U | Y | Y | Y | Y | Y |
| 96 | Schwenzer | 2011 | Y | U | Y | N | N | U | U | Y | Y | Y | Y |
| 97 | Shany | 2019 | Y | Y | Y | Y | Y | Y | Y | Y | Y | Y | Y |

| | | | | | | | | | | | | | |
|---|---|---|---|---|---|---|---|---|---|---|---|---|---|
| 98 | Sikka | 2015 | Y | Y | Y | Y | Y | Y | Y | Y | Y | Y | Y |
| 99 | Singer | 2016 | Y | Y | Y | Y | Y | Y | Y | Y | Y | Y | Y |
| 100 | Skouras | 2014 | Y | Y | Y | Y | Y | Y | Y | Y | Y | Y | Y |
| 101 | Spada | 2014 | Y | Y | Y | Y | Y | Y | Y | Y | Y | Y | Y |
| 102 | Tabei | 2015 | Y | Y | Y | Y | Y | Y | Y | Y | Y | Y | Y |
| 103 | Tachibana | 2010 | Y | Y | Y | Y | Y | Y | Y | Y | Y | Y | Y |
| 104 | Taruffi | 2017 | Y | Y | Y | Y | Y | Y | Y | Y | Y | Y | Y |
| 105 | Tervaniemi | 2000 | Y | Y | Y | Y | Y | Y | Y | Y | Y | Y | Y |
| 106 | Tervaniemi | 2006 | Y | Y | Y | Y | Y | Y | Y | Y | Y | N | Y |
| 107 | Thaut | 2008 | Y | Y | Y | Y | Y | Y | Y | Y | Y | Y | Y |
| 108 | Tillmann | 2003 | Y | Y | Y | Y | N | Y | Y | Y | Y | N | Y |
| 109 | Tillmann | 2006 | Y | Y | Y | Y | N | Y | Y | Y | Y | N | Y |
| 110 | Toiviainen | 2014 | Y | Y | Y | Y | Y | Y | Y | Y | Y | Y | Y |
| 111 | Trost | 2012 | Y | Y | Y | Y | Y | Y | Y | Y | Y | Y | Y |
| 112 | Trost | 2014 | Y | Y | Y | Y | Y | Y | Y | Y | Y | Y | N |
| 113 | Tsai | 2010 | Y | U | Y | Y | Y | Y | Y | Y | Y | N | Y |
| 114 | Tsai | 2012 | Y | Y | Y | N | Y | Y | Y | Y | Y | Y | Y |
| 115 | Tsai | 2018 | Y | Y | Y | N | Y | Y | Y | Y | Y | Y | Y |
| 116 | Tsai | 2019 | Y | Y | Y | N | Y | Y | Y | Y | Y | Y | Y |
| 117 | Uhlig | 2013 | Y | Y | Y | Y | Y | U | Y | Y | Y | Y | Y |
| 118 | Villarreal | 2013 | Y | Y | Y | Y | Y | Y | Y | Y | Y | Y | Y |
| 119 | Vuust | 2011 | Y | Y | Y | Y | Y | Y | Y | Y | Y | Y | Y |
| 120 | Wallmark | 2018 | Y | Y | Y | Y | Y | Y | Y | Y | Y | Y | Y |
| 121 | Watanabe | 2008 | Y | Y | Y | Y | Y | Y | Y | Y | Y | N | Y |
| 122 | Whitehead | 2018 | Y | Y | Y | Y | N | U | U | Y | Y | Y | Y |
| 123 | Wilson | 2010 | Y | Y | Y | Y | Y | Y | Y | Y | Y | Y | Y |
| 124 | Yoo | 2001 | Y | Y | Y | N | Y | U | U | Y | Y | N | Y |
| 125 | Zarate | 2008 | Y | N | Y | Y | Y | Y | U | Y | Y | Y | Y |
| 126 | Zarate | 2010 | Y | Y | Y | Y | Y | U | U | Y | Y | Y | Y |
| 127 | Zatorre | 1994 | Y | N | Y | Y | N | N | N | N | N | Y | Y |
| 128 | Zatorre | 1996 | Y | Y | Y | Y | N | N | N | U | Y | Y | Y |
| 129 | Zatorre | 2010 | Y | Y | Y | Y | Y | Y | Y | Y | Y | N | Y |
| 130 | Zvyagintsev | 2013 | Y | Y | Y | Y | Y | Y | Y | Y | Y | Y | Y |

Y, yes; N, no; U, unclear.

**Supplementary Table 3. Contrast analyses comparing music perception, production and imagery, at cluster level inference p < 0.05 (FWE).**

| Cluster number | Volume (mm³) | MNI coordinates x | y | z | ALE | P | Z | Label (Side, region) |
|---|---|---|---|---|---|---|---|---|
| ***a. Perception + Production*** | | | | | | | | |
| 1 | 3448 | 66 | -24 | 10 | 2E-02 | - | - | R Superior Temporal Gyrus BA42 |
| | | 50 | -20 | 6 | 2E-02 | - | - | R Superior Temporal Gyrus BA13 |
| | | 64 | -26 | 4 | 2E-02 | - | - | R Superior Temporal Gyrus BA22 |
| | | 62 | -22 | 2 | 2E-02 | - | - | R Superior Temporal Gyrus BA41 |
| | | 50 | -10 | 4 | 2E-02 | - | - | R Insula BA13 |
| | | 64 | -28 | 14 | 2E-02 | - | - | R Superior Temporal Gyrus BA42 |
| | | 68 | -16 | 8 | 2E-02 | - | - | R Transverse Temporal Gyrus BA42 |
| 2 | 1816 | -42 | -28 | 6 | 3E-02 | - | - | L Superior Temporal Gyrus BA13 |
| 3 | 152 | 0 | -4 | 60 | 2E-02 | - | - | L Medial Frontal Gyrus BA6 |
| 4 | 88 | -4 | 6 | 54 | 2E-02 | - | - | L Medial Frontal Gyrus BA6 |
| 5 | 24 | 50 | 10 | 24 | 1E-02 | - | - | R Inferior Frontal Gyrus BA9 |
| ***b. Perception - Production*** | | | | | | | | |
| 1 | 1096 | -56 | -16 | -4 | - | 8E-03 | 2.4 | L Superior Temporal Gyrus BA21 |
| 2 | 168 | 50 | -18 | -6 | - | 2E-02 | 2.1 | R Superior Temporal Gyrus BA22 |
| 3 | 112 | 52 | -10 | -12 | - | 3E-02 | 1.9 | R Superior Temporal Gyrus BA22 |
| ***c. Production - Perception*** | | | | | | | | |
| 1 | 2816 | -43.7 | -23.7 | 45.4 | - | 1E-04 | 3.7 | L Postcentral Gyrus BA2 |
| | | -42.7 | -24 | 47.6 | - | 1E-04 | 3.7 | L Postcentral Gyrus BA40 |
| | | -40 | -21 | 44 | - | 7E-04 | 3.2 | L Postcentral Gyrus BA2 |
| | | -44 | -12 | 42 | - | 1E-03 | 3.0 | L Precentral Gyrus BA4 |
| | | -34 | -44 | 49 | - | 2E-03 | 2.9 | L Inferior Parietal Lobule BA40 |
| | | -48 | -10 | 42 | - | 3E-03 | 2.8 | L Precentral Gyrus BA4 |
| | | -50 | -6 | 40 | - | 3E-03 | 2.8 | L Precentral Gyrus BA6 |
| 2 | 2128 | -8 | -7 | 51 | - | 0E+00 | 3.9 | L Medial Frontal Gyrus BA6 |
| | | -12 | -9 | 50 | - | 1E-04 | 3.7 | L Medial Frontal Gyrus BA6 |
| | | -12 | -7 | 54 | - | 2E-04 | 3.5 | L Medial Frontal Gyrus BA6 |
| | | -4 | -2 | 56 | - | 2E-03 | 2.9 | L Medial Frontal Gyrus BA6 |
| | | 4 | 0 | 54 | - | 3E-03 | 2.8 | R Medial Frontal Gyrus BA6 |
| | | -2 | 6 | 44 | - | 1E-02 | 2.2 | L Cingulate Gyrus BA24 |
| 3 | 1040 | 54.3 | 3.4 | 29.9 | - | 0E+00 | 3.9 | R Precentral Gyrus BA6 |
| 4 | 608 | -42 | -34 | 4 | - | 8E-03 | 2.4 | L Superior Temporal Gyrus BA41 |
| | | -42 | -38 | 8 | - | 8E-03 | 2.4 | L Superior Temporal Gyrus BA41 |
| | | -42 | -28 | 6 | - | 1E-02 | 2.3 | L Superior Temporal Gyrus BA13 |
| | | -41 | -28 | 0 | - | 1E-02 | 2.3 | L Insula BA13 |
| 5 | 584 | -20 | -60 | -22 | - | 7E-04 | 3.2 | L Cerebellum |
| 6 | 536 | 56 | 8 | -4 | - | 3E-03 | 2.8 | R Superior Temporal Gyrus BA22 |
| 7 | 432 | -50 | 4 | 0 | - | 7E-03 | 2.5 | L Superior Temporal Gyrus BA22 |
| 8 | 424 | 2 | 22 | 38 | - | 3E-03 | 2.8 | L Cingulate Gyrus BA32 |
| 9 | 400 | -62 | -42 | 20 | - | 2E-02 | 2.1 | L Superior Temporal Gyrus BA22 |
| 10 | 352 | 68 | -22 | 12 | - | 1E-02 | 2.3 | R Superior Temporal Gyrus BA42 |
| | | 66 | -24 | 16 | - | 2E-02 | 2.1 | R Superior Temporal Gyrus BA42 |
| 11 | 320 | -30 | 0 | 2 | - | 2E-03 | 2.8 | L Putamen |
| | | -24 | -4 | 2 | - | 7E-03 | 2.4 | L Putamen |
| 12 | 288 | -60 | -4 | 2 | - | 3E-03 | 2.8 | L Superior Temporal Gyrus BA22 |
| 13 | 216 | 28 | 6 | -10 | - | 6E-03 | 2.5 | R Putamen |
| 14 | 200 | 44 | 0 | 50 | - | 4E-03 | 2.6 | R Middle Frontal Gyrus BA6 |
| 15 | 168 | -4 | 14 | 62 | - | 6E-03 | 2.5 | L Superior Frontal Gyrus BA6 |
| 16 | 120 | -68 | -18 | 6 | - | 2E-02 | 2.1 | L Superior Temporal Gyrus BA22 |
| 17 | 112 | -36 | 20 | 2 | - | 2E-02 | 2.0 | L Insula BA13 |
| ***d. Production + Imagery*** | | | | | | | | |
| 1 | 800 | -2 | -2 | 58 | 2E-02 | - | - | L Medial Frontal Gyrus BA6 |
| | | -4 | 4 | 56 | 2E-02 | - | - | L Medial Frontal Gyrus BA6 |
| ***e. Production - Imagery*** | | | | | | | | |
| 1 | 1824 | -36 | -28 | 7 | - | 6E-04 | 3.2 | L Superior Temporal Gyrus BA13 |
| | | -44 | -30 | 8 | - | 9E-04 | 3.1 | L Superior Temporal Gyrus BA41 |
| | | -40 | -29.3 | 13.3 | - | 2E-03 | 2.9 | L Superior Temporal Gyrus BA41 |
| | | -38 | -36 | 12 | - | 2E-03 | 3.0 | L Transverse Temporal Gyrus BA41 |
| | | -41 | -24 | 12 | - | 4E-03 | 2.7 | L Transverse Temporal Gyrus BA41 |
| 2 | 344 | -42 | -24 | 54 | - | 1E-02 | 2.2 | L Postcentral Gyrus BA40 |
| 3 | 272 | 48 | -24 | 4 | - | 8E-03 | 2.4 | R Superior Temporal Gyrus BA22 |

| | | | | | | | | |
|---|---|---|---|---|---|---|---|---|
| | | 52 | -26 | 4 | - | 2E-02 | 2.0 | R Superior Temporal Gyrus BA41 |
| 4 | 240 | -18 | -60 | -18 | - | 2E-02 | 2.0 | L Cerebellum |
| | | -20 | -64 | -20 | - | 3E-02 | 1.8 | L Cerebellum |
| 5 | 232 | 54 | 2 | -2 | - | 8E-03 | 2.4 | R Superior Temporal Gyrus BA22 |
| ***f. Imagery - Production*** | | | | | | | | |
| 1 | 2552 | -36.7 | -58.7 | 59.3 | - | 4E-04 | 3.4 | L Superior Parietal Lobule BA7 |
| | | -36 | -64 | 56 | - | 1E-03 | 3.0 | L Superior Parietal Lobule BA7 |
| | | -30 | -54 | 38 | - | 4E-03 | 2.6 | L Angular Gyrus BA39 |
| 2 | 760 | 0 | 4 | 62 | - | 5E-03 | 2.6 | L Medial Frontal Gyrus BA6 |
| 3 | 696 | 56 | 6 | 48 | - | 9E-03 | 2.4 | R Precentral Gyrus BA6 |
| ***g. Imagery + Perception*** | | | | | | | | |
| 1 | 1848 | 0 | 6 | 58 | 2E-02 | - | - | L Medial Frontal Gyrus BA6 |
| | | -2 | 0 | 62 | 2E-02 | - | - | L Medial Frontal Gyrus BA6 |
| 2 | 624 | 54 | 0 | 50 | 2E-02 | - | - | R Precentral Gyrus BA6 |
| 3 | 72 | -20 | -2 | 6 | 1E-02 | - | - | L Lateral Globus Pallidus |
| ***h. Imagery - Perception*** | | | | | | | | |
| 1 | 3064 | -32.6 | -56.9 | 53.6 | - | 0E+00 | 3.9 | L Inferior Parietal Lobule BA7 |
| | | -38 | -60 | 50 | - | 2E-04 | 3.5 | L Inferior Parietal Lobule BA7 |
| | | -32 | -50 | 38 | - | 1E-03 | 3.0 | L Inferior Parietal Lobule BA40 |
| 2 | 2672 | -3.1 | 2.2 | 56.4 | - | 0E+00 | 3.9 | L Medial Frontal Gyrus BA6 |
| | | 2 | 5.3 | 56.7 | - | 1E-04 | 3.7 | L Medial Frontal Gyrus BA6 |
| 3 | 1248 | -16 | -10 | 4 | - | 2E-03 | 2.8 | L Thalamus |
| | | -18 | -18 | 8 | - | 3E-03 | 2.7 | L Thalamus |
| | | -18 | -6 | 14 | - | 1E-02 | 2.2 | L Thalamus |
| 4 | 296 | 52 | 10 | 48 | - | 2E-03 | 3.0 | R Middle Frontal Gyrus BA6 |
| 5 | 144 | 56 | -2 | 54 | - | 2E-02 | 2.2 | R Precentral Gyrus BA4 |
| ***i. Perception - Imagery*** | | | | | | | | |
| 1 | 4936 | 46 | -24 | 2 | - | 5E-04 | 3.3 | R Superior Temporal Gyrus BA22 |
| | | 52 | -4 | 0 | - | 8E-03 | 2.4 | R Superior Temporal Gyrus BA22 |
| | | 40 | -14 | -2 | - | 2E-02 | 2.2 | R Claustrum |
| | | 62 | -26 | -2 | - | 3E-02 | 1.9 | R Superior Temporal Gyrus BA22 |
| 2 | 3272 | -44 | -28 | 12 | - | 3E-03 | 2.8 | L Transverse Temporal Gyrus BA41 |
| | | -34 | -30 | 8 | - | 4E-03 | 2.7 | L Transverse Temporal Gyrus BA41 |
| 3 | 984 | -44 | -6 | -6 | - | 8E-03 | 2.4 | L Insula BA13 |
| | | -50 | -10 | -2 | - | 2E-02 | 2.0 | L Superior Temporal Gyrus BA22 |
| | | -46 | 0 | 4 | - | 2E-02 | 2.0 | L Insula BA13 |
| 4 | 544 | -8 | 10 | -4 | - | 7E-03 | 2.5 | L Caudate Head |

ALE, anatomic likelihood estimation; BA, Brodmann area; P, p-value; Z, peak z-value; R, right; L, left.

**Supplementary Table 4. Meta-analytic connectivity modeling results from primary outcomes (n = 17 ROIs), at cluster level inference p < 0.05 (FWE).**

| Cluster number | Volume (mm³) | MNI coordinates x | y | z | ALE | P | Z | Label (Side, region) |
|---|---|---|---|---|---|---|---|---|
| **1. Music perception** | | | | | | | | |
| *a. STG BA22 R: 824 foci, 43 experiments, 610 subjects (x=52, y=-20, z=4)* | | | | | | | | |
| 1 | 13592 | -54 | -18 | 2 | 6E-02 | 4E-20 | 9.1 | L Superior Temporal Gyrus BA22 |
| | | -58 | -28 | 6 | 5E-02 | 9E-15 | 7.7 | L Superior Temporal Gyrus BA22 |
| | | -42 | -30 | 8 | 5E-02 | 5E-13 | 7.1 | L Superior Temporal Gyrus BA41 |
| | | -48 | -36 | 14 | 4E-02 | 7E-11 | 6.4 | L Superior Temporal Gyrus BA41 |
| 2 | 10480 | 52 | -20 | 4 | 2E-01 | 0E+00 | 19.9 | R Superior Temporal Gyrus BA13 |
| | | 60 | -6 | 2 | 3E-02 | 7E-07 | 4.8 | R Superior Temporal Gyrus BA22 |
| 3 | 4312 | 34 | 20 | 2 | 4E-02 | 1E-10 | 6.3 | R Claustrum |
| | | 50 | 10 | 2 | 3E-02 | 8E-07 | 4.8 | R Insula BA13 |
| | | 46 | 8 | 2 | 3E-02 | 2E-06 | 4.6 | R Precentral Gyrus BA44 |
| | | 54 | 18 | -4 | 2E-02 | 2E-05 | 4.1 | R Inferior Frontal Gyrus |
| 4 | 4088 | -2 | 2 | 62 | 4E-02 | 1E-10 | 6.3 | L Medial Frontal Gyrus BA6 |
| | | 4 | 10 | 54 | 3E-02 | 3E-08 | 5.4 | R Superior Frontal Gyrus BA6 |
| | | 4 | 16 | 40 | 2E-02 | 1E-05 | 4.2 | R Cingulate Gyrus BA32 |
| | | -2 | 12 | 44 | 2E-02 | 2E-05 | 4.1 | L Medial Frontal Gyrus BA32 |
| 5 | 2368 | 50 | -6 | 38 | 3E-02 | 4E-08 | 5.4 | R Precentral Gyrus BA6 |
| | | 42 | 6 | 40 | 2E-02 | 1E-04 | 3.6 | R Middle Frontal Gyrus BA6 |
| 6 | 2056 | -54 | 12 | -4 | 3E-02 | 6E-09 | 5.7 | L Superior Temporal Gyrus BA22 |
| | | -52 | 2 | -8 | 2E-02 | 2E-04 | 3.6 | L Superior Temporal Gyrus BA22 |
| 7 | 1512 | -38 | 16 | 4 | 3E-02 | 1E-07 | 5.2 | L Insula BA13 |
| 8 | 1048 | -12 | -64 | -16 | 3E-02 | 8E-07 | 4.8 | L Cerebellum |
| | | -26 | -66 | -18 | 2E-02 | 3E-05 | 4.0 | L Cerebellum |
| *b. STG BA22 L: 1143 foci, 60 experiments, 850 subjects (x=-54, y=-16, z=2)* | | | | | | | | |
| 1 | 19976 | -54 | -16 | 2 | 3E-01 | 0E+00 | 24.2 | L Superior Temporal Gyrus BA22 |
| | | -62 | -30 | 8 | 6E-02 | 4E-15 | 7.8 | L Superior Temporal Gyrus BA42 |
| | | -60 | -34 | 12 | 6E-02 | 7E-15 | 7.7 | L Superior Temporal Gyrus BA22 |
| | | -44 | -34 | 14 | 4E-02 | 2E-08 | 5.5 | L Superior Temporal Gyrus BA41 |
| | | -52 | -40 | 20 | 3E-02 | 2E-06 | 4.6 | L Superior Temporal Gyrus BA13 |
| | | -50 | 2 | -8 | 3E-02 | 4E-06 | 4.5 | L Superior Temporal Gyrus BA22 |
| | | -50 | 10 | -6 | 3E-02 | 8E-06 | 4.3 | L Insula BA13 |
| | | -44 | 2 | 0 | 2E-02 | 2E-05 | 4.1 | L Insula BA13 |
| | | -54 | 6 | 8 | 2E-02 | 5E-05 | 3.9 | L Precentral Gyrus BA44 |
| 2 | 19720 | 58 | -14 | 0 | 1E-01 | 3E-36 | 12.5 | R Superior Temporal Gyrus BA22 |
| | | 64 | -32 | 8 | 5E-02 | 5E-14 | 7.4 | R Superior Temporal Gyrus BA42 |
| | | 54 | 12 | -12 | 3E-02 | 2E-06 | 4.6 | R Superior Temporal Gyrus BA22 |
| 3 | 1824 | 36 | 22 | 4 | 3E-02 | 4E-06 | 4.5 | R Insula BA13 |
| | | 36 | 22 | -8 | 2E-02 | 2E-05 | 4.1 | R Insula |
| *c. MedFG BA6 L: 1773 foci, 58 experiments, 838 subjects (x=-2, y=-2, z=66)* | | | | | | | | |
| 1 | 28752 | -2 | -2 | 64 | 3E-01 | 0E+00 | 22.8 | L Medial Frontal Gyrus BA6 |
| | | 52 | 2 | 50 | 5E-02 | 8E-12 | 6.7 | R Precentral Gyrus BA6 |
| | | 36 | 24 | 2 | 5E-02 | 6E-10 | 6.1 | R Insula BA13 |
| | | -6 | 12 | 42 | 4E-02 | 7E-09 | 5.7 | L Cingulate Gyrus BA32 |
| | | 56 | 8 | 38 | 4E-02 | 7E-09 | 5.7 | R Middle Frontal Gyrus BA6 |
| | | 8 | 12 | 38 | 4E-02 | 1E-08 | 5.6 | R Cingulate Gyrus BA32 |
| | | 42 | -4 | 54 | 4E-02 | 1E-07 | 5.1 | R Precentral Gyrus BA6 |
| | | 30 | -6 | 60 | 4E-02 | 2E-07 | 5.0 | R Middle Frontal Gyrus BA6 |
| | | 46 | 8 | 4 | 4E-02 | 3E-07 | 5.0 | R Insula BA13 |
| | | 54 | 14 | 14 | 3E-02 | 8E-06 | 4.3 | R Inferior Frontal Gyrus BA44 |
| | | 62 | 10 | 16 | 3E-02 | 8E-06 | 4.3 | R Inferior Frontal Gyrus BA44 |
| | | 52 | -8 | 38 | 3E-02 | 8E-05 | 3.8 | R Precentral Gyrus BA4 |
| | | 18 | -4 | 66 | 3E-02 | 2E-04 | 3.5 | R Superior Frontal Gyrus BA6 |
| | | 6 | 30 | 36 | 2E-02 | 5E-04 | 3.3 | R Cingulate Gyrus BA32 |
| 2 | 19104 | -50 | -8 | 50 | 5E-02 | 1E-11 | 6.7 | L Precentral Gyrus BA4 |
| | | -30 | 22 | 4 | 5E-02 | 7E-10 | 6.1 | L Insula BA13 |
| | | -54 | 2 | 38 | 5E-02 | 2E-09 | 5.9 | L Precentral Gyrus BA6 |
| | | -30 | -8 | 56 | 5E-02 | 3E-09 | 5.8 | L Precentral Gyrus BA6 |

| | | | | | | | | |
|---|---|---|---|---|---|---|---|---|
| | | -54 | 12 | 0 | 4E-02 | 5E-09 | 5.7 | L Superior Temporal Gyrus BA22 |
| | | -54 | 8 | 20 | 4E-02 | 2E-07 | 5.0 | L Inferior Frontal Gyrus BA44 |
| | | -38 | -26 | 54 | 3E-02 | 3E-05 | 4.0 | L Postcentral Gyrus BA3 |
| | | -48 | 12 | 28 | 3E-02 | 5E-05 | 3.9 | L Inferior Frontal Gyrus BA9 |
| | | -60 | -4 | 20 | 2E-02 | 3E-04 | 3.4 | L Precentral Gyrus BA4 |
| 3 | 8864 | -22 | 2 | 4 | 7E-02 | 1E-15 | 7.9 | L Lentiform Nucleus |
| | | -12 | -18 | 4 | 5E-02 | 7E-12 | 6.8 | L Thalamus |
| 4 | 8048 | 16 | -16 | 6 | 6E-02 | 8E-15 | 7.7 | R Thalamus |
| | | 22 | 6 | 4 | 6E-02 | 1E-13 | 7.3 | R Lentiform Nucleus |
| 5 | 5008 | 8 | -66 | -18 | 4E-02 | 1E-08 | 5.6 | R Declive |
| | | 18 | -60 | -20 | 4E-02 | 2E-08 | 5.5 | R Culmen |
| | | 32 | -62 | -26 | 4E-02 | 1E-07 | 5.2 | R Culmen |
| | | 38 | -68 | -24 | 3E-02 | 2E-06 | 4.6 | R Declive |
| | | 24 | -74 | -20 | 3E-02 | 1E-04 | 3.7 | R Declive |
| 6 | 4584 | 66 | -32 | 14 | 4E-02 | 8E-08 | 5.2 | R Superior Temporal Gyrus BA22 |
| | | 58 | -14 | 4 | 4E-02 | 3E-07 | 5.0 | R Superior Temporal Gyrus BA22 |
| | | 56 | -22 | 4 | 3E-02 | 4E-06 | 4.4 | R Superior Temporal Gyrus BA41 |
| | | 66 | -22 | 4 | 3E-02 | 1E-05 | 4.2 | R Superior Temporal Gyrus BA41 |
| | | 60 | -6 | 4 | 3E-02 | 5E-05 | 3.9 | R Superior Temporal Gyrus BA22 |
| 7 | 4056 | -28 | -60 | -26 | 5E-02 | 4E-12 | 6.8 | L Culmen |
| | | -12 | -52 | -22 | 3E-02 | 5E-06 | 4.4 | L Dentate |
| | | -12 | -64 | -18 | 3E-02 | 7E-06 | 4.3 | L Declive |
| | | -16 | -64 | -18 | 3E-02 | 9E-06 | 4.3 | L Declive |
| 8 | 2360 | -30 | -54 | 54 | 4E-02 | 3E-07 | 5.0 | L Superior Parietal Lobule BA7 |
| | | -22 | -60 | 58 | 3E-02 | 3E-06 | 4.5 | L Precuneus BA7 |
| 9 | 2320 | -60 | -32 | 8 | 3E-02 | 2E-06 | 4.6 | L Superior Temporal Gyrus BA42 |
| | | -64 | -34 | 10 | 3E-02 | 2E-06 | 4.6 | L Superior Temporal Gyrus BA22 |
| | | -52 | -42 | 20 | 3E-02 | 1E-05 | 4.2 | L Superior Temporal Gyrus BA13 |
| | | -58 | -22 | 16 | 3E-02 | 2E-04 | 3.6 | L Postcentral Gyrus BA40 |
| *d. PUT R: 1821 foci, 70 experiments, 1052 subjects (x=22, y=8, z=6)* | | | | | | | | |
| 1 | 34416 | -22 | 4 | 6 | 1E-01 | 3E-28 | 11.0 | L Lentiform Nucleus |
| | | -30 | 20 | 2 | 8E-02 | 3E-18 | 8.6 | L Claustrum |
| | | -12 | -14 | 6 | 7E-02 | 7E-16 | 8.0 | L Thalamus |
| | | -52 | 8 | 18 | 5E-02 | 2E-10 | 6.3 | L Inferior Frontal Gyrus BA44 |
| | | -30 | -8 | 56 | 5E-02 | 3E-09 | 5.8 | L Precentral Gyrus BA6 |
| | | -46 | 4 | 4 | 5E-02 | 7E-09 | 5.7 | L Precentral Gyrus BA44 |
| | | -44 | 6 | 26 | 4E-02 | 4E-08 | 5.4 | L Inferior Frontal Gyrus BA9 |
| | | -46 | -6 | 54 | 4E-02 | 4E-08 | 5.4 | L Precentral Gyrus BA4 |
| | | -54 | 6 | 34 | 4E-02 | 1E-07 | 5.2 | L Precentral Gyrus BA6 |
| | | -48 | -2 | 38 | 3E-02 | 5E-06 | 4.4 | L Precentral Gyrus BA6 |
| | | -48 | -10 | 44 | 3E-02 | 1E-05 | 4.2 | L Precentral Gyrus BA4 |
| | | -30 | -14 | -2 | 2E-02 | 7E-04 | 3.2 | L Lentiform Nucleus |
| 2 | 29528 | 22 | 8 | 6 | 3E-01 | 0E+00 | 24.2 | R Lentiform Nucleus |
| | | 36 | 20 | 2 | 6E-02 | 9E-12 | 6.7 | R Insula BA13 |
| | | 14 | -12 | 6 | 5E-02 | 3E-10 | 6.2 | R Thalamus |
| | | 54 | 2 | 46 | 5E-02 | 8E-10 | 6.0 | R Precentral Gyrus BA6 |
| | | 58 | 8 | 24 | 4E-02 | 1E-08 | 5.6 | R Inferior Frontal Gyrus BA9 |
| | | 54 | 6 | 12 | 4E-02 | 2E-07 | 5.1 | R Inferior Frontal Gyrus BA44 |
| | | 14 | -16 | 14 | 4E-02 | 8E-07 | 4.8 | R Thalamus |
| | | 48 | 12 | 2 | 3E-02 | 8E-06 | 4.3 | R Precentral Gyrus BA44 |
| | | 46 | 8 | -4 | 3E-02 | 3E-05 | 4.0 | R Insula |
| 3 | 11672 | -2 | 0 | 62 | 7E-02 | 8E-15 | 7.7 | L Medial Frontal Gyrus BA6 |
| | | -4 | 2 | 58 | 6E-02 | 4E-14 | 7.5 | L Medial Frontal Gyrus BA6 |
| | | 8 | 2 | 60 | 6E-02 | 1E-13 | 7.3 | R Medial Frontal Gyrus BA6 |
| | | -2 | 6 | 50 | 6E-02 | 2E-12 | 7.0 | L Medial Frontal Gyrus BA6 |
| | | 8 | 14 | 40 | 4E-02 | 3E-07 | 5.0 | R Cingulate Gyrus BA32 |
| 4 | 3472 | 42 | -44 | 46 | 4E-02 | 3E-08 | 5.4 | R Inferior Parietal Lobule BA40 |
| | | 44 | -38 | 46 | 4E-02 | 3E-07 | 5.0 | R Inferior Parietal Lobule BA40 |
| | | 58 | -30 | 42 | 3E-02 | 4E-05 | 3.9 | R Inferior Parietal Lobule BA40 |
| | | 40 | -40 | 58 | 3E-02 | 1E-04 | 3.6 | R Inferior Parietal Lobule BA40 |
| 5 | 1784 | 36 | 42 | 24 | 4E-02 | 2E-07 | 5.1 | R Middle Frontal Gyrus BA9 |
| *e. PUT L: 1493 foci, 69 experiments, 1139 subjects (x=-22, y=4, z=6)* | | | | | | | | |
| 1 | 31472 | -22 | 4 | 6 | 3E-01 | 0E+00 | 25.1 | L Lentiform Nucleus |
| | | -34 | 24 | 0 | 6E-02 | 2E-13 | 7.3 | L Insula BA13 |
| | | -12 | -16 | 6 | 6E-02 | 2E-13 | 7.2 | L Thalamus |
| | | -52 | -4 | 46 | 4E-02 | 2E-08 | 5.5 | L Precentral Gyrus BA4 |

| | | | | | | | | |
|---|---|---|---|---|---|---|---|---|
| | | -50 | 6 | 32 | 4E-02 | 3E-08 | 5.4 | L Precentral Gyrus BA6 |
| | | -54 | 6 | 6 | 4E-02 | 6E-08 | 5.3 | L Superior Temporal Gyrus BA22 |
| | | -54 | 12 | 2 | 4E-02 | 1E-07 | 5.1 | L Precentral Gyrus BA44 |
| | | -52 | 10 | 18 | 3E-02 | 9E-07 | 4.8 | L Inferior Frontal Gyrus BA44 |
| | | -40 | 34 | 12 | 2E-02 | 2E-04 | 3.5 | L Middle Frontal Gyrus BA46 |
| | | -30 | -12 | -2 | 2E-02 | 2E-04 | 3.5 | L Lentiform Nucleus |
| | | -10 | -18 | -10 | 2E-02 | 5E-04 | 3.3 | L Brainstem |
| 2 | 17664 | 22 | 6 | 6 | 1E-01 | 8E-32 | 11.7 | R Lentiform Nucleus |
| | | 34 | 18 | 4 | 6E-02 | 2E-14 | 7.6 | R Claustrum |
| | | 14 | -14 | 4 | 5E-02 | 5E-10 | 6.1 | R Thalamus |
| | | 34 | 26 | -8 | 3E-02 | 2E-06 | 4.6 | R Insula BA13 |
| | | 50 | 14 | 0 | 3E-02 | 3E-05 | 4.0 | R Insula BA13 |
| | | 54 | 8 | 6 | 3E-02 | 1E-04 | 3.7 | R Precentral Gyrus BA44 |
| 3 | 11768 | -4 | 0 | 62 | 7E-02 | 2E-18 | 8.7 | L Medial Frontal Gyrus BA6 |
| | | 8 | 16 | 42 | 5E-02 | 5E-10 | 6.1 | R Cingulate Gyrus BA32 |
| | | 8 | 4 | 60 | 4E-02 | 1E-07 | 5.1 | R Medial Frontal Gyrus BA6 |
| | | -8 | 12 | 38 | 3E-02 | 1E-05 | 4.2 | L Cingulate Gyrus BA32 |
| | | 4 | 16 | 52 | 3E-02 | 2E-05 | 4.1 | R Superior Frontal Gyrus BA6 |
| | | 8 | 28 | 38 | 2E-02 | 4E-04 | 3.4 | R Cingulate Gyrus BA32 |
| 4 | 3632 | 54 | -32 | 4 | 4E-02 | 4E-07 | 4.9 | R Superior Temporal Gyrus BA22 |
| | | 54 | -26 | 0 | 3E-02 | 8E-07 | 4.8 | R Superior Temporal Gyrus BA22 |
| | | 66 | -22 | 6 | 3E-02 | 5E-06 | 4.4 | R Superior Temporal Gyrus BA42 |
| | | 68 | -32 | 14 | 2E-02 | 4E-04 | 3.4 | R Superior Temporal Gyrus BA22 |
| 5 | 3624 | 54 | 2 | 46 | 5E-02 | 1E-10 | 6.3 | R Precentral Gyrus BA6 |
| | | 48 | 10 | 28 | 3E-02 | 3E-06 | 4.5 | R Inferior Frontal Gyrus BA9 |
| | | 56 | 10 | 34 | 3E-02 | 4E-05 | 3.9 | R Middle Frontal Gyrus BA9 |
| 6 | 2920 | -28 | -60 | -26 | 6E-02 | 8E-13 | 7.1 | L Culmen |
| 7 | 2768 | -52 | -42 | 18 | 3E-02 | 6E-06 | 4.4 | L Superior Temporal Gyrus BA13 |
| | | -62 | -30 | 8 | 3E-02 | 3E-05 | 4.0 | L Superior Temporal Gyrus BA42 |
| | | -60 | -16 | 6 | 3E-02 | 5E-05 | 3.9 | L Superior Temporal Gyrus BA41 |
| | | -60 | -36 | 14 | 3E-02 | 5E-05 | 3.9 | L Superior Temporal Gyrus BA22 |
| | | -54 | -40 | 12 | 3E-02 | 6E-05 | 3.8 | L Superior Temporal Gyrus BA22 |
| | | -52 | -16 | 2 | 3E-02 | 1E-04 | 3.7 | L Superior Temporal Gyrus BA22 |
| *f. CRBL L: 1201 foci, 47 experiments, 688 subjects (x=-28, y=-64, z=-26)* | | | | | | | | |
| 1 | 11472 | 36 | 18 | 6 | 4E-02 | 3E-09 | 5.8 | R Insula BA13 |
| | | 42 | 24 | -2 | 4E-02 | 4E-09 | 5.8 | R Insula BA13 |
| | | 48 | 18 | -2 | 4E-02 | 7E-08 | 5.3 | R Insula BA13 |
| | | 56 | 4 | 34 | 3E-02 | 1E-07 | 5.2 | R Precentral Gyrus BA6 |
| | | 54 | 0 | 46 | 3E-02 | 2E-07 | 5.0 | R Precentral Gyrus BA4 |
| | | 32 | 24 | -8 | 3E-02 | 5E-07 | 4.9 | R Claustrum |
| | | 52 | 12 | 14 | 3E-02 | 2E-06 | 4.6 | R Insula BA13 |
| | | 50 | 10 | 22 | 3E-02 | 3E-05 | 4.0 | R Inferior Frontal Gyrus BA9 |
| | | 38 | 10 | -8 | 2E-02 | 8E-05 | 3.8 | R Claustrum |
| | | 52 | -6 | 36 | 2E-02 | 4E-04 | 3.4 | R Precentral Gyrus BA6 |
| 2 | 10608 | 30 | -62 | -28 | 8E-02 | 5E-22 | 9.6 | R Culmen |
| | | 10 | -74 | -20 | 4E-02 | 5E-09 | 5.7 | R Declive |
| | | -6 | -74 | -28 | 3E-02 | 8E-07 | 4.8 | L Uvula |
| | | 2 | -58 | -22 | 3E-02 | 2E-05 | 4.1 | R.Culmen |
| 3 | 9624 | 0 | 14 | 44 | 4E-02 | 1E-10 | 6.3 | L Medial Frontal Gyrus BA32 |
| | | 8 | 16 | 44 | 4E-02 | 6E-10 | 6.1 | R Medial Frontal Gyrus BA32 |
| | | -4 | 2 | 60 | 4E-02 | 2E-09 | 5.9 | L Medial Frontal Gyrus BA6 |
| | | 6 | 10 | 56 | 4E-02 | 3E-09 | 5.8 | R Medial Frontal Gyrus BA6 |
| 4 | 9448 | -34 | 22 | 2 | 5E-02 | 3E-13 | 7.2 | L Insula BA13 |
| | | -22 | 4 | 6 | 5E-02 | 6E-13 | 7.1 | L Lentiform Nucleus |
| | | -12 | -14 | 4 | 5E-02 | 6E-11 | 6.4 | L Thalamus |
| 5 | 7488 | -58 | -16 | 6 | 4E-02 | 3E-08 | 5.4 | L Superior Temporal Gyrus BA41 |
| | | -42 | -34 | 12 | 3E-02 | 1E-07 | 5.2 | L Transverse Temporal Gyrus BA41 |
| | | -54 | -40 | 12 | 3E-02 | 2E-07 | 5.1 | L Superior Temporal Gyrus BA22 |
| | | -62 | -34 | 12 | 3E-02 | 3E-07 | 5.0 | L Superior Temporal Gyrus BA22 |
| | | -60 | -36 | 24 | 3E-02 | 2E-06 | 4.6 | L Insula BA13 |
| 6 | 6056 | -28 | -62 | -26 | 2E-01 | 0E+00 | 19.7 | L Culmen |
| 7 | 3568 | 58 | -14 | 2 | 3E-02 | 8E-07 | 4.8 | R Superior Temporal Gyrus BA22 |
| | | 58 | -32 | 4 | 3E-02 | 6E-06 | 4.4 | R Middle Temporal Gyrus BA22 |
| | | 56 | -28 | 0 | 3E-02 | 6E-06 | 4.4 | R Superior Temporal Gyrus |
| | | 54 | -24 | 6 | 3E-02 | 2E-05 | 4.1 | R Superior Temporal Gyrus BA41 |
| | | 66 | -22 | 6 | 2E-02 | 6E-05 | 3.8 | R Superior Temporal Gyrus BA42 |

| | | | | | | | | |
|---|---|---|---|---|---|---|---|---|
| | | 68 | -32 | 12 | 2E-02 | 1E-04 | 3.7 | R Superior Temporal Gyrus BA22 |
| 8 | 2256 | -54 | 10 | 2 | 4E-02 | 8E-09 | 5.6 | L Precentral Gyrus BA44 |
| 9 | 2152 | 22 | 8 | 6 | 4E-02 | 3E-10 | 6.2 | R Lentiform Nucleus |
| 10 | 2024 | 12 | -14 | 6 | 4E-02 | 2E-08 | 5.5 | R Thalamus |
| | | 6 | -24 | -2 | 2E-02 | 8E-05 | 3.8 | R Thalamus |
| *g. INS L: 1787 foci, 87 experiments, 1332 subjects (x=-32, y=18, z=10)* | | | | | | | | |
| 1 | 45704 | -32 | 18 | 8 | 4E-01 | 0E+00 | 29.1 | L Insula BA13 |
| | | 36 | 22 | 6 | 1E-01 | 1E-42 | 13.6 | R Insula BA13 |
| | | -10 | -16 | 8 | 9E-02 | 1E-24 | 10.2 | L Thalamus |
| | | -52 | 8 | 22 | 6E-02 | 6E-13 | 7.1 | L Inferior Frontal Gyrus BA9 |
| | | 10 | -16 | 6 | 5E-02 | 2E-11 | 6.6 | R Thalamus |
| | | 18 | 10 | 4 | 5E-02 | 1E-09 | 6.0 | R Caudate |
| | | -18 | 6 | 6 | 4E-02 | 1E-08 | 5.5 | L Lentiform Nucleus |
| | | -44 | 4 | -2 | 4E-02 | 1E-07 | 5.2 | L Insula BA13 |
| | | -46 | 0 | 4 | 4E-02 | 2E-07 | 5.1 | L Insula BA13 |
| | | -38 | -4 | 14 | 4E-02 | 3E-07 | 5.0 | L Insula BA13 |
| | | 8 | -22 | -10 | 3E-02 | 2E-04 | 3.5 | R Brainstem |
| 2 | 12360 | -2 | 10 | 50 | 7E-02 | 2E-15 | 7.8 | L Medial Frontal Gyrus BA6 |
| | | 10 | 16 | 42 | 4E-02 | 2E-08 | 5.5 | R Medial Frontal Gyrus BA32 |
| | | -6 | 16 | 36 | 4E-02 | 8E-07 | 4.8 | L Cingulate Gyrus BA32 |
| | | 0 | 30 | 32 | 3E-02 | 6E-05 | 3.9 | L Cingulate Gyrus BA32 |
| | | -2 | 26 | 44 | 3E-02 | 3E-04 | 3.4 | L Medial Frontal Gyrus BA8 |
| 3 | 6064 | 50 | 10 | 28 | 9E-02 | 1E-21 | 9.5 | R Inferior Frontal Gyrus BA9 |
| | | 56 | 16 | 16 | 3E-02 | 1E-04 | 3.7 | R Inferior Frontal Gyrus BA44 |
| 4 | 3864 | -34 | -48 | 44 | 4E-02 | 6E-07 | 4.8 | L Inferior Parietal Lobule BA40 |
| | | -44 | -38 | 50 | 4E-02 | 1E-06 | 4.7 | L Inferior Parietal Lobule BA40 |
| | | -30 | -56 | 52 | 3E-02 | 3E-05 | 4.0 | L Superior Parietal Lobule BA7 |
| | | -24 | -66 | 48 | 3E-02 | 4E-05 | 4.0 | L Superior Parietal Lobule BA7 |
| 5 | 3360 | -30 | -8 | 56 | 5E-02 | 1E-09 | 6.0 | L Precentral Gyrus BA6 |
| | | -42 | -4 | 48 | 3E-02 | 2E-04 | 3.5 | L Precentral Gyrus BA6 |
| 6 | 3200 | 44 | 42 | 18 | 4E-02 | 2E-06 | 4.7 | R Middle Frontal Gyrus BA10 |
| | | 42 | 38 | 28 | 3E-02 | 4E-06 | 4.5 | R Middle Frontal Gyrus BA9 |
| | | 40 | 50 | 12 | 3E-02 | 1E-04 | 3.7 | R Middle Frontal Gyrus BA10 |
| 7 | 2744 | -34 | 42 | 26 | 4E-02 | 2E-06 | 4.6 | L Superior Frontal Gyrus BA9 |
| | | -42 | 38 | 12 | 3E-02 | 4E-06 | 4.5 | L Middle Frontal Gyrus BA46 |
| | | -34 | 50 | 12 | 3E-02 | 5E-05 | 3.9 | L Middle Frontal Gyrus BA10 |
| | | -42 | 32 | 26 | 3E-02 | 1E-04 | 3.6 | L Middle Frontal Gyrus BA9 |
| 8 | 2416 | 34 | 2 | 62 | 4E-02 | 1E-08 | 5.6 | R Middle Frontal Gyrus BA6 |
| | | 34 | 2 | 50 | 3E-02 | 9E-06 | 4.3 | R Middle Frontal Gyrus BA6 |
| *h. PreCG R: 1613 foci, 63 experiments, 931 subjects (x=54, y=0, z=46)* | | | | | | | | |
| 1 | 20416 | 54 | 0 | 46 | 3E-01 | 0E+00 | 23.8 | R Precentral Gyrus BA4 |
| | | 34 | 24 | 4 | 5E-02 | 2E-10 | 6.3 | R Insula BA13 |
| | | 38 | 22 | -2 | 5E-02 | 1E-09 | 6.0 | R Insula BA13 |
| | | 44 | 14 | 6 | 4E-02 | 2E-08 | 5.5 | R Insula BA13 |
| | | 50 | 14 | 2 | 4E-02 | 5E-08 | 5.3 | R Precentral Gyrus BA44 |
| | | 36 | -4 | 58 | 4E-02 | 6E-08 | 5.3 | R Precentral Gyrus BA6 |
| | | 52 | 10 | 22 | 4E-02 | 1E-07 | 5.1 | R Inferior Frontal Gyrus BA9 |
| | | 52 | 6 | 28 | 4E-02 | 3E-07 | 5.0 | R Precentral Gyrus BA6 |
| | | 60 | 6 | 12 | 3E-02 | 3E-06 | 4.5 | R Precentral Gyrus BA6 |
| | | 58 | 8 | -6 | 3E-02 | 7E-05 | 3.8 | R Superior Temporal Gyrus BA22 |
| 2 | 17984 | -50 | -6 | 48 | 9E-02 | 6E-24 | 10.0 | L Precentral Gyrus BA4 |
| | | -34 | 20 | 4 | 5E-02 | 2E-11 | 6.6 | L Insula BA13 |
| | | -54 | 8 | 24 | 5E-02 | 2E-09 | 5.9 | L Inferior Frontal Gyrus BA9 |
| | | -58 | 0 | 16 | 4E-02 | 9E-09 | 5.6 | L Precentral Gyrus BA4 |
| | | -56 | 6 | 6 | 4E-02 | 2E-08 | 5.5 | L Superior Temporal Gyrus BA22 |
| | | -50 | 12 | 0 | 4E-02 | 7E-08 | 5.3 | L Insula BA13 |
| 3 | 14664 | -2 | 0 | 60 | 9E-02 | 3E-23 | 9.9 | L Medial Frontal Gyrus BA6 |
| | | -2 | 20 | 38 | 4E-02 | 2E-08 | 5.5 | L Cingulate Gyrus BA32 |
| | | 8 | 14 | 38 | 4E-02 | 5E-07 | 4.9 | R Cingulate Gyrus BA32 |
| 4 | 6080 | -54 | -18 | 0 | 4E-02 | 1E-08 | 5.6 | L Superior Temporal Gyrus BA22 |
| | | -54 | -40 | 22 | 4E-02 | 6E-08 | 5.3 | L Superior Temporal Gyrus BA13 |
| | | -62 | -30 | 8 | 4E-02 | 7E-08 | 5.3 | L Superior Temporal Gyrus BA42 |
| 5 | 5912 | 56 | -32 | 4 | 4E-02 | 3E-08 | 5.4 | R Superior Temporal Gyrus BA22 |
| | | 64 | -34 | 26 | 4E-02 | 5E-07 | 4.9 | R Inferior Parietal Lobule BA40 |
| | | 60 | -18 | 4 | 3E-02 | 3E-06 | 4.5 | R Superior Temporal Gyrus BA41 |
| 6 | 5056 | 12 | -14 | 6 | 4E-02 | 3E-09 | 5.8 | R Thalamus |

| | | | | | | | | |
|---|---|---|---|---|---|---|---|---|
| | | 24 | 4 | 4 | 4E-02 | 2E-08 | 5.5 | R Putamen |
| | | 24 | 0 | 6 | 4E-02 | 1E-07 | 5.1 | R Putamen |
| 7 | 1984 | -22 | 4 | 6 | 5E-02 | 2E-09 | 5.9 | L Putamen |
| | | -28 | -6 | -4 | 3E-02 | 5E-05 | 3.9 | L Putamen |
| | | -22 | -2 | 16 | 2E-02 | 3E-04 | 3.5 | L Putamen |
| **2. Music production** | | | | | | | | |
| *a. STG BA42 R: 753 foci, 32 experiments, 455 subjects (x=66, y=-24, z=10)* | | | | | | | | |
| 1 | 7648 | 64 | -24 | 10 | 1E-01 | 0E+00 | 17.2 | R Superior Temporal Gyrus BA41 |
| | | 54 | -16 | 4 | 2E-02 | 9E-05 | 3.7 | R Superior Temporal Gyrus BA22 |
| | | 62 | -40 | 2 | 2E-02 | 2E-04 | 3.6 | R Middle Temporal Gyrus BA22 |
| 2 | 7224 | -60 | -24 | 10 | 5E-02 | 3E-15 | 7.8 | L Superior Temporal Gyrus BA41 |
| | | -42 | -34 | 14 | 2E-02 | 1E-04 | 3.7 | L Transverse Temporal Gyrus BA41 |
| | | -42 | -26 | 12 | 1E-02 | 9E-04 | 3.1 | L Transverse Temporal Gyrus BA41 |
| 3 | 3192 | -4 | 6 | 52 | 3E-02 | 1E-08 | 5.6 | L Medial Frontal Gyrus BA6 |
| | | -4 | -2 | 62 | 2E-02 | 1E-06 | 4.7 | L Medial Frontal Gyrus BA6 |
| | | 2 | 0 | 70 | 2E-02 | 5E-05 | 3.9 | L Medial Frontal Gyrus BA6 |
| | | 2 | 16 | 48 | 2E-02 | 7E-05 | 3.8 | L Medial Frontal Gyrus BA32 |
| 4 | 2736 | -34 | 24 | 0 | 3E-02 | 7E-07 | 4.8 | L Insula BA13 |
| | | -52 | 8 | -2 | 3E-02 | 1E-06 | 4.7 | L Superior Temporal Gyrus BA22 |
| | | -48 | 14 | 0 | 2E-02 | 6E-06 | 4.4 | L Insula BA13 |
| | | -52 | 0 | 4 | 2E-02 | 6E-04 | 3.2 | L Precentral Gyrus BA44 |
| 5 | 2624 | 52 | 4 | -8 | 3E-02 | 2E-08 | 5.5 | R Superior Temporal Gyrus BA22 |
| | | 48 | 20 | -6 | 2E-02 | 4E-05 | 4.0 | R Insula |
| 6 | 1672 | -12 | -22 | -6 | 2E-02 | 6E-06 | 4.4 | L Brainstem |
| | | -12 | -18 | 0 | 2E-02 | 7E-06 | 4.3 | L Thalamus |
| | | -10 | -12 | 4 | 2E-02 | 6E-05 | 3.8 | L Thalamus |
| 7 | 1608 | 54 | 0 | 44 | 3E-02 | 5E-07 | 4.9 | R Precentral Gyrus BA6 |
| | | 54 | 8 | 30 | 2E-02 | 7E-05 | 3.8 | R Inferior Frontal Gyrus BA9 |
| *b. PreCG L: 1372 foci, 79 experiments, 1034 subjects (x=-48, y=0, z=42)* | | | | | | | | |
| 1 | 25824 | -48 | 0 | 42 | 3E-01 | 0E+00 | 26.8 | L Precentral Gyrus BA6 |
| | | -32 | 18 | 2 | 6E-02 | 1E-13 | 7.4 | L Claustrum |
| | | -56 | 8 | 18 | 5E-02 | 2E-12 | 6.9 | L Inferior Frontal Gyrus BA44 |
| | | -46 | 16 | 22 | 5E-02 | 3E-12 | 6.9 | L Inferior Frontal Gyrus BA9 |
| | | -22 | -8 | 58 | 4E-02 | 5E-09 | 5.7 | L Middle Frontal Gyrus BA6 |
| | | -32 | -6 | 52 | 2E-02 | 7E-05 | 3.8 | L Precentral Gyrus BA6 |
| | | -42 | 28 | -2 | 2E-02 | 8E-05 | 3.8 | L Inferior Frontal Gyrus BA47 |
| | | -56 | -6 | -6 | 2E-02 | 1E-04 | 3.7 | L Superior Temporal Gyrus BA22 |
| 2 | 13904 | -2 | 10 | 50 | 9E-02 | 2E-24 | 10.1 | L Medial Frontal Gyrus BA6 |
| | | 0 | 6 | 62 | 7E-02 | 9E-18 | 8.5 | L Medial Frontal Gyrus BA6 |
| 3 | 8792 | 52 | 2 | 36 | 5E-02 | 2E-10 | 6.3 | R Precentral Gyrus BA6 |
| | | 50 | 10 | 34 | 4E-02 | 9E-10 | 6.0 | R Precentral Gyrus BA9 |
| | | 38 | 0 | 54 | 4E-02 | 1E-07 | 5.1 | R Middle Frontal Gyrus BA6 |
| | | 42 | 16 | 24 | 3E-02 | 8E-06 | 4.3 | R Middle Frontal Gyrus BA9 |
| 4 | 7520 | -36 | -52 | 44 | 5E-02 | 2E-11 | 6.6 | L Inferior Parietal Lobule BA40 |
| | | -42 | -38 | 42 | 3E-02 | 3E-06 | 4.5 | L Inferior Parietal Lobule BA40 |
| | | -26 | -64 | 44 | 3E-02 | 2E-05 | 4.1 | L Precuneus BA7 |
| 5 | 3016 | -60 | -36 | 4 | 5E-02 | 1E-10 | 6.3 | L Middle Temporal Gyrus BA22 |
| | | -60 | -26 | 4 | 3E-02 | 1E-06 | 4.7 | L Superior Temporal Gyrus BA22 |
| | | -54 | -38 | 14 | 2E-02 | 1E-04 | 3.6 | L Superior Temporal Gyrus BA41 |
| 6 | 2760 | -40 | -76 | -10 | 4E-02 | 1E-08 | 5.6 | L Fusiform Gyrus BA19 |
| | | -38 | -70 | -12 | 3E-02 | 6E-07 | 4.9 | L.Declive |
| | | -42 | -60 | -8 | 3E-02 | 7E-06 | 4.4 | L Fusiform Gyrus BA37 |
| 7 | 2608 | 28 | -60 | 54 | 3E-02 | 3E-06 | 4.6 | R Superior Parietal Lobule BA7 |
| | | 36 | -54 | 50 | 3E-02 | 5E-06 | 4.4 | R Inferior Parietal Lobule BA7 |
| | | 40 | -44 | 44 | 3E-02 | 7E-06 | 4.3 | R Inferior Parietal Lobule BA40 |
| 8 | 2208 | 36 | -62 | -28 | 3E-02 | 2E-06 | 4.6 | R Culmen |
| | | 30 | -72 | -18 | 3E-02 | 1E-05 | 4.3 | R Declive |
| 9 | 1880 | 36 | 20 | 0 | 4E-02 | 6E-08 | 5.3 | R Claustrum |
| | | 48 | 16 | 2 | 2E-02 | 9E-05 | 3.8 | R Precentral Gyrus BA44 |
| *c. MedFG BA6 L: 403 foci, 21 experiments, 261 subjects (x=-10, y=-10, z=52)* | | | | | | | | |
| 1 | 11352 | -36 | -24 | 58 | 4E-02 | 2E-14 | 8E+00 | L Precentral Gyrus BA4 |
| | | -36 | -34 | 46 | 3E-02 | 2E-08 | 6E+00 | L Inferior Parietal Lobule BA40 |
| | | -48 | -18 | 52 | 2E-02 | 1E-07 | 5E+00 | L Postcentral Gyrus BA2 |
| 2 | 6112 | -8 | -8 | 52 | 1E-01 | 0E+00 | 1E+01 | L Medial Frontal Gyrus BA6 |
| | | 8 | -12 | 50 | 2E-02 | 6E-06 | 4E+00 | R Paracentral Lobule BA31 |

| | | | | | | | | |
|---|---|---|---|---|---|---|---|---|
| 3 | 2280 | 22 | -52 | -24 | 2E-02 | 4E-08 | 5E+00 | R Culmen |
| | | 14 | -56 | -14 | 1E-02 | 5E-04 | 3E+00 | R Culmen |
| 4 | 1952 | -56 | -24 | 22 | 2E-02 | 1E-06 | 5E+00 | L Postcentral Gyrus BA40 |
| | | -46 | -28 | 20 | 2E-02 | 3E-06 | 5E+00 | L Insula BA13 |
| | | -52 | -28 | 22 | 2E-02 | 4E-06 | 4E+00 | L Insula BA13 |
| 5 | 1816 | -38 | -2 | 4 | 2E-02 | 1E-06 | 5E+00 | L Claustrum |
| | | -44 | -4 | 4 | 2E-02 | 2E-05 | 4E+00 | L Insula BA13 |
| | | -30 | -14 | 6 | 1E-02 | 1E-04 | 4E+00 | L Lentiform Nucleus |
| | | -30 | -8 | 6 | 1E-02 | 2E-04 | 4E+00 | L Lentiform Nucleus |
| 6 | 1224 | -16 | -22 | 6 | 2E-02 | 2E-07 | 5E+00 | L Thalamus |
| *d. STG BA41 L: 529 foci, 38 experiments, 522 subjects (x=-42, y=-28, z=6)* | | | | | | | | |
| 1 | 13576 | 50 | -26 | 6 | 5E-02 | 2E-17 | 8.4 | R Superior Temporal Gyrus BA41 |
| | | 50 | -18 | 4 | 5E-02 | 4E-17 | 8.3 | R Superior Temporal Gyrus BA13 |
| | | 66 | -30 | 4 | 4E-02 | 4E-12 | 6.9 | R Middle Temporal Gyrus BA22 |
| 2 | 11104 | -42 | -28 | 6 | 2E-01 | 0E+00 | 17.6 | L Superior Temporal Gyrus BA13 |
| | | -50 | -14 | 0 | 3E-02 | 1E-08 | 5.6 | L Superior Temporal Gyrus BA22 |
| | | -62 | -30 | 12 | 3E-02 | 8E-08 | 5.2 | L Superior Temporal Gyrus BA42 |
| | | -60 | -24 | 6 | 3E-02 | 1E-07 | 5.2 | L Superior Temporal Gyrus BA41 |
| 3 | 1248 | -28 | -10 | -4 | 3E-02 | 2E-08 | 5.5 | L Putamen |
| *e. PreCG R: 1105 foci, 51 experiments, 846 subjects (x=54, y=4, z=32)* | | | | | | | | |
| 1 | 11192 | -2 | 10 | 48 | 5E-02 | 1E-13 | 7.3 | L Cingulate Gyrus BA24 |
| | | 6 | 16 | 42 | 5E-02 | 7E-13 | 7.1 | R Cingulate Gyrus BA32 |
| | | 4 | 8 | 50 | 5E-02 | 6E-11 | 6.4 | R Medial Frontal Gyrus BA6 |
| | | -2 | -2 | 62 | 4E-02 | 1E-08 | 5.5 | L Medial Frontal Gyrus BA6 |
| 2 | 9016 | 54 | 4 | 32 | 2E-01 | 0E+00 | 20.8 | R Precentral Gyrus BA6 |
| | | 56 | 10 | 12 | 3E-02 | 2E-06 | 4.6 | R Inferior Frontal Gyrus BA44 |
| 3 | 8128 | -50 | 6 | 28 | 5E-02 | 5E-13 | 7.1 | L Precentral Gyrus BA6 |
| | | -50 | 2 | 42 | 5E-02 | 5E-11 | 6.5 | L Precentral Gyrus BA6 |
| 4 | 2792 | -28 | -62 | -26 | 3E-02 | 2E-06 | 4.7 | L Culmen |
| | | -42 | -58 | -18 | 3E-02 | 3E-06 | 4.5 | L Fusiform Gyrus BA37 |
| | | -14 | -62 | -18 | 3E-02 | 7E-06 | 4.4 | L Declive |
| 5 | 2600 | -28 | -52 | 54 | 3E-02 | 1E-07 | 5.1 | L Superior Parietal Lobule BA7 |
| | | -24 | -60 | 54 | 3E-02 | 1E-06 | 4.8 | L Superior Parietal Lobule BA7 |
| 6 | 2480 | -58 | -34 | 28 | 3E-02 | 6E-06 | 4.4 | L Inferior Parietal Lobule BA40 |
| | | -50 | -28 | 38 | 3E-02 | 2E-05 | 4.1 | L Postcentral Gyrus BA2 |
| | | -38 | -28 | 52 | 2E-02 | 4E-05 | 4.0 | L Postcentral Gyrus BA3 |
| | | -46 | -30 | 24 | 2E-02 | 9E-05 | 3.7 | L Insula BA13 |
| 7 | 1816 | 36 | 22 | 4 | 4E-02 | 3E-09 | 5.8 | R Insula BA13 |
| 8 | 1816 | 30 | -56 | 50 | 3E-02 | 1E-07 | 5.1 | R Superior Parietal Lobule BA7 |
| | | 24 | -60 | 60 | 2E-02 | 3E-04 | 3.4 | R Superior Parietal Lobule BA7 |
| 9 | 1656 | 60 | -32 | 24 | 3E-02 | 1E-06 | 4.7 | R Inferior Parietal Lobule BA40 |
| | | 50 | -32 | 30 | 2E-02 | 4E-05 | 3.9 | R Inferior Parietal Lobule BA40 |
| | | 62 | -22 | 28 | 2E-02 | 5E-05 | 3.9 | R Inferior Parietal Lobule BA40 |
| 10 | 1408 | 10 | -18 | 8 | 4E-02 | 9E-10 | 6.0 | R Thalamus |
| **3. Music imagery** | | | | | | | | |
| *a. MedFG L: 3287 foci, 161 experiments, 2114 subjects (x=0, y=6, z=58)* | | | | | | | | |
| 1 | 59384 | -34 | 24 | 0 | 1E-01 | 1E-26 | 10.6 | L Insula BA13 |
| | | -48 | 4 | 34 | 1E-01 | 3E-24 | 10.1 | L Precentral Gyrus BA6 |
| | | -36 | -4 | 54 | 9E-02 | 1E-19 | 9.0 | L Precentral Gyrus BA6 |
| | | -40 | -4 | 54 | 9E-02 | 2E-19 | 9.0 | L Precentral Gyrus BA6 |
| | | -48 | -4 | 48 | 9E-02 | 2E-19 | 8.9 | L Precentral Gyrus BA4 |
| | | -10 | -18 | 6 | 9E-02 | 4E-17 | 8.3 | L Thalamus |
| | | -50 | 12 | 0 | 9E-02 | 5E-17 | 8.3 | L Insula BA13 |
| | | 10 | -16 | 6 | 7E-02 | 2E-11 | 6.6 | R Thalamus |
| | | 20 | 2 | 2 | 6E-02 | 2E-10 | 6.3 | R Lentiform Nucleus |
| | | -24 | 2 | -2 | 6E-02 | 2E-10 | 6.2 | L Lentiform Nucleus |
| | | -22 | 4 | 2 | 6E-02 | 3E-10 | 6.2 | L Lentiform Nucleus |
| | | -24 | -4 | 6 | 6E-02 | 2E-09 | 5.9 | L Lentiform Nucleus |
| | | -12 | -2 | 6 | 6E-02 | 4E-09 | 5.7 | L Thalamus |
| | | -48 | 28 | 24 | 4E-02 | 9E-06 | 4.3 | L Middle Frontal Gyrus BA46 |
| | | -56 | -20 | 22 | 4E-02 | 2E-05 | 4.1 | L Postcentral Gyrus BA40 |
| | | -58 | -16 | 26 | 4E-02 | 7E-05 | 3.8 | L Postcentral Gyrus BA3 |
| 2 | 26592 | 38 | 22 | -4 | 9E-02 | 2E-19 | 8.9 | R Insula |
| | | 36 | 20 | 0 | 9E-02 | 3E-19 | 8.9 | R Claustrum |
| | | 46 | 2 | 48 | 8E-02 | 2E-16 | 8.2 | R Middle Frontal Gyrus BA6 |

| | | | | | | | | |
|---|---|---|---|---|---|---|---|---|
| | | 36 | -2 | 54 | 8E-02 | 4E-14 | 7.5 | R Precentral Gyrus BA6 |
| | | 54 | 2 | 44 | 7E-02 | 3E-13 | 7.2 | R Precentral Gyrus BA6 |
| | | 48 | 12 | 26 | 6E-02 | 5E-09 | 5.7 | R Inferior Frontal Gyrus BA9 |
| | | 48 | 16 | -2 | 6E-02 | 7E-09 | 5.7 | R Insula BA13 |
| | | 52 | -10 | 40 | 5E-02 | 5E-07 | 4.9 | R Precentral Gyrus BA4 |
| 3 | 21456 | 0 | 6 | 58 | 5E-01 | 0E+00 | 36.8 | L Medial Frontal Gyrus BA6 |
| 4 | 14040 | -28 | -54 | 50 | 8E-02 | 1E-15 | 7.9 | L Superior Parietal Lobule BA7 |
| | | -24 | -66 | 52 | 7E-02 | 8E-14 | 7.4 | L Superior Parietal Lobule BA7 |
| | | -44 | -42 | 46 | 5E-02 | 9E-08 | 5.2 | L Inferior Parietal Lobule BA40 |
| | | -38 | -48 | 58 | 4E-02 | 1E-05 | 4.2 | L Inferior Parietal Lobule BA40 |
| | | -6 | -72 | 56 | 4E-02 | 1E-04 | 3.7 | L Precuneus BA7 |
| | | -46 | -32 | 58 | 4E-02 | 1E-04 | 3.6 | L Inferior Parietal Lobule BA40 |
| | | -40 | -24 | 58 | 3E-02 | 7E-04 | 3.2 | L Postcentral Gyrus BA3 |
| 5 | 7480 | 44 | -40 | 48 | 5E-02 | 3E-08 | 5.4 | R Inferior Parietal Lobule BA40 |
| | | 32 | -60 | 54 | 5E-02 | 7E-07 | 4.8 | R Superior Parietal Lobule BA7 |
| | | 18 | -68 | 54 | 4E-02 | 2E-06 | 4.6 | R Precuneus BA7 |
| | | 38 | -50 | 50 | 4E-02 | 2E-06 | 4.6 | R Inferior Parietal Lobule BA40 |
| | | 26 | -64 | 56 | 4E-02 | 2E-06 | 4.6 | R Precuneus BA7 |
| 6 | 5064 | -42 | -62 | -16 | 6E-02 | 1E-10 | 6.3 | L Declive |
| | | -32 | -58 | -30 | 5E-02 | 2E-07 | 5.1 | L Cerebellum |
| | | -40 | -68 | -28 | 4E-02 | 4E-06 | 4.5 | L Tuber |
| 7 | 3808 | 32 | -62 | -26 | 6E-02 | 8E-10 | 6.0 | R Culmen |
| *b. SPL L: 1325 foci, 64 experiments, 875 subjects (x=-34, y=-58, z=56)* | | | | | | | | |
| 1 | 14816 | -34 | -58 | 56 | 3E-01 | 0E+00 | 23.4 | L Superior Parietal Lobule BA7 |
| | | -42 | -38 | 36 | 4E-02 | 7E-09 | 5.7 | L Supramarginal Gyrus BA40 |
| | | -20 | -72 | 52 | 4E-02 | 7E-08 | 5.3 | L Precuneus BA7 |
| | | -50 | -34 | 40 | 3E-02 | 3E-07 | 5.0 | L Inferior Parietal Lobule BA40 |
| | | -42 | -28 | 52 | 3E-02 | 7E-06 | 4.3 | L Inferior Parietal Lobule BA40 |
| | | -42 | -30 | 48 | 3E-02 | 9E-06 | 4.3 | L Inferior Parietal Lobule BA40 |
| | | -46 | -40 | 48 | 2E-02 | 2E-04 | 3.6 | L Inferior Parietal Lobule BA40 |
| 2 | 8528 | -2 | 8 | 48 | 5E-02 | 3E-11 | 6.5 | L Cingulate Gyrus BA24 |
| | | 4 | 4 | 60 | 5E-02 | 4E-11 | 6.5 | R Medial Frontal Gyrus BA6 |
| | | -2 | 4 | 60 | 5E-02 | 1E-10 | 6.3 | L Medial Frontal Gyrus BA6 |
| | | 4 | 16 | 48 | 4E-02 | 2E-08 | 5.5 | R Superior Frontal Gyrus BA6 |
| | | 0 | 24 | 44 | 3E-02 | 3E-06 | 4.6 | L Medial Frontal Gyrus BA6 |
| 3 | 7384 | 34 | -54 | 48 | 5E-02 | 1E-11 | 6.7 | R Superior Parietal Lobule BA7 |
| | | 34 | -58 | 56 | 5E-02 | 6E-11 | 6.4 | R Superior Parietal Lobule BA7 |
| | | 16 | -64 | 52 | 3E-02 | 8E-06 | 4.3 | R Precuneus BA7 |
| | | 4 | -64 | 48 | 2E-02 | 1E-04 | 3.7 | R Precuneus BA7 |
| 4 | 6584 | 50 | 8 | 26 | 5E-02 | 9E-12 | 6.7 | R Inferior Frontal Gyrus BA9 |
| | | 44 | -4 | 48 | 3E-02 | 1E-06 | 4.7 | R Precentral Gyrus BA6 |
| | | 36 | -2 | 54 | 3E-02 | 2E-05 | 4.1 | R Precentral Gyrus BA6 |
| 5 | 5968 | -46 | 8 | 26 | 5E-02 | 1E-12 | 7.0 | L Inferior Frontal Gyrus BA9 |
| | | -50 | -8 | 42 | 3E-02 | 1E-05 | 4.2 | L Precentral Gyrus BA4 |
| | | -54 | 2 | 40 | 3E-02 | 5E-05 | 3.9 | L Precentral Gyrus BA6 |
| | | -44 | 24 | 26 | 2E-02 | 8E-05 | 3.8 | L Middle Frontal Gyrus BA9 |
| 6 | 3184 | -30 | 24 | 4 | 5E-02 | 2E-10 | 6.3 | L Insula BA13 |
| | | -22 | -6 | 6 | 2E-02 | 1E-04 | 3.7 | L Lentiform Nucleus |
| | | -20 | 2 | 0 | 2E-02 | 3E-04 | 3.4 | L Lentiform Nucleus |
| 7 | 2056 | 36 | 22 | -10 | 3E-02 | 2E-07 | 5.1 | R Inferior Frontal Gyrus BA47 |
| | | 36 | 22 | -2 | 3E-02 | 7E-07 | 4.8 | R Insula |
| *c. THA L: 2413 foci, 113 experiments, 1571 subjects (x=-14, y=-14, z=8)* | | | | | | | | |
| 1 | 64248 | -14 | -14 | 8 | 4E-01 | 0E+00 | 30.8 | L Thalamus |
| | | 12 | -14 | 8 | 2E-01 | 0E+00 | 15.2 | R Thalamus |
| | | -32 | 20 | 4 | 1E-01 | 6E-22 | 9.6 | L Insula BA13 |
| | | 34 | 22 | 2 | 8E-02 | 4E-16 | 8.1 | R Claustrum |
| | | 26 | 6 | 4 | 7E-02 | 3E-14 | 7.5 | R Lentiform Nucleus |
| | | 50 | 14 | -2 | 6E-02 | 3E-11 | 6.5 | R Insula BA13 |
| | | -8 | -20 | -10 | 5E-02 | 2E-09 | 5.9 | L Brainstem |
| | | -54 | 12 | 2 | 5E-02 | 2E-08 | 5.5 | L Precentral Gyrus BA44 |
| | | -40 | 2 | 6 | 5E-02 | 2E-08 | 5.5 | L Insula BA13 |
| | | 4 | -24 | -8 | 5E-02 | 5E-08 | 5.3 | R Brainstem |
| | | 52 | 12 | 22 | 5E-02 | 7E-08 | 5.3 | R Inferior Frontal Gyrus BA9 |
| | | 54 | 14 | 8 | 4E-02 | 2E-07 | 5.1 | R Precentral Gyrus BA44 |
| | | 54 | 10 | 32 | 4E-02 | 4E-07 | 4.9 | R Inferior Frontal Gyrus BA9 |
| | | -12 | 8 | 0 | 4E-02 | 3E-06 | 4.5 | L Caudate |

| | | | | | | | | |
|---|---|---|---|---|---|---|---|---|
| | | 42 | 2 | 34 | 3E-02 | 2E-04 | 3.6 | R Precentral Gyrus BA6 |
| 2 | 20032 | 0 | 8 | 48 | 9E-02 | 1E-20 | 9.3 | L Cingulate Gyrus BA24 |
| | | -2 | 0 | 58 | 8E-02 | 2E-16 | 8.1 | L Medial Frontal Gyrus BA6 |
| | | 8 | 28 | 28 | 4E-02 | 5E-07 | 4.9 | R Cingulate Gyrus BA32 |
| | | 20 | 2 | 56 | 3E-02 | 8E-04 | 3.2 | R Sub-Gyral BA6 |
| 3 | 13944 | -38 | -20 | 58 | 6E-02 | 4E-11 | 6.5 | L Precentral Gyrus BA4 |
| | | -30 | -6 | 50 | 5E-02 | 9E-09 | 5.6 | L Middle Frontal Gyrus BA6 |
| | | -44 | -38 | 48 | 4E-02 | 8E-07 | 4.8 | L Inferior Parietal Lobule BA40 |
| | | -50 | -26 | 18 | 4E-02 | 2E-06 | 4.7 | L Superior Temporal Gyrus BA41 |
| | | -56 | -44 | 22 | 4E-02 | 1E-05 | 4.3 | L Superior Temporal Gyrus BA13 |
| | | -38 | -46 | 42 | 4E-02 | 2E-05 | 4.1 | L Inferior Parietal Lobule BA40 |
| | | -56 | -18 | 16 | 4E-02 | 2E-05 | 4.1 | L Postcentral Gyrus BA43 |
| | | -50 | -30 | 34 | 3E-02 | 3E-05 | 4.0 | L Inferior Parietal Lobule BA40 |
| | | -60 | -16 | 22 | 3E-02 | 4E-05 | 3.9 | L Postcentral Gyrus BA43 |
| 4 | 3536 | -52 | 10 | 32 | 5E-02 | 6E-08 | 5.3 | L Inferior Frontal Gyrus BA9 |
| | | -46 | 24 | 26 | 3E-02 | 1E-04 | 3.7 | L Middle Frontal Gyrus BA9 |
| 5 | 3136 | 56 | -30 | 22 | 4E-02 | 2E-07 | 5.1 | R Insula BA13 |
| | | 64 | -22 | 18 | 4E-02 | 5E-06 | 4.4 | R Postcentral Gyrus BA40 |
| 6 | 2776 | 30 | -58 | -30 | 5E-02 | 2E-08 | 5.5 | R Cerebellum |
| | | 22 | -50 | -22 | 4E-02 | 1E-05 | 4.2 | R Culmen |
| | | 40 | -52 | -26 | 3E-02 | 1E-04 | 3.6 | R Culmen |
| 7 | 2192 | 2 | -62 | -18 | 4E-02 | 2E-06 | 4.7 | R Culmen |
| | | 10 | -70 | -18 | 4E-02 | 6E-06 | 4.4 | R Declive |
| | | 2 | -46 | -10 | 3E-02 | 7E-04 | 3.2 | L Cerebellar Lingual |
| *d. PreCG BA6 R: 1011 foci, 37 experiments, 569 subjects (x=54, y=-0, z=50)* | | | | | | | | |
| 1 | 18304 | -48 | -4 | 52 | 6E-02 | 4E-15 | 7.8 | L Precentral Gyrus BA4 |
| | | -34 | 20 | 4 | 5E-02 | 4E-12 | 6.9 | L Insula BA13 |
| | | -50 | 8 | 2 | 4E-02 | 3E-09 | 5.8 | L Insula BA13 |
| | | -54 | 2 | 38 | 4E-02 | 6E-09 | 5.7 | L Precentral Gyrus BA6 |
| | | -56 | 6 | 6 | 4E-02 | 1E-08 | 5.6 | L Superior Temporal Gyrus BA22 |
| | | -58 | 0 | 16 | 3E-02 | 5E-07 | 4.9 | L Precentral Gyrus BA4 |
| | | -54 | 8 | 14 | 3E-02 | 6E-07 | 4.9 | L Inferior Frontal Gyrus BA44 |
| | | -42 | 6 | 24 | 3E-02 | 8E-07 | 4.8 | L Precentral Gyrus BA6 |
| 2 | 11216 | -4 | 0 | 62 | 7E-02 | 4E-22 | 9.6 | L Medial Frontal Gyrus BA6 |
| | | -4 | 18 | 36 | 4E-02 | 2E-08 | 5.5 | L Cingulate Gyrus BA32 |
| | | 8 | 12 | 38 | 3E-02 | 5E-08 | 5.3 | R Cingulate Gyrus BA32 |
| 3 | 5928 | 54 | 0 | 48 | 2E-01 | 0E+00 | 19.6 | R Precentral Gyrus BA4 |
| | | 58 | 4 | 34 | 2E-02 | 3E-05 | 4.0 | R Precentral Gyrus BA6 |
| | | 34 | -6 | 58 | 2E-02 | 4E-04 | 3.4 | R Precentral Gyrus BA6 |
| 4 | 5624 | 50 | 10 | 2 | 4E-02 | 5E-10 | 6.1 | R Insula BA13 |
| | | 38 | 22 | 0 | 3E-02 | 9E-07 | 4.8 | R Insula BA13 |
| | | 56 | 14 | -12 | 3E-02 | 3E-06 | 4.5 | R Superior Temporal Gyrus BA22 |
| 5 | 4824 | 24 | 6 | 4 | 5E-02 | 5E-13 | 7.1 | R Putamen |
| | | 14 | -10 | 6 | 3E-02 | 1E-07 | 5.2 | R Thalamus |
| | | 22 | 0 | -8 | 2E-02 | 8E-05 | 3.8 | R Globus Pallidus |
| 6 | 3480 | 62 | -32 | 10 | 3E-02 | 7E-08 | 5.3 | R Superior Temporal Gyrus BA42 |
| | | 56 | -32 | 4 | 3E-02 | 9E-08 | 5.2 | R Superior Temporal Gyrus BA22 |
| 7 | 3424 | -52 | -18 | 2 | 3E-02 | 2E-07 | 5.1 | L Superior Temporal Gyrus BA22 |
| | | -62 | -30 | 8 | 3E-02 | 1E-06 | 4.7 | L Superior Temporal Gyrus BA42 |
| 8 | 2576 | -22 | 4 | 6 | 5E-02 | 2E-13 | 7.3 | L Putamen |
| 9 | 1496 | -52 | -40 | 22 | 4E-02 | 2E-09 | 5.9 | L Superior Temporal Gyrus BA13 |

BA, Brodmann area; ROIs, regions-of-interest; ALE, anatomic likelihood estimation; P, p-value; Z, peak z-value; R, right; L, left. **ROIs**: CRBL, cerebellum; INS, insula; MedFG, medial frontal gyrus; PreCG, precentral gyrus (primary motor cortex or M1); PUT, putamen; SPL, superior parietal lobule; STG, superior temporal gyrus (primary auditory cortex); THA, thalamus. Music-related ROIs were created in Mango (http://rii.uthscsa.edu/mango//userguide.html) with a 5mm-radius sphere.
Last search in Sleuth, 01.09.2021 (http://www.brainmap.org/sleuth/).

**Supplementary Table 5. Functional characterization of brain regions resulted from primary outcomes (n = 17 ROIs) according to BrainMap database .**

| | |
|---|---|
| **1. Music perception** | |
| *a. STG BA22 R: 824 foci, 43 experiments, 610 subjects (x=52, y=-20, z=4)* | |
| Action | Execution, speech, motor learning, observation |
| Cognition | Attention, phonology, semantics, speech, explicit memory, working memory, **music**, reasoning, spatial |
| Emotion | Anxiety, sadness, positive emotion, reward/gain, valence |
| Interoception | Sexuality |
| Perception | Audition, somesthesis, vision, shape |
| Paradigms | Acupuncture, classical conditioning, cued explicit recognition/recall, drawing, emotion induction, encoding, face discrimination, finger tapping/button pressing, flexion/extension, mental rotation, **music comprehension, music production**, n-back, naming (overt), oddball discrimination, paired associate recall, passive listening, passive viewing, phonological discrimination, pitch discrimination, reading overt, reading covert, reasoning/problem solving, recitation/repetition, reward, semantic discrimination, sequence recall/learning, sexual arousal/gratification, tone discrimination, visuospatial attention |
| *b. STG BA22 L: 1143 foci, 60 experiments, 850 subjects (x=-54, y=-16, z=2)* | |
| Action | Execution, speech, inhibition, observation |
| Cognition | Attention, phonology, semantics, speech, explicit memory, working memory, **music**, reasoning, somatic, spatial |
| Emotion | Anger, fear, guilt, sadness, positive emotion, happiness, reward/gain, valence |
| Interoception | Sexuality |
| Perception | Audition, vision, shape |
| Paradigms | Counting/calculation, cued explicit recognition/recall, emotion induction, encoding, face discrimination, film viewing, finger tapping/button pressing, flexion/extension, go/no-go, hand-eye coordination, imagined objects/scene, mental rotation, **music comprehension, music production**, oddball discrimination, paired associate recall, passive listening, passive viewing, phonological discrimination, pitch discrimination, reading overt, reading covert, reasoning/problem solving, recitation/repetition, reward, semantic discrimination, sequence recall/learning, sexual arousal/gratification, tone discrimination, visuospatial attention, word generation (covert), word generation (overt) |
| *c. MedFG BA6 L: 1773 foci, 58 experiments, 838 subjects (x=-2, y=-2, z=66)* | |
| Action | Execution, speech, imagination, inhibition, motor learning, observation |
| Cognition | Attention, language, phonology, semantics, speech, explicit memory, working memory, **music**, reasoning, social cognition, somatic, temporal |
| Emotion | Sadness, positive emotion |
| Interoception | - |
| Perception | Audition, gustation, somesthesis, pain, vision, color, motion |
| Paradigms | Acupuncture, affective words, counting/calculation, cued explicit recognition/recall, deception, delayed match to sample, emotion induction, episodic recall, estimation, face discrimination, finger tapping/button pressing, flanker, flexion/extension, go/no-go, imagined movement, imagined objects/scene, tongue movement, **music comprehension, music production**, n-back, pain discrimination, passive listening, passive viewing, pitch discrimination, manual tracking, reading overt, reading covert, recitation/repetition, saccades, semantic discrimination, sequence recall/learning, tactile discrimination, tone discrimination, visual object identification, visual tracking, visuospatial attention, writing |
| *d. PUT R: 1821 foci, 70 experiments, 1052 subjects (x=22, y=8, z=6)* | |
| Action | Execution, speech, imagination, inhibition, observation |
| Cognition | Attention, orthography, phonology, semantics, speech, explicit memory, working memory, **music**, reasoning, somatic, spatial, temporal |
| Emotion | Anger, fear, reward/gain, valence |
| Interoception | Baroregulation, respiration regulation, sexuality |
| Perception | Audition, gustation, olfaction, somesthesis, pain, vision, color, motion, shape |
| Paradigms | Affective pictures, affective words, anti-saccades, chewing/swallowing, counting/calculation, cued explicit recall, encoding, face discrimination, film viewing, finger tapping/button press, flexion/extension, gambling, go/no-go, hypercapnia, imagined movement, isometric force, magnitude comparison, **music comprehension, music production**, olfactory discrimination, pain discrimination, paired associate recall, passive listening, passive viewing, pitch discrimination, pointing, reading covert, reading overt, reasoning, recitation, reward, saccades, semantic discrimination, sequence recall/learning, sexual arousal, Stroop-color, tactile discrimination, task switching, tone discrimination, visual object identification, visuospatial attention, word generation |
| *e. PUT L: 1493 foci, 69 experiments, 1139 subjects (x=-22, y=4, z=6)* | |
| Action | Execution, speech, inhibition, preparation |
| Cognition | Attention, phonology, semantics, speech, explicit memory, working memory, **music**, reasoning, spatial, temporal |
| Emotion | Negative emotion, sadness, positive emotion, happiness, reward/gain, valence |
| Interoception | Gastrointestinal/Genitourinary, hunger, sexuality |
| Perception | Audition, gustation, olfaction, somesthesis, pain, vision, color, motion, shape |
| Paradigms | Acupuncture, affective pictures, chewing/swallowing, classical conditioning, cued explicit recognition/recall, deception, delayed match to sample, emotion induction, encoding, episodic recall, estimation, face discrimination, film viewing, finger |

| | |
|---|---|
| | tapping/button pressing, flanker, flexion/extension, go/no-go, hunger, imagined movement, isometric force, micturition, **music comprehension**, n-back, naming covert, olfactory discrimination, orthographic discrimination, pain discrimination, passive listening, passive viewing, phonological discrimination, pitch discrimination, reading overt, recitation/repetition, reward, semantic discrimination, sexual arousal/gratification, Stroop-color, task switching, taste, tone discrimination, visual object identification, visuospatial attention, Wisconsin card sorting, word generation (covert) |
| *f. CRBL L: 1201 foci, 47 experiments, 688 subjects (x=-28, y=-64, z=-26)* | |
| Action | Execution, speech, imagination, inhibition, preparation |
| Cognition | Attention, language orthography, phonology, semantics, speech, explicit memory, working memory, **music**, reasoning |
| Emotion | Negative emotion, anger, sadness, positive emotion, happiness, valence |
| Interoception | Gastrointestinal/Genitourinary, sexuality, sleep, vestibular |
| Perception | Audition, somesthesis, pain, vision, shape |
| Paradigms | Affective words, counting/calculation, cued explicit recognition/recall, deception, delayed match to sample, divided auditory attention, emotion induction, encoding, episodic recall, face discrimination, film viewing, finger tapping/button pressing, go/no-go, imagined movement, micturition, **music comprehension, music production**, pain discrimination, paired associate recall, passive listening, passive viewing, pitch discrimination, reading overt, reading covert, recitation/repetition, reward, semantic discrimination, sequence recall/learning, sexual arousal/gratification, tactile discrimination, tone discrimination, vestibular stimulation, visual object identification, visuospatial attention, word generation (covert) |
| *g. INS L: 1787 foci, 87 experiments, 1332 subjects (x=-32, y=18, z=10)* | |
| Action | Execution, speech, imagination, inhibition, observation, preparation |
| Cognition | Attention, orthography, phonology, semantics, speech, syntax, explicit memory, implicit memory, working memory, **music**, reasoning, social cognition, somatic, spatial |
| Emotion | Negative emotion, anxiety, embarrassment, fear, positive emotion, happiness, reward/gain, valence |
| Interoception | Sexuality |
| Perception | Audition, gustation, olfaction, pain, vision, color, motion, shape |
| Paradigms | Affective words, chewing/swallowing, classical conditioning, counting/calculation, cued explicit recognition/recall, deception, delayed match to sample, driving, emotion induction, encoding, face discrimination, figurative language, film viewing, finger tapping/button pressing, flanker, gambling, go/no-go, grasping, imagined movement, imagined objects/scene, isometric force, multi-tasking, **music comprehension**, oddball discrimination, olfactory discrimination, orthographic discrimination, pain discrimination, passive listening, passive viewing, phonological discrimination, reading overt, reading covert, reasoning/problem solving, reward, saccades, semantic discrimination, sexual arousal/gratification, Stroop-color, task-switching, taste, theory of mind, tone discrimination, visual object identification, visuospatial attention, Wisconsin card sorting test, word generation (overt) |
| *h. PreCG R: 1613 foci, 63 experiments, 931 subjects (x=54, y=0, z=46)* | |
| Action | Execution, speech, imagination, inhibition, motor learning, observation, preparation |
| Cognition | Attention, orthography, phonology, semantics, speech, explicit memory, implicit memory, working memory, **music**, reasoning, social cognition, temporal |
| Emotion | Intensity, negative emotion, embarrassment, fear, sadness, positive emotion, happiness, reward/gain |
| Interoception | - |
| Perception | Audition, somesthesis, pain, vision, motion |
| Paradigms | Affective pictures, affective words, chewing/swallowing, counting/calculation, cued explicit recognition/recall, deception, delay discounting, delayed match to sample, driving, emotion induction, encoding, episodic recall, face discrimination, film viewing, finger tapping/button pressing, flexion/extension, go/no-go, grasping, imagined movement, multitasking, **music comprehension, music production**, pain discrimination, passive listening, passive viewing, pitch discrimination, pointing, reading overt, reading covert, reasoning/problem solving, recitation/repetition, reward, saccades, semantic discrimination, Stroop-color, theory of mind, tone discrimination, visual pursuit, visuospatial attention, word stem generation |
| **2. Music production** | |
| *a. STG BA42 R: 753 foci, 32 experiments, 455 subjects (x=66, y=-24, z=10)* | |
| Action | Execution, speech, observation |
| Cognition | Attention, language, phonology, semantics, speech, syntax, explicit memory, working memory, **music**, reasoning, social cognition |
| Emotion | Negative emotion, anxiety, disgust, sadness, reward/gain |
| Interoception | Thermoregulation |
| Perception | Audition, pain, vision, motion, shape |
| Paradigms | Affective pictures, counting/calculation, cued explicit recognition/recall, emotion induction, face discrimination, film viewing, finger tapping/button pressing, go/no-go, **music comprehension, music production**, naming, oddball discrimination, pain discrimination, passive listening, passive viewing, phonological discrimination, pitch discrimination, reading overt, reasoning/problem solving, recitation/repetition, reward, semantic discrimination, sequence recall, theory of mind, tone discrimination, visual object identification, visual pursuit/tracking, word generation (overt) |
| *b. PreCG L: 1372 foci, 79 experiments, 1034 subjects (x=-48, y=0, z=42)* | |
| Action | Execution, speech, imagination, inhibition, observation |
| Cognition | Attention, language, orthography, phonology, semantics, speech, syntax, explicit memory, working memory, **music**, reasoning, temporal |

| | |
|---|---|
| Emotion | Fear, sadness |
| Interoception | Respiration regulation, sexuality |
| Perception | Audition, somesthesis, vision, motion, shape |
| Paradigms | Counting/calculation, cued explicit recognition/recall, delayed match to sample, encoding, face discrimination, film viewing, finger tapping/button pressing, go/no-go, hypercapnia/air hunger, imagined movement, mental rotation, **music comprehension, music production**, n-back, naming covert, naming overt, orthographic discrimination, paired associate recall, passive viewing, phonological discrimination, pitch discrimination, pointing, pursuit rotor/manual tracking, reading overt, reading covert, reasoning/problem solving, recitation/repetition, saccades, semantic discrimination, sequence recall/learning, sexual arousal/gratification, tactile discrimination, task-switching, visual pursuit/tracking, visuospatial attention, word generation (covert), word generation (overt) |
| *c. MedFG BA6 L: 403 foci, 21 experiments, 261 subjects (x=-10, y=-10, z=52)* | |
| Action | Execution, imagination, inhibition |
| Cognition | Attention, orthography, semantics, explicit memory, working memory, **music**, social cognition |
| Emotion | Positive emotion, reward/gain |
| Interoception | Respiration regulation |
| Perception | Somesthesis, vision, color |
| Paradigms | Affective pictures, affective words, competition/cooperation, deception, finger tapping/button pressing, flexion/extension, go/no-go, hypercapnia/air hunger, imagined movement, isometric force, **music comprehension, music production**, n-back, reading covert, reward, semantic discrimination, sequence recall/learning, tactile discrimination, trauma recall, videogames, visual pursuit/tracking, visuospatial attention |
| *d. STG BA41 L: 529 foci, 38 experiments, 522 subjects (x=-42, y=-28, z=6)* | |
| Action | Execution, speech, imagination, observation, preparation |
| Cognition | Attention, language, phonology, semantics, speech, explicit memory, working memory, **music**, social cognition, spatial |
| Emotion | Negative emotion, anger, anxiety, fear, sadness, positive emotion, happiness |
| Interoception | - |
| Perception | Audition, somesthesis, vision |
| Paradigms | Affective words, deception, emotion induction, episodic recall, face discrimination, film viewing, finger tapping/button pressing, fixation, flexion/extension, hand-eye coordination, imagined movement, imagined objects/scenes, **music comprehension, music production**, naming overt, oddball discrimination, pain discrimination, passive listening, passive viewing, phonological discrimination, pitch discrimination, recitation/repetition, semantic discrimination, tone discrimination, visual object identification, visuospatial attention, word generation covert |
| *e. PreCG R: 1105 foci, 51 experiments, 846 subjects (x=54, y=4, z=32)* | |
| Action | Execution, speech, imagination, inhibition, observation |
| Cognition | Attention, orthography, phonology, speech, syntax, explicit memory, working memory, **music**, reasoning, social cognition, temporal |
| Emotion | Negative emotion, anger, fear, sadness, positive emotion, happiness, valence |
| Interoception | Respiration regulation, sexuality |
| Perception | Audition, gustation, pain, vision, motion |
| Paradigms | Affective pictures, affective words, chewing/swallowing, counting/calculation, cued explicit recognition/recall, delayed match to sample, emotion induction, encoding, face discrimination, film viewing, finger tapping/button pressing, fixation, flexion/extension, go/no-go, grasping, hypercapnia/air hunger, imagined movement, imagined objects/scenes, **music comprehension, music production**, orthographic discrimination, pain discrimination, paired passive viewing, phonological discrimination, pitch discrimination, pointing, pursuit rotor/manual tracking, reading overt, reading covert, reasoning/problem solving, recitation/repetition, saccades, sexual arousal, theory of mind, visual object identification, visual pursuit/tracking, visuospatial attention, word generation (overt), word completion covert, word completion overt, writing |
| **3. Music imagery** | |
| *a. MedFG L: 3287 foci, 161 experiments, 2114 subjects (x=0, y=6, z=58)* | |
| Action | Execution, speech, imagination, inhibition, observation, preparation |
| Cognition | Attention, language, phonology, semantics, speech, syntax, explicit memory, working memory, **music**, reasoning, social cognition, somatic, spatial, temporal |
| Emotion | Negative emotion, anger, anxiety, sadness, happiness, humor, reward/gain, valence |
| Interoception | Heartbeat detection, sexuality |
| Perception | Audition, somesthesis, pain, vision, motion, shape |
| Paradigms | Affective pictures, anti-saccades, chewing/swallowing, classical conditioning, competition/cooperation, counting/calculation, cued explicit recognition/recall, delayed match to sample, drawing, emotion induction, encoding, face discrimination, film viewing, finger tapping/button pressing, fixation, flexion/extension, go/no-go, grasping, imagined movement, imagined objects/scenes, isometric force, lexical decision, mental rotation, **music comprehension, music production,** n-back,, naming covert, naming overt, orthographic discrimination, pain discrimination, paired associate recall, passive listening, phonological discrimination, pitch discrimination, pointing, reading overt, reading covert, reasoning/problem solving, recitation/repetition, reward, saccades, semantic discrimination, sequence recall/learning, sexual arousal/gratification, Stroop-color, Stroop-emotional, syntactic discrimination, tactile discrimination, task-switching, tone discrimination, Tower of London, videogames, visual object identification, visuospatial attention, Wisconsin card sorting test, word generation (covert), word generation (overt), word completion |

| | |
|---|---|
| *b. SPL L: 1325 foci, 64 experiments, 875 subjects (x=-34, y=-58, z=56)* | |
| Action | Execution, speech, imagination, inhibition, observation |
| Cognition | Attention, orthography, phonology, semantics, speech, explicit memory, working memory, **music**, reasoning, social cognition, somatic, spatial |
| Emotion | Negative emotion, anxiety, disgust, positive emotion, happiness, reward/gain |
| Interoception | Sexuality |
| Perception | Audition, gustation, somesthesis, pain, vision, color, motion, shape |
| Paradigms | Affective pictures, anti-saccades, counting/calculation, cued explicit recognition/recall, delay discounting, delayed match to sample, emotion induction, encoding, face discrimination, film viewing, finger tapping/button pressing, flexion/extension, gambling, go/no-go, imagined movement, isometric force, magnitude comparison, mental rotation, **music comprehension**, n-back, naming, orthographic discrimination, pain discrimination, paired associate recall, passive viewing, pitch discrimination, reading overt, reading covert, reasoning/problem solving, reward, saccades, semantic discrimination, sequence recall/learning, sexual arousal/gratification, Stroop-color, tactile discrimination, task-switching, visual object identification, visual pursuit/tracing, visuospatial attention, word generation (overt) |
| *c. THA L: 2413 foci, 113 experiments, 1571 subjects (x=-14, y=-14, z=8)* | |
| Action | Execution, speech, imagination, inhibition, motor learning |
| Cognition | Attention, language, orthography, phonology, semantics, speech, explicit memory, working memory, **music**, reasoning, social cognition, temporal |
| Emotion | Negative emotion, anxiety, fear, sadness, positive emotion, happiness, reward/gain, valence |
| Interoception | Gastrointestinal/Genitourinary, respiration regulation, sleep, thermoregulation |
| Perception | Audition, gustation, somesthesis, pain, vision, color, motion, shape |
| Paradigms | Affective words, anti-saccades, classical conditioning, counting/calculation, cued explicit recognition/recall, delay discounting, delayed match to sample, emotion induction, encoding, episodic recall, face discrimination, film viewing, finger tapping/button pressing, flanker, flexion/extension, gambling, go/no-go, grasping, hypercapnia, air hunger, imagined objects/scene, isometric force, lexical decision, micturition, motor learning, **music comprehension, music production**, n-back, naming, oddball discrimination, orthographic discrimination, pain discrimination, paired associate recall, passive listening, passive viewing, phonological discrimination, pointing, pursuit rotor/manual tracking, reading overt, reading covert, reasoning/problem solving, recitation/repetition, reward, saccades, sequence recall/learning, Stroop-counting, tactile discrimination, task-switching, taste, tone discrimination, transcranial magnetic stimulation, trauma recall, videogames, visual object identification, visual pursuit/tracking, visuospatial attention, word generation (covert), word generation (overt), writing |
| *d. PreCG BA6: 1011 foci, 37 experiments, 569 subjects (x=54, y=-0, z=50)* | |
| Action | Execution, speech, imagination, motor learning, observation |
| Cognition | Attention, orthography, semantics, speech, explicit memory, working memory, **music**, reasoning |
| Emotion | Intensity, negative emotion, sadness, positive emotion, happiness, reward/gain |
| Interoception | Thermoregulation |
| Perception | Audition, olfaction, pain, vision, motion |
| Paradigms | Affective pictures, affective words, chewing/swallowing, counting/calculation, cued explicit recognition/recall, delay discounting, delayed match to sample, emotion induction, face discrimination, film viewing, finger tapping/button pressing, flexion/extension, go/no-go, imagined movement, multi-tasking, **music comprehension, music production**, olfactory discrimination, pain discrimination, passive listening, passive viewing, phonological discrimination, pitch discrimination, reading overt, reading covert, reasoning/problem solving, recitation/repetition, reward, saccades, semantic discrimination, tactile discrimination, task-switching, tone discrimination, visuospatial attention |

BA, Brodmann area; ROIs, regions-of-interest; ALE, anatomic likelihood estimation; P, p-value; Z, peak z-value; R, right; L, left. **ROIs**: CRBL, cerebellum; INS, insula; MedFG, medial frontal gyrus; PreCG, precentral gyrus (primary motor cortex or M1); PUT, putamen; SPL, superior parietal lobule; STG, superior temporal gyrus (primary auditory cortex); THA, thalamus. Music-related ROIs were created in Mango (http://rii.uthscsa.edu/mango//userguide.html) with a 5mm-radius sphere.
Last search in Sleuth, 01.09.2021 (http://www.brainmap.org/sleuth/).

**Supplementary Table 6. FSN robustness assessment for significant ALE maps of music perception, production, and imagery**

| Cluster number | Volume ($mm^3$) | MNI coordinates x | y | z | ALE | Label (Side, region) | Contributing studies (k) | FSN |
|---|---|---|---|---|---|---|---|---|
| 1. Music perception: 1898 foci, 105 experiments, 2035 subjects, minimum FSN = 32 | | | | | | | | |
| 1 | 24296 | 52 | -20 | 4 | 1E-01 | R Superior Temporal Gyrus BA13 | 75 | 292 |
| 2 | 23864 | -54 | -16 | 2 | 1E-01 | L Superior Temporal Gyrus BA22 | 79 | 292 |
| 3 | 6136 | -2 | -2 | 66 | 5E-02 | L Medial Frontal Gyrus BA6 | 31 | 105 |
| 4 | 3328 | 22 | 8 | 6 | 6E-02 | R Putamen | 21 | 105 |
| 5 | 2304 | -22 | 4 | 6 | 5E-02 | L Putamen | 16 | 50 |
| 6 | 1920 | -28 | -64 | -26 | 5E-02 | L Cerebellum | 13 | 50 |
| 7 | 1816 | -32 | 18 | 10 | 4E-02 | L Insula BA13 | 11 | 75 |
| 8 | 1728 | 54 | 0 | 46 | 6E-02 | R Precentral Gyrus BA4 | 14 | 35 |
| 2. Music production: 499 foci, 19 experiments, 292 subjects, minimum FSN = 6 | | | | | | | | |
| 1 | 3512 | 66 | -24 | 10 | 2E-02 | R Superior Temporal Gyrus BA42 | 8 | 75 |
| 2 | 2920 | -48 | 0 | 42 | 2E-02 | L Precentral Gyrus BA6 | 7 | 57 |
| 3 | 2080 | -10 | -10 | 52 | 2E-02 | L Medial Frontal Gyrus BA6 | 9 | 61 |
| 4 | 1912 | -42 | -28 | 6 | 3E-02 | L Superior Temporal Gyrus BA13 | 5 | 40 |
| 5 | 1072 | 54 | 4 | 32 | 2E-02 | R Precentral Gyrus BA6 | 5 | <6 |
| 3. Music imagery: 263 foci, 15 experiments, 189 subjects, minimum FSN = 5 | | | | | | | | |
| 1 | 3256 | 0 | 6 | 58 | 2E-02 | L Medial Frontal Gyrus BA6 | 10 | 56 |
| 2 | 3128 | -34 | -58 | 56 | 2E-02 | L Superior Parietal Lobule BA7 | 9 | 30 |
| 3 | 1384 | -14 | -14 | 8 | 2E-02 | L Thalamus | 3 | 35 |
| 4 | 944 | 54 | 0 | 50 | 2E-02 | L Precentral Gyrus BA6 | 6 | 10 |

FSN, Fail-Safe N analysis; ALE, anatomic likelihood estimation; BA, Brodmann area; R, right; L, left.

[Citations of included studies]

**Abbreviations**

| | |
|---|---|
| ACC | anterior cingulate cortex |
| AF | arcuate fasciculus |
| AnG | angular gyrus |
| CalC | calcarine cortex |
| CC | corpus callosum |
| CAU | caudate |
| CLAU | claustrum |
| CRBL | cerebellum |
| CST | corticospinal tract |
| CUN | cuneus |
| DLPFC | dorsolateral prefrontal cortex |
| EC | entorhinal cortex |
| Fmaj | forceps major |
| Fmin | forceps minor |
| FO | frontal operculum |
| FusG | fusiform gyrus |
| GP | globus pallidus |
| HG | Helsch's gyrus |
| HIPP | hippocampus |
| IC | internal capsule |
| IF | inferior colliculus |
| IFG | inferior frontal gyrus |
| IFOF | inferior fronto-occipital fasciculus |
| ILF | inferior longitudinal fasciculus |
| INS | insula |
| IPL | inferior parieal lobule |
| ITG | inferior temporal gyrus |
| LG | lingual gyrus |
| LOC | lateral occipital cortex |
| MB | midbrain |
| MCC | middle cingulate cortex |
| MCP | middle cerebellar peduncle |
| MedFG | medial frontal gyrus |
| MidFG | middle frontal gyrus |
| MidTG | middle temporal gyrus |
| OFC | orbitofrontal cortex |
| PaHIPP | parahippocampal gyrus |
| PCC | posterior cingulate cortex |
| PMC | premotor cortex |
| PO | parietal operculum |
| PostCG | postcentral gyrus (primary somatosensory cortex or SI) |
| PP | planum polare |
| PreCG | precentral gyrus (primiary motor cortex or M1) |
| PT | planum temporale |
| PUT | putamen |
| RN | red nucleus |
| SCP | superior cerebellar peduncle |
| SFG | superior frontal gyrus |
| SII | secondary somatosensory cortex |
| SLF | superior longitudial fasciculus |
| SMA | supplementary motor area |
| SMG | supramarginal gyrus |
| SPL | superior parietal lobule |
| STG | superior temporal gyrus |
| STS | superior temporal sulcus |
| THA | thalamus |
| TP | temporal pole |
| TPG | temporoparietal junction |
| VER | Vermis |
